# Bulk RNA-Sequencing of small airway cell cultures from IPF and post-COVID lung fibrosis patients illustrates disease signatures and differential responses to TGF-*β*1 treatment

**DOI:** 10.1101/2023.03.01.530431

**Authors:** Katie Uhl, Shreya Paithankar, Dmitry Leshchiner, Tara E Jager, Mohamed Abdelgied, Kaylie Tripp, Angela M Peraino, Maximiliano Tamae Kakazu, Cameron Lawson, Dave W Chesla, Edward R. Murphy, Jeremy Prokop, Bin Chen, Reda E Girgis, Xiaopeng Li

## Abstract

IPF is a condition in which an injury to the lung leads to the accumulation of scar tissue. This fibrotic tissue reduces lung compliance and impairs gas exchange. Studies have shown that infection with COVID-19 significantly worsens the clinical outcomes of IPF patients. The exact etiology of IPF is unknown, but recent evidence suggests that the distal small airways, (those having a diameter less than 2 mm in adults), play a role in the early pathogenesis of IPF. TGF-β1 is a main driver of fibrosis in a variety of tissues; the binding of TGF-β1 to its receptor triggers a signaling cascade that results in inflammatory signaling, accumulation of collagen and other components of the extracellular matrix, and immune system activation. This study aimed to investigate possible mechanisms that contribute to worsening lung fibrosis in IPF patients after being diagnosed with COVID-19, with a particular focus on the role of TGF-β1. Small airway cell cultures derived from IPF and post-COVID-19 IPF patient transplant tissues were submitted for RNA-sequencing and differential gene expression analysis. The genetic signatures for each disease state were determined by comparing the differentially expressed genes present in the cells cultured under control conditions to cells cultured with TGF-β1. The genes shared between the culture conditions laid the framework for determining the genetic signatures of each disease. Our data found that genes associated with pulmonary fibrosis appeared to be more highly expressed in the post-COVID fibrosis samples, under both control and TGF-β1-treated conditions. A similar trend was noted for genes involved in the TGF-β1 signaling pathway; the post-COVID fibrosis cell cultures seemed to be more responsive to treatment with TGF-β1. Gene expression analysis, RT-PCR, and immunohistochemistry confirmed increased levels of BMP signaling in the IPF small airway cell cultures. These findings suggest that TGF-β1 signaling in IPF small airway cells could be inhibited by BMP signaling, leading to the differences in genetic signatures between IPF and post-COVID fibrosis.

## 1 Introduction

The etiology driving idiopathic pulmonary fibrosis (IPF), a disease characterized by the accumulation of scar tissue in the lungs, has so far remained undiscovered (1). It is estimated that approximately 50,000 people will be diagnosed with IPF annually and that 40,000 people will die from the disease in the same time frame (2). The fibrotic tissue begins to form after an injury to the lung and eventually inhibits the lung’s ability to exchange oxygen. Structural changes observed in IPF patients include widening of the airways (known as traction bronchiectasis), heterogenous reticulation, and the formation of honeycomb cysts (1). Recent evidence has been found to suggest that small airways, (those having a diameter of less than 2 mm), play an essential part in the pathogenesis of IPF(3–6). Still, our understanding of that role remains limited (1). One study found that patients suffering from IPF demonstrate a significant decrease in the number of terminal airways in the lung, even in minimally fibrotic regions (1).

The novel severe acute respiratory syndrome coronavirus 2 (SARS-CoV-2, also known as COVID-19) that emerged in 2019 quickly became associated with respiratory illness and injury amongst humans. Patients diagnosed with COVID-19 experience mild flu-like symptoms that can progress into a more severe disease manifestation characterized by alveolar damage, epithelial injury, and pulmonary fibrosis (7). A previous study performed on IPF patient lungs and fibrotic lesions from post-COVID-19 patient lungs found numerous mechanistic similarities, including an increase in extracellular matrix proteins, the presence of both fibrotic and inflammatory cytokines, and reduced lung function (8, 9).

Transforming growth factor beta (TGF-β1) is a pro-fibrotic cytokine whose expression is increased in human IPF lungs and bleomycin-induced fibrosis in animal models (6, 10, 11). The cellular sources for the upregulation of the three isoforms of TGF-β, (of which TGF-β1 is the most prevalent isoform found in pulmonary fibrosis), are the bronchial epithelium, myofibroblasts, alveolar macrophages, eosinophils, and hyperplastic type II alveolar epithelial cells (AECs) (10). The binding of TGF-β1 to its receptor leads to the recruitment of the SMAD2 and SMAD3 transcription factors. When these factors are phosphorylated, they form a complex with SMAD4 and move into the nucleus. This complex will interact with other transcription factors in a signaling cascade that eventually leads to the cellular characteristics of fibrosis, such as collagen deposition, fibroblast accumulation, immune system activation, and inflammatory signaling. The introduction of active TGF-β1 via adenovirus into rodent lungs was found to promote the development of severe pulmonary fibrosis (10).

The goal of this study was to use bulk RNA-sequencing of patient-derived small airway cell cultures to determine the genetic disease signatures of IPF and post-COVID fibrosis, with a focus on the role that TGF-β1 signaling plays in each disease. Considering the common structural characteristics between IPF and post-COVID fibrosis lungs, we compared the bulk RNA-sequencing data/transcriptome of small airway cells from both diseases with each other, as well as with normal small airway cell cultures. In addition, the effect of TGF-β1 treatment was compared among the three groups, and immunohistochemical staining was performed to provide possible explanations for the observed differences in responses to TGF-β1 between the two diseases. An understanding of the mechanisms behind the worsening of fibrosis in IPF patients after COVID-19 infection would provide insight into potential targets for future therapeutics.

## 2 Materials and Methods

### 2.1 Human lung explant samples

A total of 12 lung explant samples, (7 IPF patients and 5 post-COVID fibrosis patients), were obtained from patients undergoing lung transplantation at Corewell Health. A pathological assessment confirmed the diagnosis of fibrosis in the subjects used for this study. Normal control cultures were generated from 3 patients undergoing routine lobectomies for RNA-sequencing and RT-PCR; three of each disease lung explant samples were cultured for RNA-sequencing and RT-PCR as well (Supplementary Table 1). With the assistance of Lung Bioengineering Inc., 3 control lung tissue samples were collected from donors’ lungs that did not meet the criteria for transplantation. These samples were used as controls for immunohistochemical staining (Supplementary Table 1). For this study, “normal” refers to lung samples in which no disease was detected. “Control” refers to a small airway cell culture that has not been treated with TGF-β1.

### 2.2 Isolation and propagation of distal small airways

The small airways were micro-dissected from the patient’s lungs and digested in a buffer containing protease from *Streptomyces griseus*, (P5147, Millipore-Sigma), 1X Gentamicin, (15750060, Gibco), 1X penicillin/streptomycin, (15140122, Gibco), and DNase I (10104159001, Millipore-Sigma). Epithelial cells were isolated and expanded by culturing with PneumaCult™-Ex Plus Medium, (05040, STEMCELL Technologies), at 37°C, 5% CO_2_, for approximately 1 week of expansion. The cells were then cultured at the air-liquid interface (ALI), with or without TGF-β1, for 2-3 weeks before being collected for experimentation. As mentioned previously, TGF-β1 is a primary mediator of IPF, signaling a pro-fibrotic cascade when activated following lung injury or illness.

### 2.3 RNA isolation and sequencing

The sample tissue was first homogenized in TRIzol™ Reagent (15596026, Invitrogen) for RNA isolation and then treated with chloroform. The aqueous phase was added to an equal volume of 70% ethanol and transferred to a column from the RNeasy Mini Kit (74104, QIAGEN). The protocol for RNA extraction was carried out according to the manufacturer’s instructions. RNA concentration and quality measurements were taken using a NanoDrop™ OneC Microvolume UV-Vis Spectrophotometer (840274200, Thermo Scientific). Samples that met the RNA sequencing criteria were submitted to the Genomics Core at the Van Andel Institute for bulk RNA-sequencing with an Illumina NovaSeq 6000 instrument. The resulting data was uploaded to the National Center for Biotechnology Information (NCBI) database (GSE225549).

### 2.4 cDNA synthesis and Real-Time PCR

cDNA was synthesized from the isolated RNA samples using the SuperScript™ IV VILO™ Master Mix (11756050, Invitrogen) with ezDNase enzyme treatment according to the manufacturer’s protocol. cDNA concentration was quantified and diluted to 1 μg/μL. Individual master mixes for each gene of interest were prepared using the TB Green® Premix Ex Taq II (TLi RNaseH Plus) reagent kit (RR820A, Takara Bio) and primer sets ordered from Sigma-Aldrich (Supplementary Table 2). Each master mix contained a 1X concentration of TB Green® Premix Ex Taq II (TLi RNaseH Plus), 0.4μM of forward and reverse primers, and brought to a final volume with nuclease-free water. To each well of a 96-well PCR plate (AB-0600-L, Thermo Scientific), 18 μL of the prepared master mix was added, followed by 2 μL of template cDNA. The reaction was carried out using the CFX96™ Real-Time System (Bio-Rad) using the following settings: a holding stage of 30 seconds at 95°C, followed by 40 cycles of 95°C for 5 seconds and 60°C for 30 seconds, and ending with the melt curve stage. The instrument software from Bio-Rad was used to quantify the results.

### 2.5 Differential expression analysis

Transcript abundance of the RNA-sequencing reads was executed using Salmon (12). Differential expression analysis was performed by submitting the count data to DESeq2 and EdgeR (13, 14). For all downstream analyses, only genes with a p-value ≤ 0.01 and |log 2 Fold Change| ≥ 1 were considered to be significant. The Ingenuity Pathway Analysis tool from QIAGEN was used to identify pathways and upstream regulators represented by the list of differentially expressed genes (15). Gene ontology analysis was performed using the Gene Ontology resource (16, 17). Three DEG comparisons were carried out on the cultures: 1) the disease vs. normal lung under control conditions, 2) the disease vs. normal lung after TGF-β1 treatment, and 3) the disease control vs. the TGF-β1-treated cultures of the same disease (Figure 1C). Due to the large number of DEGs in each disease dataset, the disease signature was defined as the DEGs shared by both disease conditions (i.e., the overlap of Comparison #3). The *differences* between the IPF and COVID genetic signatures were determined by comparing the DEG results of IPF vs. COVID under normal conditions to the IPF vs. COVID DEGs under TGF-β1treatment conditions.

**Figure 1.**
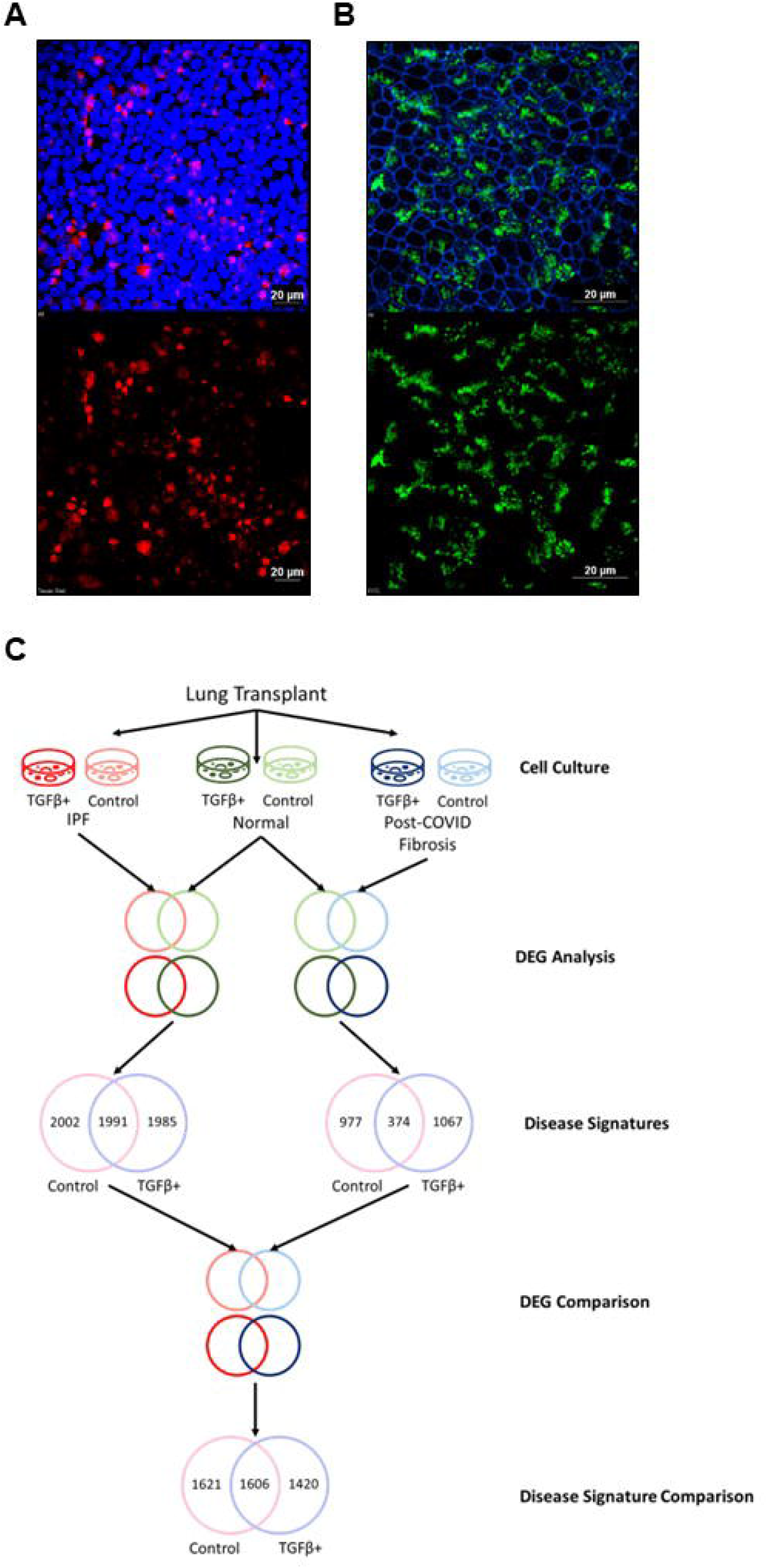
Culturing of small airway cells from patient lung tissue. **(A)** Expanded small airway cultures from IPF patient tissue express epithelial cell marker SCGB3A2 (red = SCGB3A2, blue = DAPI). The scale bar is equal to 20 μM. **(B)** IPF patient-derived small airway cultures display ciliated cells (green = cilia/alpha-tubulin, blue = actin). The scale bar is equal to 20 μM. **(C)** A flow chart displaying the sample processing and analysis aspects of this study. Small airway cells were isolated from lung explant tissue and cultured under control conditions or treated with TGF-β1. RNA-sequencing was carried out, and data were interpreted using the described dataset comparisons.

### 2.6 Immunohistochemistry

#### 2.6.1 Immunostaining of patient-derived cell cultures for epithelial cell markers

Small airway cell cultures were generated from lung explant tissue taken from IPF patients. After the cultures had been established, cells were fixed with 4% paraformaldehyde and mounted onto slides according to the protocol mentioned previously. Ciliated cells were detected by staining with mouse anti-acetylated α-tubulin (T7451, Sigma, St. Louis, MO) at a 1:1000 dilution. The epithelial cell marker SCGB3A2 was detected by staining the cell cultures with a 1:1000 dilution of rabbit anti-SCGB3A2 (ab181853, Abcam Inc., Cambridge, MA). The slides were washed with TBST and then incubated with the appropriate secondary antibody. After another series of washes with TBST and ddH_2_O, coverslips were mounted using ProLong™ Diamond Antifade Mountant with DAPI (P36962, Molecular Probes). Coverslips were sealed using nail polish and allowed to sit 24 hours before being imaged.

#### 2.6.2 Immunofluorescent detection of pSMAD1/5/8 in patient lung tissue

Small airway tissue was isolated from the patient’s lungs and fixed in 4% paraformaldehyde (4% PFA) at 4°C for at least 24 hours. The tissue was then incubated in 15% sucrose, 30% sucrose, and in a solution of 30% sucrose and 50% Scigen Tissue-Plus™ O.C.T. Compound (23-730-571, Fisher Scientific) for 24 hours each. After the sucrose gradient was complete, the tissue was embedded in O.C.T. and frozen on dry ice. Tissue slices with a thickness of 10 μM were placed onto Fisherbrand™ Tissue Path Superfrost™ Plus Gold Slides (15-188-48, Fisher Scientific). Slides were treated with 4% PFA for 20 minutes to fix the tissue, followed by washes with PBS (10010023, Gibco). Tissue permeabilization was carried out by incubating the slides with 0.1% Triton X-100 (X100, Sigma-Aldrich) for 5 minutes at room temperature. After washing again with PBS, sections were blocked with 5% BSA (A30075, Research Products International)/1% goat serum (G9023, Sigma-Aldrich) in PBS for two hours. The sections were incubated overnight with Rabbit anti-p-SMAD1/5/8 primary antibody (AB3848, Millipore-Sigma) in a solution of 2.5% BSA/0.5% goat serum at 4°C. Sections were washed with TBST and then incubated with 1:1000 secondary antibody (F(ab’)2-Goat anti-Rabbit IgG (H+L) Secondary Antibody, Alexa Fluor™ 568, A-21069) in 2.5% BSA/0.5% goat serum in PBS for 1 hour at room temperature. A series of washes with TBST and dd_2_O was carried out, and then coverslips were mounted to the slides using ProLong™ Diamond Antifade Mountant with DAPI (P36962, Molecular Probes). The coverslips were sealed with nail polish, and the slides were imaged using confocal microscopy. Signal quantification was carried out using analysis software available from Nikon and GraphPad 9.4.1 was used to perform ordinary one-way ANOVA between groups.

## 3 Results

### 3.1 IPF patient-derived small airway cell cultures maintain cell markers and characteristics found in fibrotic lung tissues

#### 3.1.1 IPF patient-derived small airway cell cultures display epithelial and ciliated cell markers

Small airway cells taken from IPF patient lungs were expanded into cell cultures. These cultures were stained for epithelial and ciliated cell markers. The results of the staining demonstrated that the small airway cell cultures express the small airways epithelial cell marker SCGB3A2 (Figure 1A) and are also positive for ciliated cells (Figure 1B).

#### 3.1.2 IPF patient-derived cell cultures demonstrate genetic characteristics of IPF whole lung and small airway culture RNA-sequencing

The sequencing results for the IPF small airway cultures from this study were compared to previously published RNA-sequencing data from a mouse PF model mimic human IPF pathology (6). A total of 101 genes were found to be in common between the two groups. These genes were submitted to string.db (https://string-db.org/) to discover the key fibrosis genes based on known and predicted protein interactions (18). A heatmap of the 17 genes identified by string.db as having the most significant protein interactions was generated based on the IPF patient-derived small airway cell cultures (Figure 2A). This list included AURKA, BUB1B, CDC20, FOXM1, and others; all of the selected genes were found to have increased expression levels in the IPF patient samples versus the normal lung samples. RT-PCR was used to confirm the significant increase in FOXM1 expression in the IPF small airway cell culture samples (Figure 2B).

**Figure 2.**
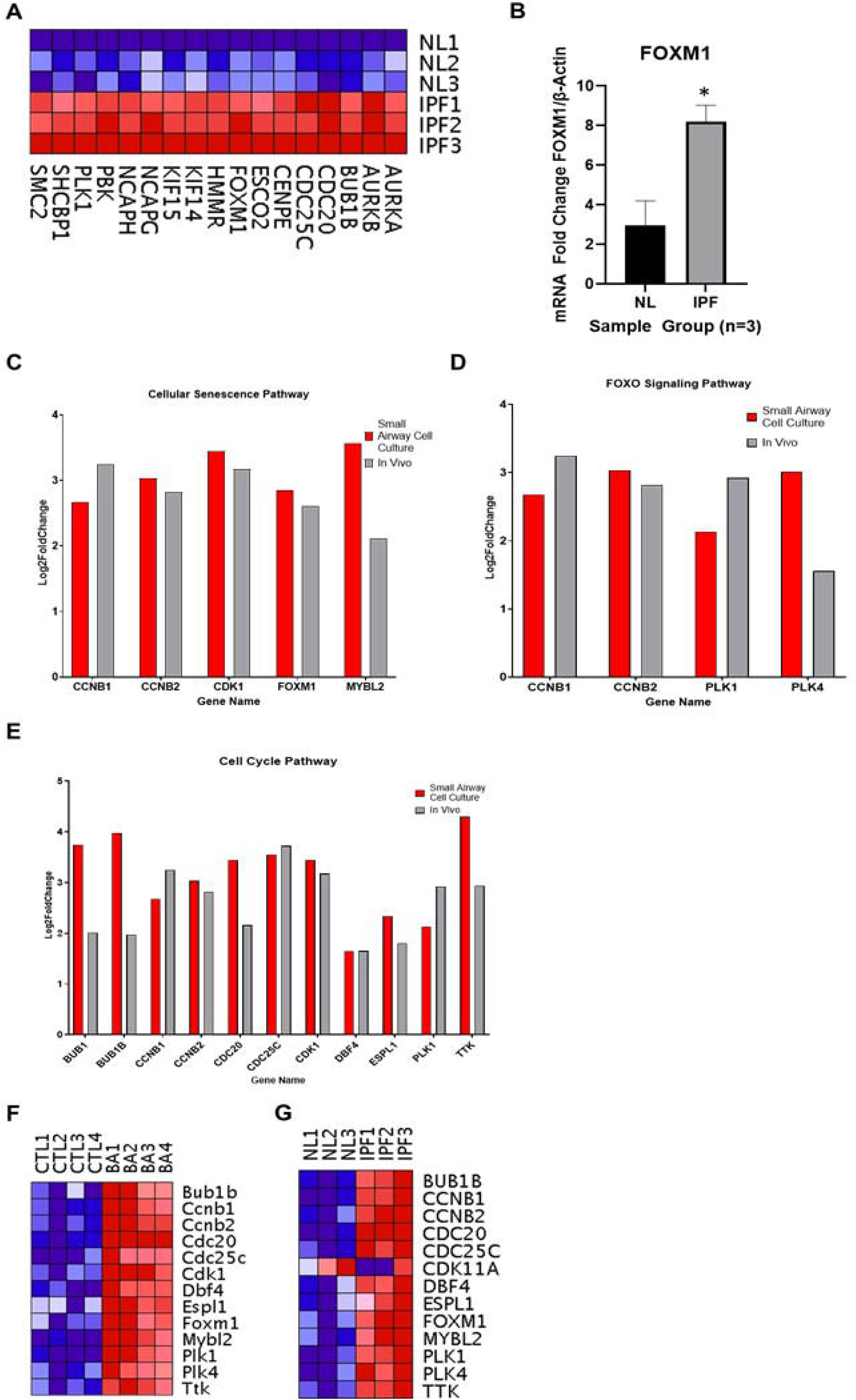
IPF patient-derived small airway cell cultures maintain cell markers and characteristics found in fibrotic lung tissues. **(A)** Heatmap displaying genes identified by string.db to be important in the pathogenesis of IPF. **(B)** RT-PCR results of FOXM1 expression in normal lung vs. IPF small airway cell cultures (* indicates p-value < 0.05). **(C-E)** DEG lists were compared between human small IPF small airway cell culture and bleomycin-ATP12A fibrosis mouse model. The expression of genes involved in the cellular senescence pathway **(C)**, the cell cycle pathway **(D)**, and the FOXO signaling pathway **(E)** are shown. The expression of these selected genes compared to their relevant controls is shown in **(F)** for the mouse samples and in (**G)** for the IPF small airway cell cultures.

#### 3.1.3 Small airway cell cultures from IPF patients reflect expression levels of select genes in an in vivo model of bleomycin-induced fibrosis

Previously published RNA-sequencing data was used to compare genetic signatures between IPF patient small airway cell cultures and a mouse PF model mimic human IPF pathology (6). In addition to treatment with bleomycin, the same mice were also treated with a viral vector that increased the expression of the non-gastric H+/K+-ATPase ATP12A. A research study performed by this group demonstrated that ATP12A is overexpressed in the small airways of IPF patients and overexpression of ATP12A in bleomycin-treated mice exacerbated pulmonary fibrosis (6). When compared, the two datasets had 101 genes in common (with one another (Supplementary Table 3). KEGG pathway analysis was performed using the Network Analyst online tool (https://www.networkanalyst.ca/) (19). The main pathways identified included the cellular senescence pathway, the cell cycle pathway, and the FOXO signaling pathway (Figure 2C-E). The expression levels of the genes corresponding to these pathways were visualized for the individual datasets using a heatmap (Figure 2F-G). Overall, the small airway cells generated from IPF patient tissue reflect the expression trend seen in the bleomycin/ATP12A-treated mice.

### 3.2 Determination of IPF genetic signature

#### 3.2.1 RNA-sequencing of IPF and normal cultured small airway cells demonstrate unique disease signatures

Small airway cells cultured from IPF patients were compared to “normal” small airway cells from patients not known to have airway disease. These cells were either treated with TGF-β1 or kept as control cultures. Under both culture conditions (1. IPF small airways control (no TGF-β1) and normal lung small airway control (no TGF-β1) and 2. IPF small airways control (TGF-β1+) and normal lung small airway control (TGF-β1+)), there were nearly 4000 DEGs out of which 1991 were found common between both signatures. (Figure 3A).

**Figure 3.**
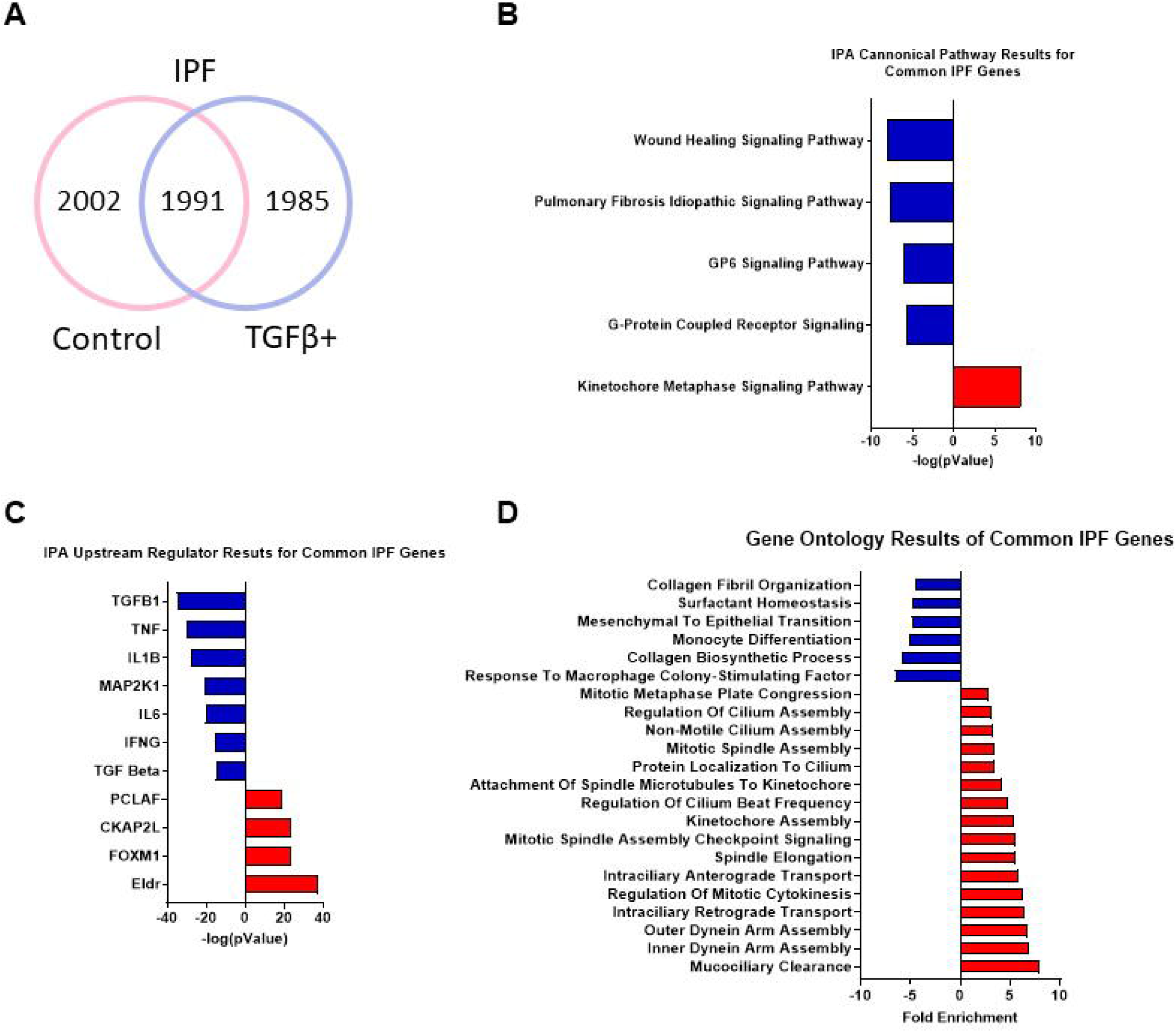
Analysis of the genetic disease signature for small airway cells cultured from IPF patients. **(A)** Venn diagram showing common DEGs found by comparing small airways between IPF lungs and normal lungs in both TGF-β1 treated and untreated culture conditions. **(B)** IPA canonical pathway results for the common IPF cell culture genes. **(C)** IPA upstream regulator results for common IPF cell culture genes. **(D)** Gene ontology results of the common genes in the IPF small airway cell cultures.

#### 3.2.2 Canonical pathway analysis of IPF vs. normal small airway cultures shows inhibition of IPF-related pathways

The list of common DEGs for the IPF vs. normal small airway culture comparison was submitted as input to the Ingenuity Pathway Analysis (IPA) tool available from QIAGEN. Genes, and their associated fold changes, are categorized based on QIAGEN’s database of pathways and regulators. The top pathway analysis results for the IPF vs. normal shared DEGs included the kinetochore metaphase signaling pathway, the wound healing pathway, and the GP6 signaling pathway (Figure 3B) (Supplementary Table 4). Interestingly, IPA predicts these pathways to be inhibited, with the expception of the kinetochore metaphase signaling pathway.

#### 3.2.3 Upstream regulators associated with common IPF genes predict the inhibition of the TGF-β1-signaling pathway and host immune response

Data from the common DEG list was used to identify upstream regulators and predict their activation states (Figure 3C) (Supplementary Table 5). Among the top regulators expected to be activated are molecules involved in the cell cycle and mediation of DNA damage, such as ELDR, FOXM1, and CKAP2L. Genes related to the host immune response and inflammation signaling, (TNF, IL-B, and IL-6), are predicted to be inhibited. Although the experimental cultures were treated with TGF-β1 before sequencing, TGF-β1 is the most significant inhibited pathway. It is worth noting that TGF-β family members TGF-β2 and TGF-β3 are also predicted to be inhibited, as represented by the TGF beta entry.

#### 3.2.4 Gene ontology results for common IPF genes show activation of ciliary movement and cell cycle regulation and a downregulation of genes associated with ECM components and immune response

The top 25 gene ontology results for the upregulated DEGs associated with the IPF vs. normal culture comparison demonstrated a trend towards ontologies relating to either the organization of cilia or regulation of the cell cycle (Figure 3D) (Supplementary Table 6). The gene ontology with the most significant fold enrichment was mucociliary clearance, followed by inner and outer dynein arm assembly, and intraciliary retrograde transport, all of which are associated with ciliary movement. Ontologies corresponding to the cell cycle include kinetochore assembly, regulation of mitotic cytokinesis, and mitotic spindle assembly. The associated gene ontologies for the downregulated DEGs were more varied than the upregulated DEG results (Supplementary Table 7). The synthesis and organization of collagen fibrils are represented by the entries of the collagen biosynthetic process and collagen fibril organization. In addition, the presence of the immune-related entries monocyte differentiation and response to macrophage colony-stimulating factor reflects the trend seen in the inhibited upstream regulator results.

### 3.3 Determination of post-COVID fibrosis genetic signature

#### 3.3.1 Gene expression differences are observed between post-COVID fibrosis cells and normal small airway cells cultured from human patients

The sequencing results from the post-COVID fibrosis small airway cells were compared to the data from normal small airway cell cultures. The comparison of post-COVID fibrosis cell cultures and the normal cell cultures, without TGF-β1 treatment, demonstrated a total of 977 DEGs. The TGF-β1-treated cultures resulted in 1067 DEGs between the two groups. A comparison of the DEG list from both treatment conditions shows 374 DEGs in common (Figure 4A).

**Figure 4.**
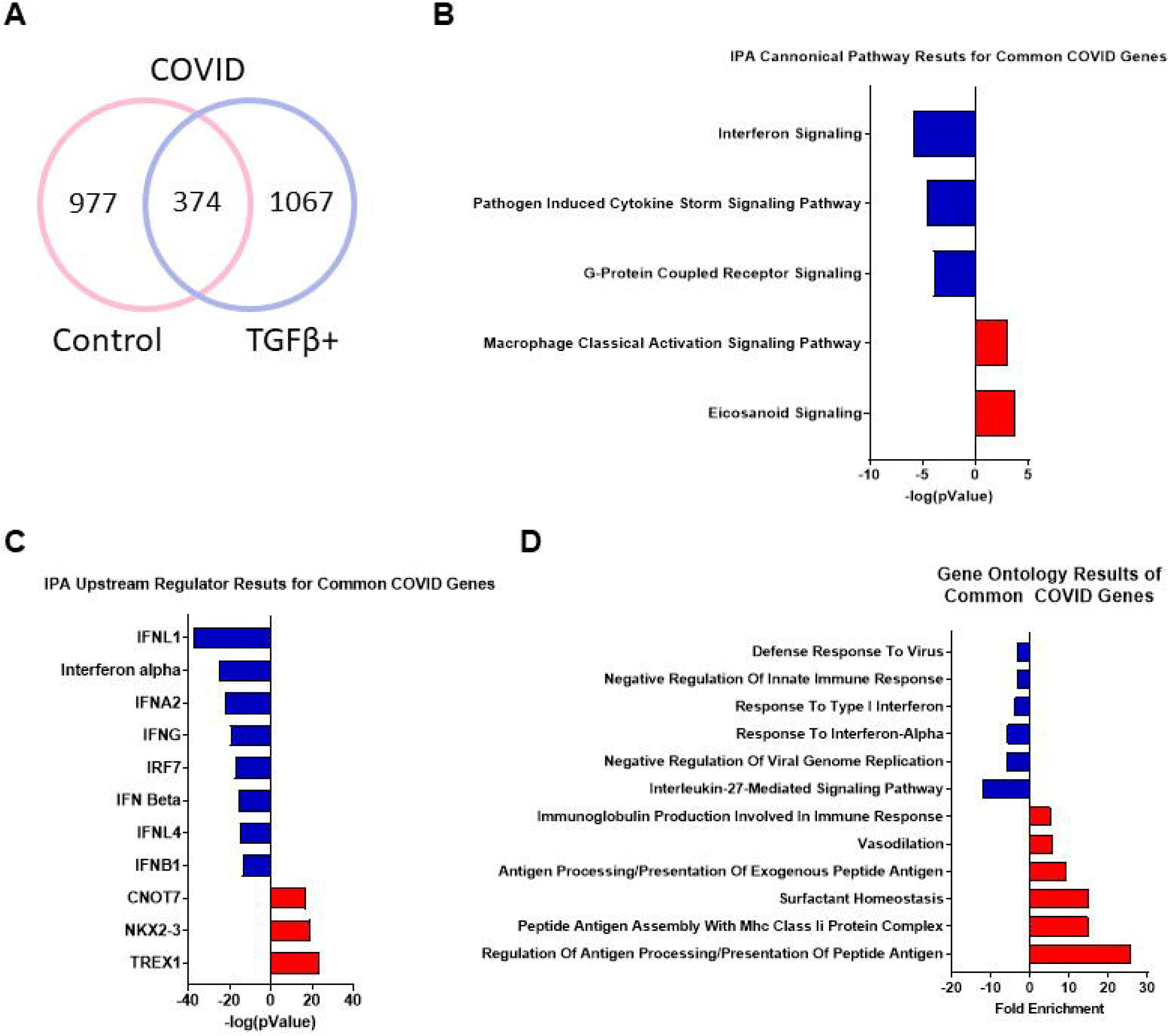
Analysis of the genetic disease signature for small airway cells cultured from post-COVID fibrosis patients. **(A)** Venn diagram showing common DEGs found by comparing small airways between IPF lungs and normal lungs in both TGF-β1 treated and untreated culture conditions. **(B)** IPA canonical pathway results for the common post-COVID fibrosis cell culture genes. **(C)** IPA upstream regulator results for common post-COVID fibrosis cell culture genes. **(D)** Gene ontology results of the common genes in the post-COVID fibrosis small airway cell cultures.

#### 3.3.2 Ingenuity Pathway Analysis of COVID vs. normal small airway cultures demonstrates a trend toward host immune response and interferon signaling

The list of 374 common DEGs between the control and TGF-β1-treated cultures was submitted for pathway analysis using the Ingenuity Pathway Analysis (IPA) tool from QIAGEN (Figure 4B) (Supplementary Table 8). Each pathway is assigned an activation state and a significance value based on the fold changes of the input genes. Included in the most significant inhibited pathways were interferon signaling and pathogen-induced cytokine storm signaling pathway. Activated pathways included eicosanoid signaling and the macrophage classical activation signaling pathway, both of which are related to the innate immune response.

#### 3.3.3 Upstream regulator results for post-COVID fibrosis common genes show predicted activation of the immune response, but not cytokine-related molecules

Another function within the IPA is the identification of significant upstream regulators, and their predicted activation states, present in the DEG list. The same list of common post-COVID fibrosis DEGs was used for upstream regulator analysis (Figure 4C) (Supplementary Table 9). Upstream regulators involved in various components of the innate immune response were predicted to be activated, such as CNOT7, NKX2-3, and TREX1. Interestingly, upstream regulators related to the role of cytokines in the immune response, (IFNL1, interferon alpha, IRF7, and IFN beta), were all predicted to be inhibited. This finding reflects the inhibition of the pathogen-induced cytokine storm signaling pathway seen in the pathway analysis of the common post-COVID fibrosis genes.

#### 3.3.4 Gene ontology analysis of common upregulated post-COVID fibrosis genes indicates activation of the immune response, and a decrease in ontologies related to cilium movement and DNA repair

The gene ontologies for the common upregulated post-COVID fibrosis genes reveal a slight trend towards the adaptive immune system, with two entries related to antigen processing and presentation (Figure 4D) (Supplementary Table 10). Also present is immunoglobulin production, another part of the body’s overall immune response. Surfactant homeostasis and vasodilation are other ontologies of note. The gene ontologies for the downregulated post-COVID fibrosis genes include several entries related to DNA repair and replication, such as double-strand break repair via break-induced replication, DNA replication initiation, and regulation of DNA-templated DNA replication (Supplementary Table 11). Downregulated gene ontologies demonstrate a predicted inhibitory effect of cilium movement with outer dynein arm assembly, epithelial cilium movement involved in extracellular fluid movement, sperm flagellum assembly, and regulation of the microtubule-based process. Entries related to the immune system are present in the downregulated gene ontologies, including responses to various interferons.

### 3.4 Comparison of IPF and COVID genetic signatures

#### 3.4.1 IPF vs. COVID genetic signature comparison reveals the predicted activation of cell cycle regulators and inhibition of fibrosis-related pathways in the IPF cell cultures

In order to highlight the transcriptomic differences between the IPF and post-COVID fibrosis cell cultures, the results from the IPF vs. COVID differential gene expression analyes were compiled. Between the two culture conditions, (Control and TGF-β1), there were a total of 1606 genes that were differentially expressed between IPF and COVID (Figure 5A) (Supplementary Table 12). This combined gene list represents the DEGs observed when the IPF cell cultures were compared to the post-COVID fibrosis cell cultures. Amongst the top canonical pathways generated by IPA for the shared DEGs are cyclins and cell cycle regulation, and mitotic roles of polo-like kinase (Figure 5B). Two pathways directly related to fibrosis, the pulmonary fibrosis idiopathic signaling pathway and the wound healing pathway, are present but are predicted to be inhibited in the IPF cell culture.

**Figure 5.**
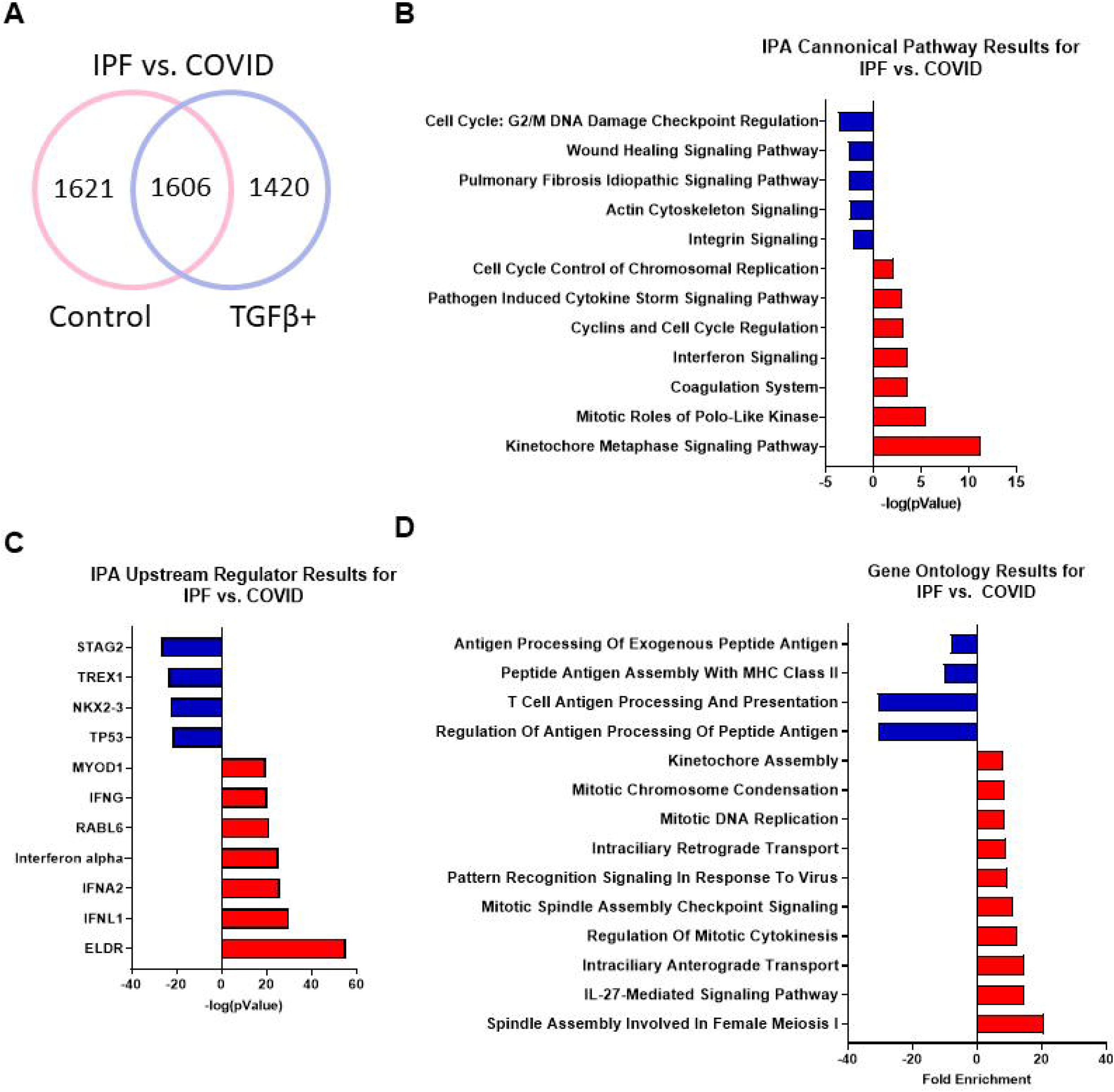
Comparison of the genetic disease signatures for small airway cells cultured from IPF and post-COVID fibrosis patients. IPF and post-COVID fibrosis RNA-sequencing data was compared under control and TGF-β1-treated conditions. **(A)** A diagram visualizing the number of common genes between the control and TGF-β1-treated cultures. **(B)** IPA canonical pathway results for the genes shared between the IPF and post-COVID fibrosis data. **(C)** IPA upstream regulator results for common DEGs between the disease groups. **(D)** Gene ontology results of the upregulated DEGs found in both disease groups. **(E)** Gene ontology results of the common downregulated genes in the IPF and post-COVID fibrosis samples.

#### 3.4.2 Upstream regulator analysis of IPF vs. COVID DEGs show activation of host immune response and regulation of the cell cycle

Like the IPA analysis of the IPF and post-COVID fibrosis datasets, activated upstream regulators for the DEGs resulting from the IPF vs. COVID comparison are predicted to include proteins involved in host immune response; IFNG, interferon alpha, IFNA2, and IFNL1 are predicted to be activated in the IPF cell culture (Figure 5C) (Supplementary Table 13). Also in the IPF small airway cell culture results are proteins involved in the cell cycle and differentiation; MYOD1 and RABL6 are predicted to be activated, while TP53 and STAG2 are predicted to be inhibited.

#### 3.4.3 Gene ontologies for IPF vs. post-COVID fibrosis DEGs demonstrate upregulation of cell growth and intraciliary transport ontologies and downregulation of antigen processing

Gene ontologies for the DEGs between IPF and post-COVID fibrosis, under both control and TGF-β1 treatment conditions, include a wide variety of cell growth, division, and meiosis-related ontologies: spindle assembly involved in female meiosis I, regulation of mitotic cytokinesis, and mitotic spindle assembly checkpoint are all present (Figure 5D) (Supplementary Table 14). Intraciliary transport, both anterograde and retrograde, was also among the top gene ontology results for the IPF small airway cell cultures. Several antigen processing and presentation ontologies can be found in the results for the differentially expressed downregulated genes (Supplementary Table 15).

#### 3.4.4 Comparison of fibrotic signatures in the IPF and post-COVID fibrosis small airway cell cultures suggest fibrosis-related genes are more highly expressed in post-COVID fibrosis samples

The read count data from both the IPF and post-COVID fibrosis datasets were used to generate a heatmap comparing expression levels of genes related to pulmonary fibrosis (Figure 6A) (20). Overall, the untreated post-COVID fibrosis samples have higher expression of the selected pulmonary fibrosis-related genes than the IPF samples. The addition of TGF-β1, a known driver of fibrosis, did increase the expression of the selected genes, although the post-COVID fibrosis samples did appear to show molecular indicators of worsening fibrosis. A comparison of selected IPA canonical pathways highlights the differences between the IPF and post-COVID fibrosis small airway cultures (Figure 6B). The wound healing signaling pathway, the pulmonary healing pathway, and the pulmonary fibrosis signaling pathway are predicted to be activated in the post-COVID fibrosis samples, but not in the IPF samples.

**Figure 6.**
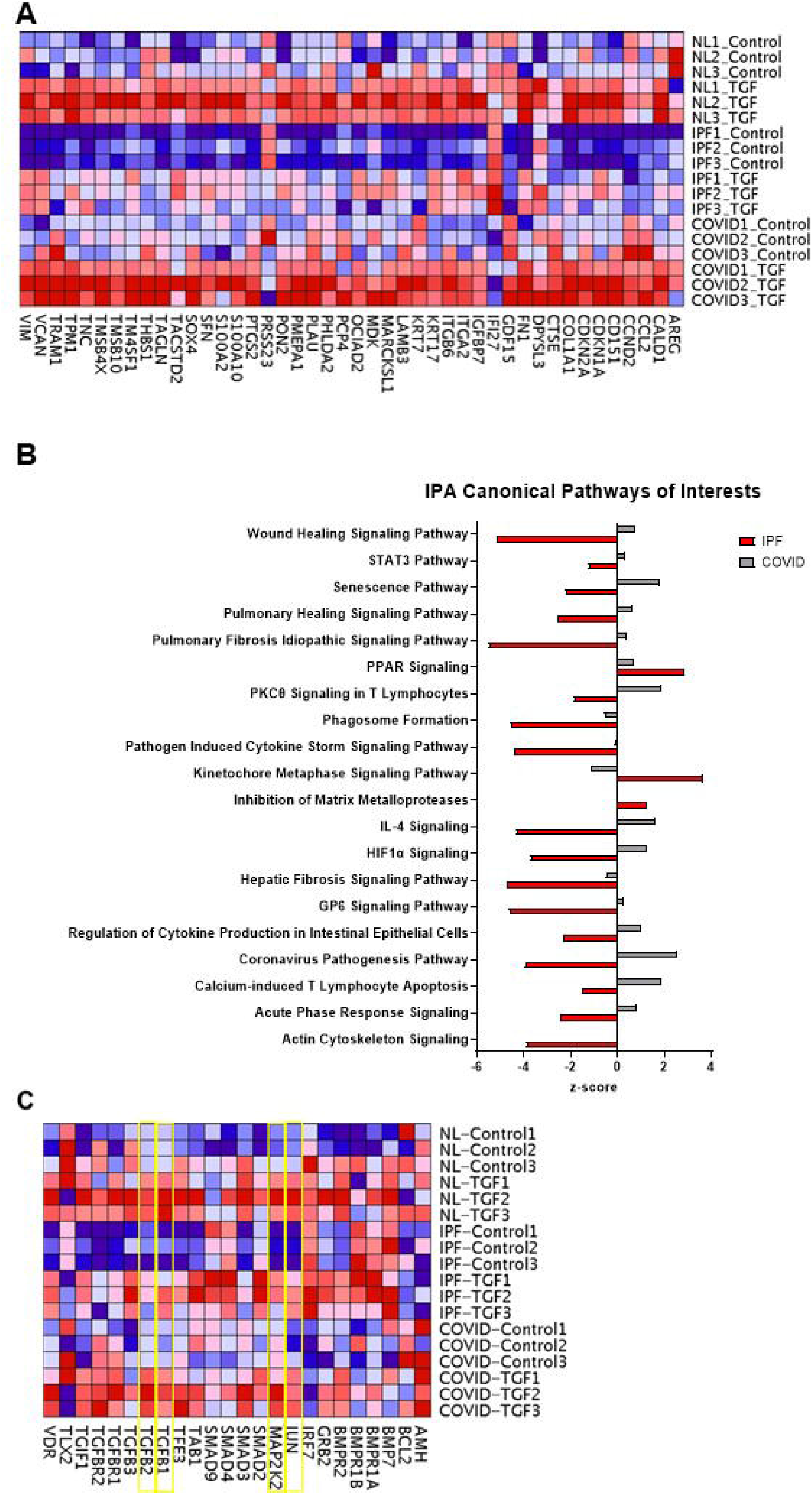
Fibrotic signatures in IPF and post-COVID fibrosis small airway cultures. **(A)** Heatmap displaying gene expression levels for signature genes involved in the development of IPF (NL = normal lung, IPF = idiopathic pulmonary fibrosis, COVID = post-COVID fibrosis). **(B)** Predicted activation states of fibrosis-related pathways in the IPF and post-COVID fibrosis small airway cultures; a negative z-score indicates that the pathway is predicted to be inhibited by the IPA software. **(C)** Heatmap displaying gene expression levels for genes involved in the TGF-β1 signaling pathway, a key driver of fibrosis. Genes of interest have been highlighted in yellow.

#### 3.4.5 Gene expression comparisons between IPF and post-COVID fibrosis small airway cell cultures identify differences in responses to TGF-β1 treatment

Genes from the TGF-β1signaling pathway were selected for gene expression analysis (Figure 6C). The read count datasets from both IPF and post-COVID fibrosis samples were visualized using a heatmap based on TGF-β1 signaling pathway genes. In normal lung samples, most genes associated with the TGF-β1 signaling pathway are expressed at low levels until the culture is treated with TGF-β1; examples of these molecules include JUN, MAP2K2, TGF-β1, and TGF-β2 (Figure 6D-G). For the IPF lung samples, the expression levels of many of the genes were very low, even compared to the normal lung samples. TGF-β1 treatment did increase expression levels, but those values were still low compared to the post-COVID fibrosis samples. The baseline expression of TGF-β1 signaling pathway-related genes is moderately low in the untreated post-COVID fibrosis samples, though levels are still higher than the IPF sample baseline. Treatment of the post-COVID fibrosis cultures with TGF-β1 results in an increase in the expression of selected genes.

#### 3.4.6 Differences in BMP signaling are observed between post-COVID fibrosis and IPF cell cultures

The possible effect of BMP signaling on the cell culture responses to TGF-β1 was explored. The BMP signaling pathway is a known inhibitor of the TGF-β1 signaling pathway (Figure 7A) (21). Expression levels of genes involved in the BMP signaling pathway were visualized across the samples with a heatmap (Figure 7B). Analysis of these genes shows higher expression in the IPF samples than in the post-COVID fibrosis samples; BMP7, BMPR1A, BMPR1B, SMAD1, and SMAD2 all demonstrate this expression pattern. RT-PCR was used to validate this observation (Figures 7C, 7D, and 7E). The fold changes for BMP7 and BMPR1B are significantly higher in the IPF samples when compared to the normal lung and post-COVID fibrosis samples. There are also differences in fold changes for BMPR1A, although only the comparison of IPF to normal lungs is considered significant.

**Figure 7.**
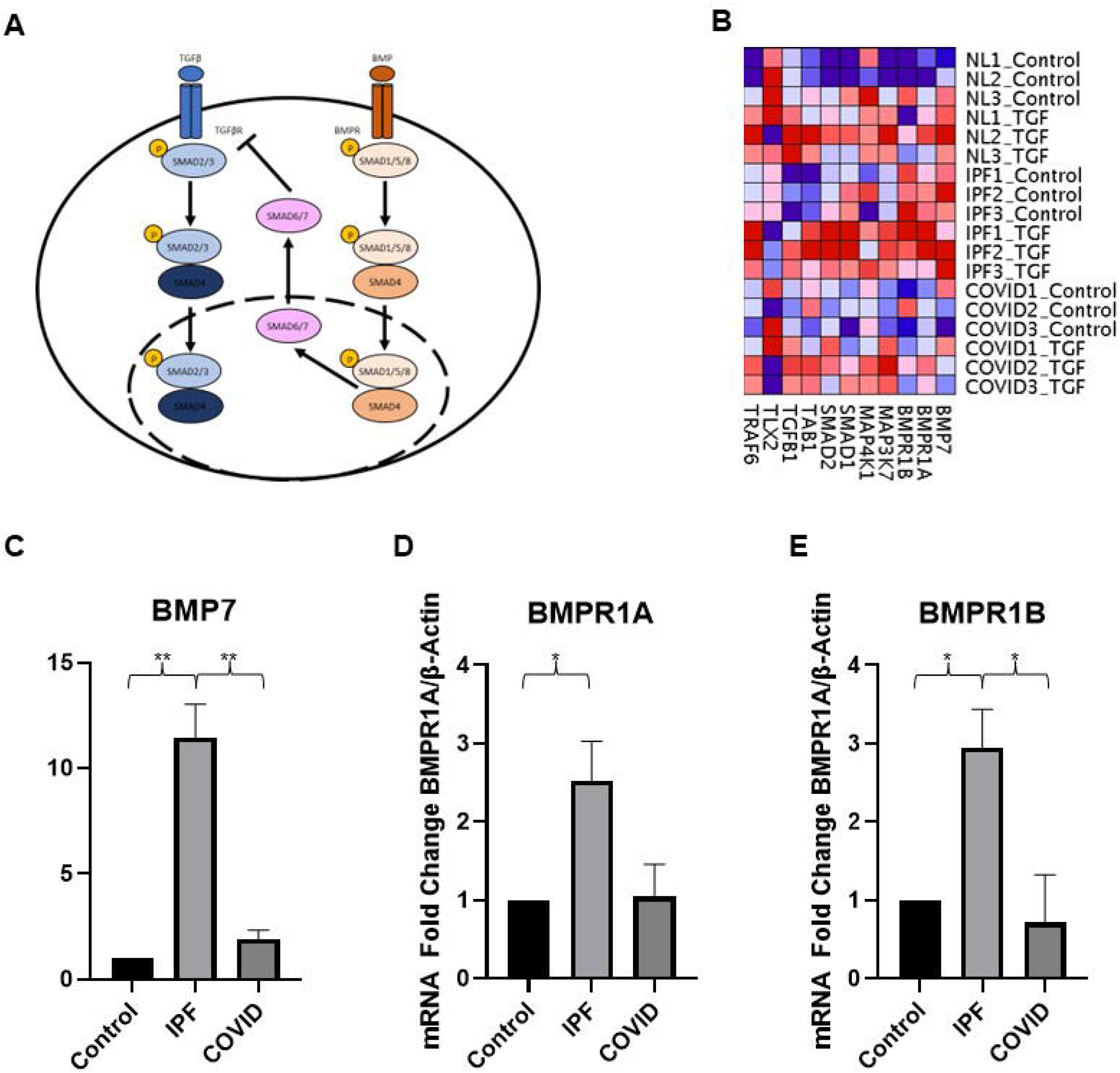
Evaluation of BMP signaling in patient-derived small airway cell cultures. **(A)** Mechanism of inhibition of TGFB signaling by BMP. **(B)** Heatmap displaying the expression levels of genes involved in the BMP signaling pathway (NL = normal lung, IPF = idiopathic pulmonary fibrosis, COVID = post-COVID fibrosis). **(C-E)** RT-PCR quantification of fold changes for genes associated with the BMP signaling pathway (** indicates p-value < 0.001, * indicates p-value < 0.05).

#### 3.4.7 pSMAD1/5/8 expression and nuclear localization in normal, IPF, and post-COVID fibrosis tissues

The expression of pSMAD1/5/8, a regulator of BMP signaling, was visualized in the normal, IPF, and post-COVID fibrosis tissues using immunofluorescence (Figure 8A). Amongst the three tissue types, the IPF samples demonstrated higher levels of pSMAD1/5/8 nuclear localization (Figure 8B). The levels of nuclear localization are the lowest in the post-COVID fibrosis tissue, followed by normal lung tissue. The comparison of the normal lung samples to post-COVID fibrosis samples and IPF samples to post-COVID fibrosis tissue have significant p-values.

**Figure 8.**
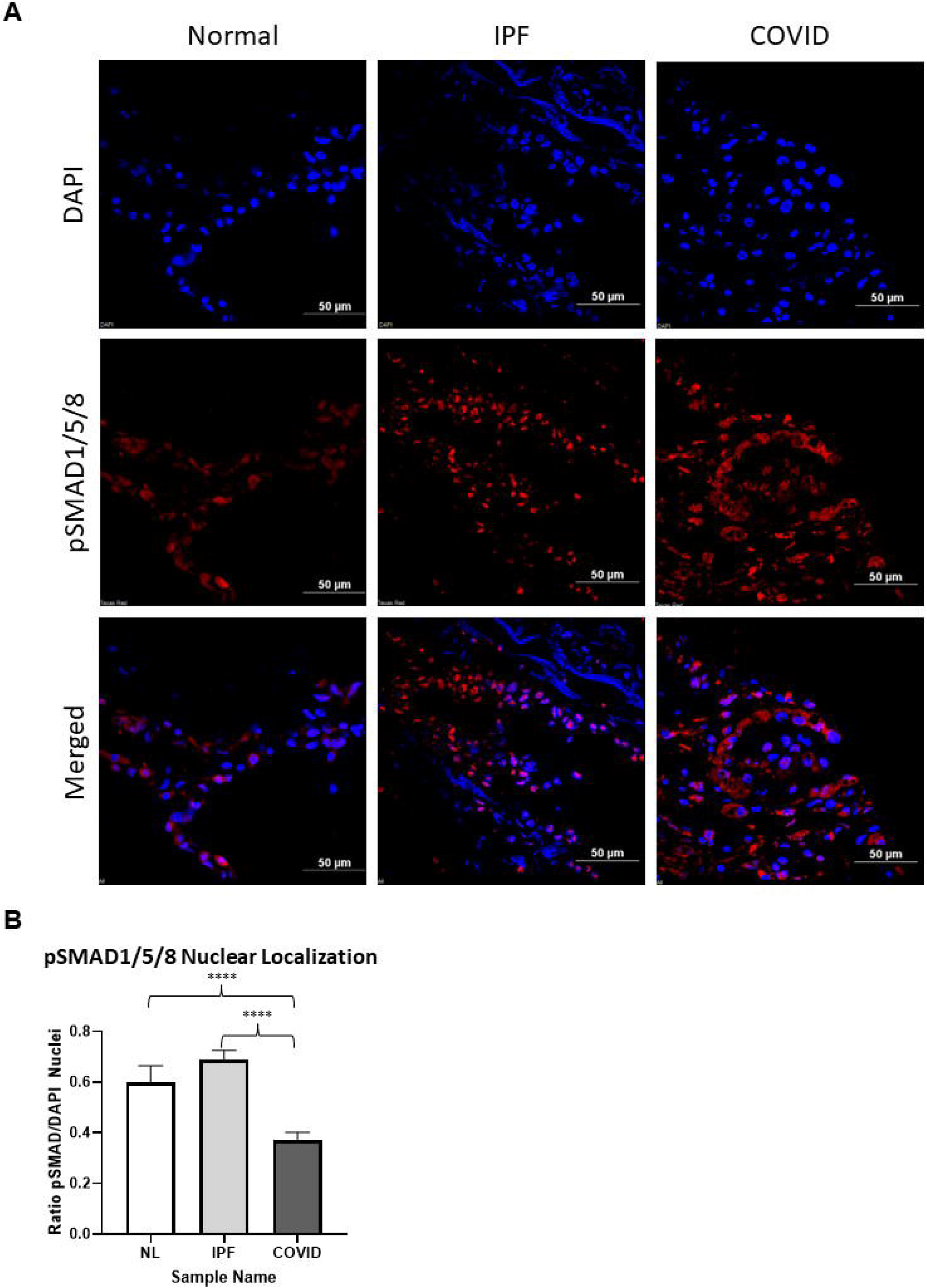
Localization of pSMAD1/5/8 to the nucleus in patient sample tissue. **(A)** IHC images of patient samples, (Normal = healthy donor lungs, IPF = idiopathic pulmonary fibrosis, COVID = post-COVID fibrosis), stained for pSMAD1/5/8 (red) and DAPI (blue). The scale bar is equal to 50 μM. **(B)** Quantification of nuclear localization of pSMAD1/5/8 across the different tissue types (**** indicates p-value < 0.0001).

## 4 Discussion

Idiopathic pulmonary fibrosis (IPF) is a devasting disease that impairs gas exchange. It is the most common idiopathic interstitial pneumonia (IIP) and accounts for 20-50% of chronic interstitial lung disease cases (22). Some of the hallmarks of IPF include the destruction of the air sacs in the lung (alveoli), the accumulation of scar tissue, and abnormal morphology changes in the airways (23). TGFB1 has been shown to play an important role in the pathogenesis of IPF, influencing disease characteristics such as myofibroblast accumulation and collagen deposition (10). The causes of IPF are not fully understood, but there is research to support that environmental factors, cigarette smoking, and certain infections may increase the risk of developing this disease (22).

Pulmonary fibrosis is a significant risk factor in developing complications following infection by COVID-19 (24). Research has shown that the mechanism behind acute respiratory distress syndrome (ARDS) is similar to pathways underlying the development of lung fibrosis (24). Infection by COVID-19 stimulates the immune system and results in systemic hyperinflammation and a massive release of cytokines known as a cytokine storm; IPF patients without COVID-19 also experience the activation of inflammatory responses and overall stimulation of the immune system (8). Other pathological characteristics shared between COVID-19 and pulmonary fibrosis patients include the accumulation of fibrin in the alveolar spaces, (acute fibrinous and organizing pneumonia), the deterioration of vascular systems, and the buildup of collagen and other extracellular matrix components (8). This study sought to further explore the differences and similarities between small airway cell cultures derived from IPF and post-COVID fibrosis patient lung tissue, with a focus on TGF-β1, a main driver of tissue fibrosis.

### 4.1 Patient-derived small airway cell cultures maintain tissue-specific markers and reflect gene expression patterns observed in fibrotic lung tissue

This study made use of small airway cultures generated from IPF and post-COVID fibrosis lung tissue. Cells from the expanded IPF cultures were stained and found to be positive for secretoglobin family 3A member 2 (SCGB3A2), a small airway epithelial cell marker (Figure 1B). Combined with the positive staining for ciliated cells (Figure 1C), this data suggests that the patient-derived cell cultures maintain cell-specific markers found in their tissue of origin, and thus can be used as an *in vitro* model to study genetic disease signatures.

Currently, there are very few studies that explore the role of small airways in IPF. A study published in 2021 by Stancil et al. found that cell cultures from IPF distal airways did demonstrate biophysical characteristics of pulmonary fibrosis disease progression (23). However, the authors focused more on the ERBB-YAP signaling pathway and its effect on the jamming transition of the IPF epithelial (23). This makes the RNA-sequencing results from this and from Stancil et al. difficult to compare, as our experiments were not performed with the intention of modifying the jammed transition state of the epithelial cells. Instead, our group elected to use a previously published in vivo mouse model of bleomycn-induced lung fibrosis as a comparison to determine if our IPF-patient derived cell cultures demonstrated key IPF tissue and cell characteristics.

An in vivo model of lung fibrosis was generated by our group (6). Mice were treated with a combination of bleomycin and an adenovirus that mediated the expression of the non-gastric ATPase known as ATP12A. This mouse model was used for comparison because the overexpression of ATP12A more closely reflects the increase in ATP12A expression seen in human IPF lungs (6). RNA-sequencing data from these mice were compared to data from small airway cell cultures derived from IPF patients. These two datasets had a total of 101 DEGs in common. This gene list was submitted to string.db for protein interaction analysis (18). This analysis identified 17 genes as being important to the development of IPF: AURKA, AURKB, BUB1B, CDC20, CDC25C, CENPE, ESCO2, FOXM1, HMMR, KIF14, KIF15, NCAPG, NCAPH, PBK, PLK1, SHCBP1, and SMC2. The expression levels of these genes was visualized on a heatmap comparing normal lung small airway cell cutlures to IPF patient-derived small airway cell cultures (Figure 2A). The expression of each gene was found to be increased in the IPF small airway cell cultures. A study published in 2019 found that AURKB expression was increased in IPF patient mesenchymal cells and may play a role in apoptosis resistance and in the control of increased cell proliferation in IPF lung fibroblasts (25). FOXM1 is a transcription factor that has been shown to signal the activation of the epithelial-to-mesenchymal transition in alveolar type II epithelial cells. Signaling by this protein stimulates the development of pulmonary fibrosis by inducing the differentiation and proliferation of activated fibroblasts (26). The increased expression of FOXM1 in the IPF small airway cultures was found to be significant when compared to cell cultures generated from normal small airway tissue (Figure 2B). Numerous studies have linked variants in the KIF15 gene to an increased risk of developing IPF (27–29). Significant genes present in our dataset reflect the characteristics seen in IPF lung tissue and IPF cell cultures.

The 101 DEGs shared between the human IPF cell cultures and the aforementioned bleomycin-induced lung fibrosis mouse model are involved with KEGG pathways known to be associated with fibrosis: the cellular senescence pathway, the cell cycle pathway, and the FOXO signaling pathway. The fold change of the individual genes for each pathway was calculated and compared between the datasets (Figure 2C-E). Cellular senescence has been shown to mediate the characteristic effects of pulmonary fibrosis; biomarkers of senescence, such as CDKN2A and TP53, are highly expressed in IPF patient lungs (11). The treatment of mice with bleomycin induces a type of lung injury whose genetic signature is similar to that seen in human IPF patients; this signature includes genes involved in senescence (11). Our analysis of genes involved in the cellular senescence pathway demonstrates similar expression patterns between the in vivo mouse model, (mice treated with the bleomycin and the ATP12A adenovirus), and small airway cultures derived from IPF patient lung tissue. A similar trend is observed in the genes involved in the cell cycle pathway and the FOXO signaling pathway (Figure 2D-E). Transcription factors involved in the Forkhead family are implicated in the development of lung fibrosis and are being explored as targets for future IPF therapies (30, 31). Likewise, genes involved in the regulation of cell cycle progression have been associated with the pathogenesis of IPF (32). The similarities between the commonly used bleomycin-induced fibrosis model and the IPF patient-derived small airway cultures can also be observed when the expression levels of the genes of interest are visualized using a heatmap (Figure 2F-G). Expression of the selected genes appears to be lower in each model’s respective control samples, (labeled as “CTL” for the bleomycin and ATP12A-treated mice, and “NL” for the patient cell cultures) than in the experimental samples. The parallels observed between the bleomycin-induced lung fibrosis model and the small airway cell cultures generated from IPF patient lungs suggests that the small airway cultures maintain features of IPF and are suitable for modeling IPF in vitro.

### 4.2 Disease signatures of IPF cell cultures include cell cycle regulators, ciliary movement, and the predicted inhibition of the host immune response

A total of 1,991 DEGs were found to be in common between the control and TGFB-treated conditions of the IPF small airway cell culture (Figure 3A). These genes were analyzed using the Ingenuity Pathway Analysis tool to elucidate a genetic disease signature for IPF. The most significant pathway predicted to be activated in the IPF cultures was the kinetochore metaphase signaling pathway (Figure 3B). When paired with the predicted activation of cell cycle regulators ELDR, CKAP2L, and PCLAF, the evidence indicates that cell cycle control/progression may play a role in the pathogenesis of IPF (Figure 3C). This is further supported by the gene ontology results for the upregulated genes shared by the control and TGF-β1-treated IPF cell culture conditions (Figure 3D). Ontologies relevant to the regulation of the cell cycle include regulator of mitotic cytokinesis, spindle elongation, attachment of spindle microtubules to the kinetochore, and mitotic metaphase plate congression. The presence of these ontologies in the top 25 most significant entries is indicative of the important part that the cell cycle plays in the development of IPF. When viewed through the lens of cell cycle regulation, cellular senescence can be seen as advanced cell cycle arrest during the G1, G1/S, or G2 phases (33).

Also present in the list of the top 25 shared genes were various ontologies related to ciliary movement. Cilia are hair-like projections that are present in the inner lining of airways that serve to help move water, mucus, and debris. Ontologies linked to the function of cilia include mucociliary clearance, inner and outer dynein arm assembly, intraciliary retrograde and anterograde transport, regulation of cilium beat frequency, and regulation of cilium assembly (Figure 3D).

Inhibition of the host immune response is also over-represented by the common list of DEGs. Amongst the upstream regulators predicted to be inhibited are TNF, IL-6, and IL-B, all of which are key regulators of immune response and inflammation (34, 35). This inhibitory effect on the immune response is supported by the gene ontology entries for the common downregulated IPF genes; monocyte differentiation and response to macrophage colony-stimulating factor are both present in the top 25 gene ontologies. While the suppression of the immune system may seem counterintuitive, there is research to suggest that anti-fibrotic macrophage functions are reduced in IPF (36). Classical immunosuppression has been linked to poor clinical outcomes, further highlighting the observation that the IPF small airway cell cultures demonstrate the inhibition of host immune response regulators.

### 4.3 Post-COVID fibrosis disease signature demonstrates predicted activation of host immune response, inflammation, and remnants of viral infection

The comparison of post-COVID fibrosis patient cell cultures treated with TGF-β1 to their respective controls revealed a total of 374 DEGs in common (Figure 4A). Ingenuity pathway analysis of the shared genes revealed several immune system-related pathways that are predicted to be activated (Figure 4B). The most significant of the activated pathways is the eicosanoid signaling pathway. Eicosanoids are a class of lipid molecules that include prostaglandins and leukotrienes and are involved in inflammation signaling and response to infection (37). Macrophage activation also plays an important role in both the innate and adaptive immune systems; this pathway is predicted to be activated according to the IPA software. Interestingly, the interferon signaling pathway and cytokine storm signaling are both predicted to be inhibited. Interferons are a part of the “family” of cytokines, which includes lymphokines, chemokines, and interleukins as well (38). Research interest in the relationship between eicosanoids and cytokines is growing, although the exact nature has yet to be fully understood (38). Like macrophages, cytokines have roles in innate and adaptive immune responses.

The inhibition of interferon and cytokine signaling is further supported by the predicted inhibition of upstream regulators IRF7, IFN beta, and IFNL1 (Figure 4C). IRF7 is a transcription factor that is involved in the innate immune response (39). More specifically, IRF7 is an interferon regulatory factor tasked with aiding the immune system in identifying and responding to viral pathogens (39). IFNL1 is also known as interleukin-29 (IL-29) and is a member of the IL-10 cytokine family (40). Previous research has shown the IFNL group of proteins, consisting of IFNL1-3, demonstrated high levels of expression in airway epithelial cells during viral respiratory infections (40). These observations are supported by the gene ontologies associated with the downregulated genes; entries for response to Type I interferon, defense response to a virus, and response to interferon-alpha are present among the top 25 results (Figure 4E). The inhibition of these proteins and pathways could be indicative of a resolved COVID-19 infection. Although the initial infection may have concluded, the exacerbation of pulmonary fibrosis remains.

Interestingly, the gene ontology results for the upregulated gene set suggest that macrophage-mediated signaling is still active in epithelial cells (Figure 4D). Three entries related to antigen processing and presentation are present in the gene ontologies for upregulated genes. The major histocompatibility complex (MHC) proteins are crucial components of the adaptive immune system (41). MHC proteins can be divided into two classes, both of which are responsible for presenting antigens on the surface of cells for identification by a type of white blood cell known as T cells (41). Research studies have proposed that CD4+ T cells play a role in the chronic inflammation seen in autoimmune diseases (42). When taken together, the results of the shared post-COVID fibrosis genes seem to point towards a persistent inflammatory immune response following the resolution of infection by COVID-19.

### 4.4 IPF and post-COVID fibrosis datasets show conflicting regulation of the cell cycle and immune system responses

In order to better describe the transcriptional differences between the IPF and post-COVID fibrosis small airway cell cultures, DEG analysis was performed under both treatment conditions. A total of 1,606 DEGs were found to be shared by the IPF vs. COVID DEGs for the control and TGF-β1-treated cultures (Figure 5A). These DEGs were used for all downstream analysis, and therefore the following results can all be viewed in the context of the IPF cell cultures. Regulation of the cell cycle is a prominent component of the canonical pathway results; the kinetochore metaphase signaling pathway, mitotic roles of polo-like kinase, cyclins and cell cycle regulation, and cell cycle control of chromosomal replication are all predicted to be upregulated in the IPF cell cultures (Figure 5B). The activation of MYOD1 and RABL6 in the IPF cell culture predicted upstream regulators support the established importance of cell cycle regulation in the development of idiopathic pulmonary fibrosis (Figure 5C) (32). Conversely, TP53 and STAG2 are predicted to be inhibited; these proteins play a role in cell division and DNA repair, respectively (32, 43).

The gene ontologies for the upregulated genes in the IPF cell cultures seem to agree with the predicted activation of the cell cycle regulators, with many aspects of mitosis being represented; regulation of mitotic cytokinesis, mitotic spindle assembly checkpoint signaling, and mitotic DNA replication are just a few of these entries (Figure 5D). In a similar pattern to the gene ontology results observed in the IPF samples, intraciliary transport, (both anterograde and retrograde), can be found in the top 25 results for the IPF vs. COVID upregulated genes. Opposite to what is seen in the post-COVID fibrosis data, antigen processing and presentation are among the ontologies for the shared downregulated genes in the IPF cell cultures (Figure 5E).

Perhaps one of the most surprising results depicted is the proposed inhibition of the wound healing signaling pathway and the pulmonary fibrosis idiopathic signaling pathway in the IPF cell cultures. This observation will be explored more thoroughly in the context of TGF-β1 signaling in the following section.

### 4.5 Post-COVID fibrosis cell cultures demonstrate greater responsiveness to treatment with TGF-β1

TGF-β1 has long been accepted as a main driver of fibrosis in various tissues, including the lungs (44). This study explored the effect of TGF-β1 treatment on patient-derived small airway cultures with the addition of TGF-β1 to the cells before harvesting and sequencing. This effect was first investigated in the context of the idiopathic pulmonary fibrosis signaling pathway. A list of signature IPF genes was curated from relevant literature and organized into a heatmap to visualize and compare expression levels across all sample types (Figure 6A) (20). The samples were divided by disease type, (NL = normal lung, IPF = idiopathic pulmonary fibrosis, and COVID = post-COVID fibrosis), and by treatment, (Control = no treatment and TGF = treatment with TGF-β1). Upon examining the IPF gene signature heatmap, it is evident that the patterns of expression in IPF and post-COVID fibrosis samples are vastly differentIn our data, the post-COVID fibrosis samples that instead show greater expression for many of the selected genes; CCL2, CCND2, KRT7, and TRAM1 are all examples of genes in which the post-COVID fibrosis samples exhibit elevated gene expression even before treatment with TGF-β1. Given the critical role that TGF-β1 plays in the pathogenesis of fibrosis, the addition of TGF-β1 should, theoretically, further raise the expression of IPF-related genes. This trend is observed in the normal lung and post-COVID fibrosis samples. While most of the genes of interest in the IPF samples did show an increase in expression with the supplementation of TGF-β1, it was not nearly as dramatic as the pattern observed in the normal and post-COVID fibrosis samples. Genes associated with IPF appear to be more highly expressed in the post-COVID fibrosis samples, under both control and TGF-β1-treated conditions.

Analysis of IPF-related pathways supports the patterns revealed in the IPF gene heatmap (Figure 6B). As mentioned previously, both the wound healing pathway and the pulmonary fibrosis idiopathic signaling pathway are predicted to be inhibited; these two pathways are predicted to be activated in the post-COVID fibrosis data.

The differences in responsiveness to the TGF-β1 treatment led us to examine the gene expression levels of genes involved in the TGF-β1 signaling pathway (Figure 6C). For the normal lung samples, most genes associated with the TGF-β1 signaling pathway, such as JUN, MAP2K2, TGF-β1, and TGF-β2, are expressed at low levels until the culture is treated with TGF-β1. The majority of the TGF-β1 signaling pathway genes have higher expression levels in the post-COVID fibrosis samples, regardless of treatment status-similar to the changes observed with the IPF-related genes.

### 4.6 Higher levels of BMP signaling in IPF samples decrease the responsiveness to TGF-β1 treatment and lead to the predicted inhibition of key fibrosis-related pathways

There is a delicate balance that exists between the TGF-β1 signaling pathway and the BMP signaling pathway. Similar in structure, these two classes of signaling molecules are known to interact in ways that can either impede or promote the activity of the other (45). The equilibrium of this relationship is founded on the assembly and activation of the SMAD4 protein (Figure 7A) (45). If the BMP-related SMAD1/5/8 becomes phosphorylated, it can win the tug-of-war and use the SMAD4 protein to translocate into the nucleus. The presence of this complex in the nucleus leads to the translocation of SMAD6/7, a known inhibitor of TGF-β1 signaling, *out of* the nucleus. SMAD6/7 downregulates both TGFBR1 and SMAD2, key molecules in the TGFB signaling pathway (45). Without these proteins, the signaling cascade falls short of promoting collagen accumulation, inflammatory responses, and other characteristics of a fibrotic response. Studies have shown that BMPs can protect lung tissue from the fibrosis-promoting TGF-β1 signaling cascade (45). The IPF samples demonstrated higher levels of BMP signaling genes when visualized on a heatmap (Figure 7B); RT-PCR confirmed higher fold changes of BMP7, BMPR1A, and BMPR1B in the IPF samples.

Visually, both the normal and IPF samples have elevated levels of pSMAD1/5/8 nuclear localization; the quantification of these nuclei confirms this observation. In contrast, while the post-COVID fibrosis samples do have pSMAD1/5/8-positive cells, the staining is confined to the cytoplasm. The translocation of the pSMAD1/5/8-SMAD4 complex to the nucleus signals the release of SMAD6/7 into the cytoplasm, which then goes on to inhibit the TGF-β1 receptors. The presence of pSMAD1/5/8 in the nuclei of the IPF cells could indicate that the IPF samples have higher levels of BMP signaling. The elevated BMP signaling in the IPF samples may explain the decreased response of IPF small airway cell cultures to treatment with TGF-β1.

## 5 Conclusion

This study provides evidence that small airway cells cultured from IPF patient lungs display similar genetic characteristics to those of bleomycin-induced pulmonary fibrosis and previously published studies of IPF tissue and cell cultures. Small airway cells cultured from IPF and post-COVID fibrosis patient lungs all display unique genetic disease signatures. Common genes shared between the control and TGF-β1-treated IPF small airway cultures demonstrate the predicted activation of pathways and upstream regulators associated with cell cycle regulation; gene ontology of the upregulated genes also focuses on cell cycle checkpoints, along with cilia movement and intraciliary transport. Curiously, the pulmonary fibrosis idiopathic signaling pathway and TGF-β1 were both predicted to be inhibited in the IPF cell cultures. The post-COVID fibrosis cell cultures demonstrated a genetic disease signature that was largely focused on the immune system. The eicosanoid signaling pathway and macrophage activation were both predicted to be activated, along with upstream regulators CNOT7, NKX2-3, and TREX1. Antigen processing and presentation were among the gene ontology results for the common post-COVID fibrosis upregulated genes. The comparison of the IPF and post-COVID sequencing data show differences in genetic disease signatures and diverse responses to TGF-β1 treatment. Genes associated with IPF tend to be more highly expressed in the post-COVID fibrosis samples, under both control and TGF-β1-treated conditions. Small airway cultures from the post-COVID fibrosis lungs appear to be more responsive to TGF-β1 treatment than the IPF cultures. To further explore this observation, the expression of genes associated with the BMP signaling pathway was analyzed. The binding of BMP to its receptor triggers the translocation of pSMAD1/5/8 to the nucleus, which leads to SMAD6/7 leaving the nucleus and inhibiting the TGF-β1 receptors. Small airway cultures from IPF patients demonstrate higher expression levels of genes associated with BMP signaling than in the post-COVID fibrosis patient cultures. Immunohistochemistry of patient tissues revealed that IPF samples have elevated levels of pSMAD1/5/8 translocating to the nucleus, which can be an indicator of active BMP signaling. The data collected by this study suggests that IPF small airway cultures have more BMP signaling than post-COVID fibrosis cultures. The BMP signaling can inhibit the signaling pathway of TGF-β1, leading to the post-COVID fibrosis samples appearing to be more responsive to the TGF-β1 treatment. The results of this study demonstrate the importance of TGF-β1 and BMP signaling in the pathogenesis of both IPF and post-COVID fibrosis and provide new insights that could lead to future therapies.

## 7 Author Contributions

K.U., M.A., K.T., T.J., and X.L. performed the biological experiments and cell culture techniques described in this study. Immunostaining of tissue samples was carried out by K.U. and X.L. A.M.P., M.T.K., A.C.B., T.E.J., C.L., D.W.C., and R.E.G. coordinated the collection of human lung tissue samples and consenting of patients. K.U., S.P., D.L., J.P., and B.C. analyzed the bulk RNA-sequencing data and carried out differential expression analysis. K.U. and X.L. wrote the manuscript. The manuscript was read and approved by all authors before submission.

## 9 Acknowledgments

The authors of this study would like to thank Lung Bioengineering, Inc. and the persons who generously donated their lungs for research purposes. The authors would also like to thank the biorepository team at Corewell Health for their assistance. The Genomics Core at the Van Andel Institute carried out the bulk RNA-sequencing for this project.

## 10 Data Availability Statement

All data presented in this study are available upon request from the corresponding authors. The data collected from RNA-sequencing has been made available in the Gene Expression Omnibus database (GSE225549).

## 11 Ethics Statement

Written informed consent was collected from all patients who participated in this study (Spectrum Health IRB no.: 2017-198). The protocols described in this study were carried out in compliance with the ethical regulations determined by the local institutional review board.

