## Supplemental Figures for "Bulk RNA-Sequencing of small airway cell cultures from IPF and post-COVID lung fibrosis patients illustrates disease signatures and differential responses to TGF-*β*1 treatment"

Reda E. Girgis, MD

Corewell Health

Michigan State University College of Human Medicine

330 Barclay Ave. NW, Suite 200

Grand Rapids, MI 49503

**Supplementary Table 1.** Supplementary Table 1. Summary of clinical data for all human samples used for RNA-sequencing, RT-PCR, and IHC (LUL = left upper lobe, RML = right middle lobe).

| **Sample Name** | **Sample ID** | **Diagnosis** | **Age** | **Gender** | **Sample Source** |
| --- | --- | --- | --- | --- | --- |
| SR-22-123 | NL1 | Normal | 82 | Male | Lobectomy |
| SR-22-270 | NL2 |  | 65 | Male | Lobectomy |
| SR-21-424 | NL3 |  | 72 | Female | Lobectomy |
| AILN102 | AILN102 |  | N/A | N/A | Donor Lung |
| AILS058 | AILS058 |  | N/A | N/A | Donor Lung |
| AJEK269 | AJEK269 |  | N/A | N/A | Donor Lung |
| IPF-116 | IPF1 | IPF | 69 | Male | Explant- LUL |
| IPF-130 | IPF2 |  | 64 | Male | Explant- LUL |
| IPF-146 | IPF3 |  | 60 | Male | Explant- LUL |
| IPF-149 | IPF-149 |  | 52 | Male | Explant- RML |
| IPF-162 | IPF-162 |  | 57 | Male | Explant- LUL |
| IPF-188 | IPF-188 |  | 67 | Male | Explant- LUL |
| IPF-218 | IPF-218 |  | 75 | Male | Explant- LUL |
| IPF-229 | IPF-229 |  | 73 | Male | Explant- RML |
| IPF-230 | IPF-230 |  | 70 | Male | Explant- LUL |
| IPF-234 | IPF-234 |  | 71 | Male | Explant- LUL |
| COVID-126 | COVID1 | Post-COVID Fibrosis | 64 | Male | Explant- LUL |
| COVID-134 | COVID2 |  | 56 | Male | Explant- LUL |
| COVID-140 | COVID3 |  | 58 | Male | Explant- LUL |
| COVID-126 | COVID-126 |  | 64 | Male | Explant- LUL |
| COVID-134 | COVID-134 |  | 56 | Male | Explant- LUL |
| COVID-140 | COVID-140 |  | 58 | Male | Explant- LUL |
| COVID-160 | COVID-160 |  | 31 | Male | Explant- LUL |
| COVID-223 | COVID-223 |  | 56 | Male | Explant- RML |

**Supplementary Table 2.** List of primer sequences used for RT-PCR analysis.

| **Target** | **Sequence (5’-3’)** |
| --- | --- |
| Human ACTB Forward | GGATCAGCAAGCAGGAGTATG |
| Human ACTB Reverse | AGAAAGGGTGTAACGCAACTAA |
| Human BMP7 Forward | CCTAACCAAGTGTCCCGATTT |
| Human BMP7 Reverse | GGAGGCTGAGTGCATACTATT T |
| Human BMPR1A Forward | AGTGGGTCTGGACTACCTTTA |
| Human BMPR1A Reverse | GCCCATCCATACTTCTCCATATC |
| Human BMPR1B Forward | CCTATACACCACAGGGCTTTAC |
| Human BMPR1B Reverse | CGAGGTCTGGTTTCTTGTCTT |
| Human FOXM1 Forward | CAGGGTGGTCCGTGTAAATAG |
| Human FOXM1 Reverse | CTTCTGGCAGTCTCTGGATAAT |

**Supplementary Table 3.** List of differentially expressed genes shared by the IPF patient-derived small airway cell cultures and mouse lungs treated with bleomycin and adenovirus-mediated ATP12A overexpression.

|  | **Human Small Airway Cell Culture** | | | **BLEO-Induced Fibrosis Mouse Model** | | |
| --- | --- | --- | --- | --- | --- | --- |
| **Gene Symbol** | **log2FoldChange** | **pvalue** | **padj** | **log2FoldChange** | **pvalue** | **padj** |
| ABCA8 | -3.993033265 | 3.49E-18 | 5.84E-14 | -1.638814335 | 0.00000144 | 0.02100958 |
| ANLN | 3.591178948 | 1.73E-24 | 2.90E-20 | 2.707033397 | 2.89E-17 | 4.5E-13 |
| ARHGAP11A | 2.584816607 | 8.82E-13 | 1.46E-08 | 1.77595776 | 6.37E-09 | 0.0000956 |
| ASF1B | 1.95379182 | 7.60E-07 | 0.012116659 | 2.094626335 | 8.46E-10 | 0.0000128 |
| ASPM | 5.21444258 | 9.59E-40 | 1.62E-35 | 2.744580219 | 5.38E-17 | 8.35E-13 |
| ATP10B | 1.809431324 | 2.72E-06 | 0.042953443 | 2.251010814 | 0.00000035 | 0.005164774 |
| AURKA | 1.949441308 | 6.07E-09 | 9.91E-05 | 2.613630526 | 9.78E-14 | 1.51E-09 |
| AURKB | 3.157763803 | 1.04E-12 | 1.73E-08 | 2.486337464 | 2.37E-13 | 3.65E-09 |
| BAALC | 1.832297719 | 3.38E-07 | 0.005415995 | 3.299397385 | 4.79E-14 | 7.39E-10 |
| BIRC5 | 4.460170179 | 2.03E-24 | 3.41E-20 | 2.482683103 | 2.58E-13 | 3.97E-09 |
| BRIP1 | 2.161111279 | 7.39E-08 | 0.001194684 | 1.882730615 | 6.83E-08 | 0.0010164 |
| BUB1 | 3.741772384 | 1.86E-25 | 3.12E-21 | 2.009111108 | 1.3E-09 | 0.0000196 |
| BUB1B | 3.977213064 | 3.24E-21 | 5.44E-17 | 1.970483434 | 1.54E-09 | 0.0000231 |
| CCDC80 | 1.74807717 | 2.35E-08 | 0.000381701 | 2.824318987 | 1.25E-21 | 1.95E-17 |
| CCNB1 | 2.675068647 | 1.05E-16 | 1.76E-12 | 3.248813157 | 2.41E-20 | 3.76E-16 |
| CCNB2 | 3.033850463 | 2.79E-17 | 4.67E-13 | 2.819478056 | 1.54E-14 | 2.38E-10 |
| CDC20 | 3.444637882 | 1.95E-21 | 3.28E-17 | 2.160411491 | 2.4E-11 | 0.000000365 |
| CDC25C | 3.548497923 | 1.29E-10 | 2.13E-06 | 3.722718422 | 1.28E-15 | 1.98E-11 |
| CDCA2 | 3.292000884 | 1.97E-15 | 3.29E-11 | 2.235251413 | 2.12E-09 | 0.0000319 |
| CDCA3 | 2.269450675 | 1.69E-07 | 0.00271945 | 2.368458822 | 1.27E-11 | 0.000000195 |
| CDCA8 | 2.083081994 | 1.76E-06 | 0.027818061 | 2.50947808 | 1.51E-10 | 0.00000229 |
| CDK1 | 3.443494979 | 5.61E-22 | 9.42E-18 | 3.175552132 | 8.36E-20 | 1.3E-15 |
| CDKN3 | 4.753601899 | 5.14E-20 | 8.61E-16 | 2.968202624 | 3.25E-12 | 4.97E-08 |
| CENPE | 4.451068212 | 3.67E-28 | 6.18E-24 | 3.066564556 | 6.87E-22 | 1.07E-17 |
| CENPF | 3.740922407 | 8.79E-26 | 1.48E-21 | 3.146206085 | 1.32E-19 | 2.06E-15 |
| CENPH | 2.512553921 | 7.85E-11 | 1.30E-06 | 2.359971769 | 7.75E-08 | 0.001152793 |
| CENPI | 3.519883553 | 1.56E-15 | 2.60E-11 | 2.108831311 | 3.54E-08 | 0.00052773 |
| CEP55 | 4.507287565 | 1.73E-23 | 2.91E-19 | 2.018549111 | 0.000000136 | 0.002017266 |
| CHI3L1 | -3.763701678 | 6.98E-15 | 1.16E-10 | -2.522864526 | 3.55E-13 | 5.45E-09 |
| CKAP2 | 1.60000844 | 2.48E-07 | 0.003980656 | 3.005316029 | 3.31E-20 | 5.17E-16 |
| CKAP2L | 4.592690409 | 3.31E-26 | 5.57E-22 | 2.458081346 | 7.98E-13 | 1.23E-08 |
| CKS1B | 1.734328311 | 4.60E-07 | 0.007349884 | 1.805854277 | 3.31E-08 | 0.000494534 |
| CKS2 | 1.821578375 | 3.97E-08 | 0.000643891 | 1.572192756 | 0.00000339 | 0.049255138 |
| CLSPN | 2.941490538 | 1.96E-06 | 0.030950848 | 2.767224094 | 2.35E-14 | 3.63E-10 |
| CTSS | 1.553004978 | 2.18E-07 | 0.003505082 | 3.495538376 | 1.23E-31 | 1.93E-27 |
| CUX2 | -3.827203438 | 4.36E-09 | 7.12E-05 | -1.82450497 | 0.00000219 | 0.031915781 |
| CYP2E1 | -2.050725725 | 5.48E-08 | 0.000886308 | -2.490818996 | 8.14E-10 | 0.0000123 |
| DBF4 | 1.643656384 | 2.17E-07 | 0.003477477 | 1.654690025 | 0.00000017 | 0.002515463 |
| DEPDC1 | 4.563262123 | 2.52E-29 | 4.25E-25 | 3.188369132 | 1.2E-16 | 1.86E-12 |
| DIAPH3 | 2.693084429 | 2.12E-10 | 3.50E-06 | 2.360484672 | 7.86E-14 | 1.21E-09 |
| DLGAP5 | 4.469128963 | 9.15E-27 | 1.54E-22 | 2.898733271 | 7.5E-17 | 1.16E-12 |
| DTL | 3.041522919 | 3.61E-08 | 0.000584698 | 1.802731868 | 0.000000694 | 0.010194208 |
| ESCO2 | 3.220482723 | 2.61E-12 | 4.32E-08 | 1.754907653 | 0.000000962 | 0.014095944 |
| ESPL1 | 2.34114904 | 2.03E-06 | 0.032145028 | 1.802276233 | 0.000000054 | 0.000804787 |
| FAM83D | 3.095290673 | 9.70E-18 | 1.62E-13 | 2.714565853 | 2.99E-13 | 4.59E-09 |
| FGF1 | -4.122833423 | 3.57E-20 | 5.99E-16 | -2.04385015 | 8.17E-09 | 0.000122492 |
| FOXM1 | 2.847076255 | 9.10E-14 | 1.51E-09 | 2.607188639 | 5.37E-13 | 8.26E-09 |
| G0S2 | -1.535892928 | 1.13E-06 | 0.017899415 | -2.423647415 | 2.19E-10 | 0.00000332 |
| GAS2L3 | 2.547982381 | 2.26E-12 | 3.74E-08 | 2.695206066 | 7.61E-14 | 1.17E-09 |
| GATA3 | -3.1916 | 1.93E-13 | 3.21E-09 | -1.827759615 | 0.000000634 | 0.009326637 |
| GLB1L3 | -3.650268775 | 9.95E-15 | 1.66E-10 | -2.916833684 | 8.88E-08 | 0.001320051 |
| GTSE1 | 3.13690852 | 1.25E-12 | 2.08E-08 | 3.208153345 | 3.68E-19 | 5.73E-15 |
| HES2 | -2.071342015 | 9.88E-12 | 1.64E-07 | -4.772182941 | 1.03E-11 | 0.000000157 |
| HLA-DOB | -2.976365603 | 1.55E-09 | 2.55E-05 | -2.344189369 | 6.88E-10 | 0.0000104 |
| HMMR | 4.576025233 | 5.00E-31 | 8.43E-27 | 3.239538624 | 9.17E-22 | 1.43E-17 |
| IQGAP3 | 2.966831932 | 1.05E-11 | 1.74E-07 | 2.882438307 | 4.83E-16 | 7.49E-12 |
| KCNA2 | -3.415445667 | 1.24E-12 | 2.06E-08 | -1.912294426 | 5.55E-08 | 0.000827795 |
| KIF11 | 2.809925258 | 1.07E-14 | 1.78E-10 | 2.282987176 | 2.55E-13 | 3.93E-09 |
| KIF14 | 4.512817902 | 2.07E-28 | 3.49E-24 | 1.83044832 | 0.000000284 | 0.004200743 |
| KIF15 | 3.192534898 | 2.29E-12 | 3.79E-08 | 2.239050038 | 1.6E-11 | 0.000000244 |
| KIF18B | 2.584952657 | 3.14E-08 | 0.000509983 | 2.604273461 | 1.72E-12 | 2.63E-08 |
| KIF20A | 3.856821281 | 1.91E-24 | 3.22E-20 | 2.856397971 | 2.61E-14 | 4.03E-10 |
| KIF20B | 2.843057895 | 1.43E-16 | 2.39E-12 | 2.059821568 | 1.98E-09 | 0.0000299 |
| KIF23 | 2.152179624 | 7.30E-10 | 1.20E-05 | 1.754636013 | 2.18E-08 | 0.00032516 |
| KIF2C | 3.179096648 | 2.28E-16 | 3.81E-12 | 2.872916874 | 7.31E-16 | 1.13E-11 |
| KIF4A | 3.46893304 | 4.19E-16 | 7.00E-12 | 2.204145335 | 3.63E-11 | 0.000000553 |
| MELK | 2.497545252 | 3.82E-13 | 6.34E-09 | 3.092303698 | 1.94E-17 | 3.02E-13 |
| MYBL2 | 3.568059195 | 1.43E-10 | 2.36E-06 | 2.112780381 | 2.19E-09 | 0.0000329 |
| NCAPG | 5.504499814 | 2.14E-19 | 3.59E-15 | 2.838381361 | 2.35E-12 | 0.000000036 |
| NCAPH | 3.07362529 | 2.63E-16 | 4.40E-12 | 1.557680661 | 0.000000762 | 0.01118945 |
| NDC80 | 4.878629995 | 1.50E-22 | 2.53E-18 | 2.325181682 | 3.12E-11 | 0.000000476 |
| NEIL3 | 5.118960206 | 6.96E-19 | 1.17E-14 | 2.412314086 | 1.4E-10 | 0.00000212 |
| NUF2 | 3.629550662 | 2.76E-20 | 4.62E-16 | 2.638280112 | 2.5E-14 | 3.86E-10 |
| NUSAP1 | 4.229611611 | 5.60E-26 | 9.42E-22 | 1.626802364 | 0.000000506 | 0.007459343 |
| PARPBP | 2.788544129 | 6.45E-13 | 1.07E-08 | 2.458793814 | 1.31E-09 | 0.0000198 |
| PBK | 4.393434419 | 9.53E-21 | 1.60E-16 | 2.126374539 | 8.47E-09 | 0.000127059 |
| PDE10A | 1.854782614 | 3.72E-08 | 0.000603743 | 2.356065744 | 8.32E-11 | 0.00000127 |
| PLK1 | 2.130816394 | 3.34E-10 | 5.50E-06 | 2.921493635 | 4.43E-18 | 6.89E-14 |
| PLK4 | 3.012228903 | 6.31E-14 | 1.05E-09 | 1.55599945 | 0.000000999 | 0.014630721 |
| POLQ | 2.895507344 | 5.81E-14 | 9.67E-10 | 1.821802023 | 0.000000756 | 0.011110652 |
| PRC1 | 3.516581899 | 1.86E-18 | 3.11E-14 | 2.557607253 | 2.66E-14 | 4.1E-10 |
| PRR11 | 3.526826257 | 1.16E-19 | 1.94E-15 | 2.461560947 | 1.93E-12 | 2.96E-08 |
| PTN | 4.148869941 | 1.22E-16 | 2.04E-12 | 2.597210918 | 9.08E-14 | 1.4E-09 |
| RACGAP1 | 2.262467062 | 5.20E-11 | 8.57E-07 | 2.303999354 | 1.24E-13 | 1.91E-09 |
| RASGEF1A | -3.01597403 | 1.43E-14 | 2.38E-10 | -1.676707939 | 0.000000431 | 0.006360813 |
| RRM2 | 3.36632084 | 3.38E-09 | 5.53E-05 | 2.364442736 | 2.52E-12 | 3.86E-08 |
| SHCBP1 | 4.004558087 | 8.26E-13 | 1.37E-08 | 3.262943564 | 1.32E-18 | 2.05E-14 |
| SKA1 | 4.263450243 | 1.30E-11 | 2.15E-07 | 2.58046943 | 4.19E-08 | 0.000624319 |
| SKA3 | 2.725294888 | 2.01E-12 | 3.33E-08 | 2.480439117 | 7.11E-11 | 0.00000108 |
| SLC38A5 | -2.965043922 | 7.22E-09 | 0.000117863 | -2.607331297 | 9.18E-13 | 1.41E-08 |
| SMC2 | 1.901264576 | 2.61E-09 | 4.27E-05 | 1.788142857 | 3.92E-09 | 0.000059 |
| SPC24 | 3.26643022 | 3.47E-09 | 5.67E-05 | 1.79334653 | 0.000000738 | 0.010841979 |
| SPC25 | 4.72198444 | 2.80E-15 | 4.66E-11 | 1.854953939 | 0.000000175 | 0.002591632 |
| STIL | 1.565226181 | 1.65E-06 | 0.026186801 | 2.511477078 | 4.4E-13 | 6.76E-09 |
| TICRR | 2.251179491 | 8.40E-07 | 0.013374183 | 2.933531856 | 4.56E-14 | 7.03E-10 |
| TOP2A | 5.17324465 | 7.82E-39 | 1.32E-34 | 2.619032466 | 2.38E-15 | 3.68E-11 |
| TPX2 | 3.215438061 | 2.74E-17 | 4.58E-13 | 3.062822871 | 6.36E-20 | 9.92E-16 |
| TROAP | 3.333009863 | 1.21E-14 | 2.01E-10 | 3.434628786 | 6.47E-13 | 9.94E-09 |
| TTK | 4.299426037 | 7.31E-28 | 1.23E-23 | 2.934262146 | 1.48E-12 | 2.27E-08 |
| UBE2C | 4.552465339 | 4.29E-19 | 7.19E-15 | 2.097370485 | 7.14E-08 | 0.001062203 |
| WNT10A | 1.93656679 | 7.17E-07 | 0.011442398 | 6.746164258 | 1.1E-19 | 1.72E-15 |

**Supplementary Table 4.** Top 100 canonical pathway analysis results for the IPF vs. Normal DEG comparison, as calculated by the Ingenuity Pathway Analysis tool (results sorted by -log(p-value)). A positive z-score indicates that the pathway is predicted to be activated, while a negative z-score signifies predicted inhibition. “N/A” is used when not enough data exists to determine the activation state of the pathway.

| **IPF Common Genes - IPA Canonical Pathway Analysis** | | | |
| --- | --- | --- | --- |
| **Ingenuity Canonical Pathways** | **-log(p-value)** | **Ratio** | **z-score** |
| Hepatic Fibrosis / Hepatic Stellate Cell Activation | 12.4 | 0.402 | N/A |
| Kinetochore Metaphase Signaling Pathway | 8.12 | 0.414 | 3.656 |
| Wound Healing Signaling Pathway | 8.08 | 0.329 | -5.159 |
| Pulmonary Fibrosis Idiopathic Signaling Pathway | 7.75 | 0.307 | -5.511 |
| Axonal Guidance Signaling | 7.41 | 0.277 | N/A |
| Chondroitin Sulfate Biosynthesis (Late Stages) | 7.25 | 0.52 | -0.784 |
| Xenobiotic Metabolism CAR Signaling Pathway | 7.08 | 0.34 | 1.861 |
| Chondroitin Sulfate Biosynthesis | 6.86 | 0.483 | -1.134 |
| LPS/IL-1 Mediated Inhibition of RXR Function | 6.22 | 0.307 | -1.414 |
| GP6 Signaling Pathway | 6.06 | 0.362 | -4.629 |
| Xenobiotic Metabolism PXR Signaling Pathway | 5.89 | 0.323 | 1.778 |
| Dermatan Sulfate Biosynthesis (Late Stages) | 5.88 | 0.489 | -0.626 |
| Dermatan Sulfate Biosynthesis | 5.87 | 0.45 | -1.347 |
| Superpathway of Melatonin Degradation | 5.83 | 0.433 | 0.186 |
| G-Protein Coupled Receptor Signaling | 5.79 | 0.25 | -3.618 |
| Breast Cancer Regulation by Stathmin1 | 5.65 | 0.256 | -2.887 |
| Melatonin Degradation I | 5.53 | 0.435 | 0.192 |
| Phagosome Formation | 5.35 | 0.247 | -4.575 |
| CREB Signaling in Neurons | 5.29 | 0.252 | -2.833 |
| Regulation of Cellular Mechanics by Calpain Protease | 5.23 | 0.382 | -2.84 |
| STAT3 Pathway | 5.23 | 0.341 | -1.257 |
| ID1 Signaling Pathway | 5.16 | 0.308 | -1.016 |
| Xenobiotic Metabolism Signaling | 5.16 | 0.285 | N/A |
| Estrogen-mediated S-phase Entry | 5.15 | 0.577 | 1.291 |
| Nicotine Degradation III | 5.08 | 0.431 | 0.408 |
| Nicotine Degradation II | 4.92 | 0.409 | 0.962 |
| Molecular Mechanisms of Cancer | 4.83 | 0.26 | N/A |
| Agranulocyte Adhesion and Diapedesis | 4.81 | 0.3 | N/A |
| RHOGDI Signaling | 4.7 | 0.295 | 3.452 |
| Glioblastoma Multiforme Signaling | 4.57 | 0.31 | -1.265 |
| Actin Cytoskeleton Signaling | 4.55 | 0.287 | -3.904 |
| S100 Family Signaling Pathway | 4.3 | 0.236 | -3.113 |
| Tumor Microenvironment Pathway | 4.29 | 0.302 | -3.81 |
| Signaling by Rho Family GTPases | 3.97 | 0.273 | -3.221 |
| ILK Signaling | 3.95 | 0.289 | -2.887 |
| Bupropion Degradation | 3.9 | 0.52 | 0.832 |
| Role Of Osteoclasts In Rheumatoid Arthritis Signaling Pathway | 3.9 | 0.265 | -4.756 |
| Serotonin Degradation | 3.8 | 0.366 | 0.392 |
| Pathogen Induced Cytokine Storm Signaling Pathway | 3.79 | 0.256 | -4.412 |
| Cardiac Hypertrophy Signaling (Enhanced) | 3.75 | 0.242 | -1.782 |
| Semaphorin Signaling in Neurons | 3.64 | 0.377 | N/A |
| Atherosclerosis Signaling | 3.63 | 0.308 | N/A |
| Semaphorin Neuronal Repulsive Signaling Pathway | 3.62 | 0.3 | -0.617 |
| Agrin Interactions at Neuromuscular Junction | 3.59 | 0.362 | -1.964 |
| PAK Signaling | 3.58 | 0.316 | -2.041 |
| Integrin Signaling | 3.55 | 0.278 | -3.501 |
| Neuroinflammation Signaling Pathway | 3.51 | 0.259 | -1.016 |
| Osteoarthritis Pathway | 3.46 | 0.271 | -1.605 |
| Role of Osteoblasts, Osteoclasts and Chondrocytes in Rheumatoid Arthritis | 3.4 | 0.272 | N/A |
| Estrogen Biosynthesis | 3.33 | 0.4 | 0.728 |
| Cellular Effects of Sildenafil (Viagra) | 3.32 | 0.293 | N/A |
| Sperm Motility | 3.3 | 0.265 | -1.3 |
| Pulmonary Healing Signaling Pathway | 3.27 | 0.276 | -2.562 |
| Heparan Sulfate Biosynthesis | 3.26 | 0.33 | -0.928 |
| Actin Nucleation by ARP-WASP Complex | 3.18 | 0.323 | -1.807 |
| Heparan Sulfate Biosynthesis (Late Stages) | 3.16 | 0.333 | -0.577 |
| Cell Cycle: G2/M DNA Damage Checkpoint Regulation | 3.16 | 0.38 | -2.357 |
| Germ Cell-Sertoli Cell Junction Signaling | 3.15 | 0.282 | N/A |
| Sertoli Cell-Sertoli Cell Junction Signaling | 3.13 | 0.272 | N/A |
| Endocannabinoid Neuronal Synapse Pathway | 3.1 | 0.289 | -1.029 |
| Granulocyte Adhesion and Diapedesis | 3.08 | 0.275 | N/A |
| Thyroid Hormone Metabolism II (via Conjugation and/or Degradation) | 3.02 | 0.4 | -0.5 |
| Hepatic Fibrosis Signaling Pathway | 3.02 | 0.241 | -4.72 |
| Mitotic Roles of Polo-Like Kinase | 2.97 | 0.343 | 1.5 |
| Caveolar-mediated Endocytosis Signaling | 2.97 | 0.333 | N/A |
| Neutrophil Extracellular Trap Signaling Pathway | 2.86 | 0.24 | 2.915 |
| CDX Gastrointestinal Cancer Signaling Pathway | 2.86 | 0.267 | 0.816 |
| Apelin Liver Signaling Pathway | 2.86 | 0.444 | -1.732 |
| Ephrin Receptor Signaling | 2.86 | 0.267 | -2.263 |
| Role Of Osteoblasts In Rheumatoid Arthritis Signaling Pathway | 2.82 | 0.258 | -3.048 |
| Notch Signaling | 2.8 | 0.395 | -0.302 |
| Regulation of Actin-based Motility by Rho | 2.77 | 0.296 | -2.132 |
| Aryl Hydrocarbon Receptor Signaling | 2.75 | 0.277 | -0.853 |
| Paxillin Signaling | 2.73 | 0.299 | -2.683 |
| Transcriptional Regulatory Network in Embryonic Stem Cells | 2.69 | 0.352 | N/A |
| Histidine Degradation VI | 2.69 | 0.429 | -0.333 |
| MSP-RON Signaling Pathway | 2.69 | 0.345 | N/A |
| Xenobiotic Metabolism AHR Signaling Pathway | 2.64 | 0.31 | 0.192 |
| Dopamine Degradation | 2.63 | 0.406 | 0.277 |
| Airway Pathology in Chronic Obstructive Pulmonary Disease | 2.56 | 0.288 | N/A |
| Adrenomedullin signaling pathway | 2.55 | 0.261 | -1.18 |
| Dilated Cardiomyopathy Signaling Pathway | 2.5 | 0.273 | 0.898 |
| Cyclins and Cell Cycle Regulation | 2.47 | 0.306 | 1.877 |
| Colorectal Cancer Metastasis Signaling | 2.46 | 0.247 | -2.177 |
| Glioma Invasiveness Signaling | 2.42 | 0.315 | -1.147 |
| Macrophage Alternative Activation Signaling Pathway | 2.41 | 0.257 | -0.277 |
| Gustation Pathway | 2.36 | 0.256 | 0 |
| HMGB1 Signaling | 2.32 | 0.263 | -1.347 |
| Intrinsic Prothrombin Activation Pathway | 2.31 | 0.357 | -2.84 |
| Pregnenolone Biosynthesis | 2.31 | 0.407 | 0 |
| Regulation of eIF4 and p70S6K Signaling | 2.31 | 0.26 | -1.155 |
| Gαi Signaling | 2.31 | 0.271 | -2.556 |
| Acetone Degradation I (to Methylglyoxal) | 2.2 | 0.349 | 1.291 |
| Leukocyte Extravasation Signaling | 2.18 | 0.254 | -2.53 |
| Reelin Signaling in Neurons | 2.17 | 0.268 | -1.89 |
| Sphingosine-1-phosphate Signaling | 2.17 | 0.275 | -0.928 |
| Bladder Cancer Signaling | 2.14 | 0.276 | -0.535 |
| Role of Tissue Factor in Cancer | 2.14 | 0.276 | N/A |
| Synaptogenesis Signaling Pathway | 2.07 | 0.235 | -4.154 |
| Human Embryonic Stem Cell Pluripotency | 2.03 | 0.249 | -1.131 |

**Supplementary Table 5.** Top 100 upstream regulator results for the IPF vs. Normal DEG comparison, as calculated by the Ingenuity Pathway Analysis tool (results sorted by p-value of the overlap) A positive z-score indicates that the regulator is predicted to be activated, while a negative z-score signifies predicted inhibition.

| **IPF Common Genes - IPA Upstream Regulator Analysis** | | | |
| --- | --- | --- | --- |
| **Upstream Regulator** | **Expr Log Ratio** | **Activation z-score** | **p-value of overlap** |
| Eldr |  | 7.319 | 1.13E-37 |
| TGFB1 |  | -9.332 | 8.24E-36 |
| TNF | -3.332 | -6.573 | 9.75E-31 |
| ERBB2 |  | -1.533 | 3.07E-28 |
| IL1B | -4.262 | -3.92 | 3.12E-28 |
| YAP1 |  | -2.129 | 6.68E-25 |
| CTNNB1 |  | -3.522 | 2.2E-24 |
| FOXM1 | 2.762 | 2.633 | 2.26E-24 |
| CKAP2L | 4.516 | 5.642 | 5.31E-24 |
| ESR2 |  | -0.378 | 6.9E-24 |
| TP53 |  | -5.282 | 1.16E-23 |
| TP63 |  | 0.126 | 1.17E-23 |
| PGR | -2.67 | -0.64 | 3.13E-23 |
| Vegf |  | 1.201 | 6.2E-23 |
| MAP2K1 |  | -3.49 | 1.4E-21 |
| AGT | -2.963 | -6.13 | 1.44E-21 |
| IL6 | -4.48 | -4.097 | 4.07E-21 |
| CDKN1A | -1.093 | -1.773 | 6.76E-21 |
| CCND1 | -2.582 | 0.26 | 1.78E-20 |
| HRAS |  | -1.046 | 2.8E-20 |
| CSF2 | -2 | 1.451 | 3.55E-20 |
| CG |  | -2.968 | 1.18E-19 |
| ZBTB17 |  |  | 1.62E-19 |
| STAT3 |  | -1.533 | 1.99E-19 |
| CEBPB |  | 2.639 | 2.3E-19 |
| PCLAF | 3.454 | 5.261 | 2.39E-19 |
| PTGER2 | 2.769 | 2.889 | 3.29E-19 |
| SMARCA4 |  | -2.976 | 2.22E-18 |
| MYOD1 |  | -0.12 | 1.61E-17 |
| ERBB3 |  | -0.463 | 9.59E-17 |
| SP1 |  | -3.624 | 9.88E-17 |
| MRTFB |  | -3.028 | 1.38E-16 |
| TCF3 |  | -2.978 | 1.42E-16 |
| AMBRA1 |  | 3.647 | 1.66E-16 |
| NUPR1 |  | -8.034 | 1.96E-16 |
| NKX2-3 |  | -0.226 | 2.1E-16 |
| AHR |  | 0.858 | 2.24E-16 |
| CDKN2A | -2.418 | -5.089 | 2.84E-16 |
| IFNG |  | -3.853 | 3.48E-16 |
| Tgf beta |  | -5.273 | 9.47E-16 |
| E2F4 |  | -1.612 | 1.66E-15 |
| GLI1 |  | -1.302 | 2.58E-15 |
| KRAS |  | -0.425 | 4.49E-15 |
| estrogen receptor |  | 2.611 | 1.47E-14 |
| TP73 | 2.136 | -1.332 | 2.4E-14 |
| HDAC1 |  | 2.609 | 2.7E-14 |
| FGF2 | -1.271 | -3.626 | 2.75E-14 |
| TWIST1 |  | -2.567 | 7.84E-14 |
| CASR | -2.238 | 1.13 | 1.24E-13 |
| HGF |  | 1.067 | 1.55E-13 |
| AREG | -1.564 | 1.767 | 1.58E-13 |
| MYC | -1.536 | 3.046 | 1.94E-13 |
| WWTR1 |  | -0.126 | 4.97E-13 |
| IGF1 | 2.18 | -2.921 | 5.42E-13 |
| YY1 |  | 0.376 | 6.62E-13 |
| SMAD3 |  | -5.08 | 7.96E-13 |
| WNT3A | 2.207 | -3.069 | 8.87E-13 |
| BRD4 |  | -0.735 | 1.22E-12 |
| RABL6 |  | 5.754 | 1.43E-12 |
| STAT1 |  | -0.733 | 1.69E-12 |
| NRAS |  | -0.316 | 1.71E-12 |
| Interferon alpha |  | 0.174 | 1.72E-12 |
| SP3 | 1.176 | 0.252 | 2.27E-12 |
| RB1 |  | -1.642 | 2.88E-12 |
| KDM1A |  | 3.467 | 3.41E-12 |
| PDGF BB |  | -2.635 | 4.12E-12 |
| NRG1 |  | -2.541 | 4.27E-12 |
| TGFB2 | -1.298 | -4.838 | 4.83E-12 |
| let-7 |  | -0.697 | 6.12E-12 |
| EDN1 | -2.459 | -5.168 | 7.34E-12 |
| MRTFA | -1.046 | -3.174 | 7.73E-12 |
| AR |  | -1.507 | 9E-12 |
| EZH2 |  | 0.142 | 9.2E-12 |
| PDLIM2 |  | 0.493 | 9.21E-12 |
| ESR1 |  | 2.595 | 1.04E-11 |
| MYCN |  | 5.907 | 1.08E-11 |
| FKBP10 | -4.06 | 4.802 | 1.44E-11 |
| KDM3B |  | -1.89 | 1.52E-11 |
| ITGB1 | -1.279 | -1.1 | 2.11E-11 |
| F2 |  | -6.066 | 2.17E-11 |
| IL13 |  | -0.389 | 2.36E-11 |
| NOTCH3 |  | -3.214 | 2.47E-11 |
| HNRNPA2B1 |  | -1.982 | 2.5E-11 |
| PELP1 |  | -1.436 | 2.65E-11 |
| FOS |  | -3.835 | 3.98E-11 |
| ZNF768 |  | 4.347 | 4.72E-11 |
| ETV5 | -2.411 | 0.148 | 4.73E-11 |
| MAPK1 |  | -2.687 | 4.9E-11 |
| HIF1A | -1.288 | -3.832 | 4.92E-11 |
| MYB | 2.351 | 0.072 | 5.43E-11 |
| EGF |  | -4.324 | 6.16E-11 |
| MACROH2A1 |  | 1.616 | 6.21E-11 |
| RARA |  | 1.634 | 6.44E-11 |
| E2f |  | 2.875 | 6.9E-11 |
| SOX9 | -1.153 | -2.267 | 8.14E-11 |
| NPM1 |  | -5.295 | 8.97E-11 |
| RUNX2 |  | -0.91 | 9.85E-11 |
| ID2 | 2.063 | 0.053 | 1.38E-10 |
| CDK4 |  | 0.059 | 1.39E-10 |
| TGFBR1 |  | -0.16 | 1.43E-10 |

**Supplementary Table 6.** Top 100 gene ontology results for the upregulated DEGs in the for the IPF vs. Normal comparison.

| **GO biological process complete** | **Fold Enrichment** | **Raw P-value** | **FDR** |
| --- | --- | --- | --- |
| mucociliary clearance (GO:0120197) | 7.99 | 2.84E-04 | 2.17E-02 |
| axonemal dynein complex assembly (GO:0070286) | 7.08 | 1.32E-13 | 9.03E-11 |
| inner dynein arm assembly (GO:0036159) | 6.99 | 1.94E-06 | 2.76E-04 |
| outer dynein arm assembly (GO:0036158) | 6.75 | 7.23E-08 | 1.47E-05 |
| epithelial cilium movement involved in extracellular fluid movement (GO:0003351) | 6.52 | 1.45E-12 | 7.84E-10 |
| intraciliary retrograde transport (GO:0035721) | 6.52 | 4.58E-05 | 4.43E-03 |
| axoneme assembly (GO:0035082) | 6.44 | 4.29E-26 | 7.48E-23 |
| regulation of mitotic cytokinesis (GO:1902412) | 6.39 | 7.19E-04 | 4.73E-02 |
| kinetochore organization (GO:0051383) | 6.39 | 1.64E-06 | 2.46E-04 |
| epithelial cilium movement involved in determination of left/right asymmetry (GO:0060287) | 6.32 | 1.31E-04 | 1.11E-02 |
| intraciliary transport (GO:0042073) | 6.22 | 6.67E-13 | 3.88E-10 |
| cerebrospinal fluid circulation (GO:0090660) | 6.09 | 6.96E-05 | 6.42E-03 |
| extracellular transport (GO:0006858) | 6.09 | 5.13E-12 | 2.59E-09 |
| intraciliary anterograde transport (GO:0035720) | 5.91 | 3.68E-05 | 3.68E-03 |
| sperm axoneme assembly (GO:0007288) | 5.87 | 1.38E-07 | 2.48E-05 |
| spindle elongation (GO:0051231) | 5.62 | 5.56E-04 | 3.94E-02 |
| mitotic spindle assembly checkpoint signaling (GO:0007094) | 5.6 | 1.08E-07 | 2.07E-05 |
| mitotic spindle checkpoint signaling (GO:0071174) | 5.6 | 1.08E-07 | 2.04E-05 |
| spindle assembly checkpoint signaling (GO:0071173) | 5.6 | 1.08E-07 | 2.02E-05 |
| negative regulation of mitotic metaphase/anaphase transition (GO:0045841) | 5.54 | 5.75E-08 | 1.20E-05 |
| kinetochore assembly (GO:0051382) | 5.48 | 2.88E-04 | 2.19E-02 |
| motile cilium assembly (GO:0044458) | 5.45 | 2.13E-13 | 1.28E-10 |
| spindle checkpoint signaling (GO:0031577) | 5.42 | 1.56E-07 | 2.78E-05 |
| microtubule bundle formation (GO:0001578) | 5.36 | 4.57E-25 | 7.17E-22 |
| negative regulation of mitotic sister chromatid separation (GO:2000816) | 5.33 | 4.40E-08 | 9.85E-06 |
| negative regulation of mitotic sister chromatid segregation (GO:0033048) | 5.33 | 4.40E-08 | 9.71E-06 |
| negative regulation of sister chromatid segregation (GO:0033046) | 5.33 | 4.40E-08 | 9.57E-06 |
| negative regulation of metaphase/anaphase transition of cell cycle (GO:1902100) | 5.22 | 1.18E-07 | 2.18E-05 |
| protein localization to chromosome, centromeric region (GO:0071459) | 5.12 | 1.11E-05 | 1.28E-03 |
| sperm flagellum assembly (GO:0120316) | 5.07 | 1.67E-07 | 2.91E-05 |
| negative regulation of chromosome segregation (GO:0051985) | 5.05 | 8.81E-08 | 1.73E-05 |
| negative regulation of chromosome separation (GO:1905819) | 5.05 | 8.81E-08 | 1.70E-05 |
| negative regulation of mitotic nuclear division (GO:0045839) | 4.9 | 6.48E-08 | 1.34E-05 |
| regulation of cilium beat frequency (GO:0003356) | 4.84 | 5.73E-04 | 4.03E-02 |
| centromere complex assembly (GO:0034508) | 4.74 | 2.14E-05 | 2.25E-03 |
| negative regulation of nuclear division (GO:0051784) | 4.38 | 1.60E-07 | 2.81E-05 |
| cilium movement (GO:0003341) | 4.3 | 2.77E-22 | 3.35E-19 |
| cilium organization (GO:0044782) | 4.25 | 2.48E-44 | 3.89E-40 |
| positive regulation of chromosome separation (GO:1905820) | 4.24 | 1.01E-04 | 8.93E-03 |
| attachment of spindle microtubules to kinetochore (GO:0008608) | 4.22 | 1.94E-04 | 1.55E-02 |
| cilium assembly (GO:0060271) | 4.14 | 2.50E-38 | 1.31E-34 |
| regulation of cilium movement (GO:0003352) | 4.04 | 4.77E-06 | 5.75E-04 |
| cilium-dependent cell motility (GO:0060285) | 3.96 | 1.53E-15 | 1.50E-12 |
| cilium or flagellum-dependent cell motility (GO:0001539) | 3.96 | 1.53E-15 | 1.41E-12 |
| regulation of mitotic sister chromatid segregation (GO:0033047) | 3.91 | 4.34E-07 | 6.87E-05 |
| cilium movement involved in cell motility (GO:0060294) | 3.64 | 4.50E-12 | 2.35E-09 |
| chromosome condensation (GO:0030261) | 3.61 | 4.62E-05 | 4.45E-03 |
| mitotic spindle organization (GO:0007052) | 3.58 | 1.60E-09 | 4.64E-07 |
| microtubule-based movement (GO:0007018) | 3.49 | 5.15E-32 | 1.35E-28 |
| sperm motility (GO:0097722) | 3.46 | 1.94E-10 | 7.23E-08 |
| flagellated sperm motility (GO:0030317) | 3.46 | 1.94E-10 | 7.06E-08 |
| protein localization to cilium (GO:0061512) | 3.44 | 4.29E-06 | 5.26E-04 |
| plasma membrane bounded cell projection assembly (GO:0120031) | 3.43 | 1.74E-33 | 6.80E-30 |
| mitotic sister chromatid segregation (GO:0000070) | 3.43 | 1.63E-10 | 6.24E-08 |
| mitotic spindle assembly (GO:0090307) | 3.4 | 3.27E-05 | 3.35E-03 |
| non-motile cilium assembly (GO:1905515) | 3.37 | 1.47E-05 | 1.61E-03 |
| cell projection assembly (GO:0030031) | 3.36 | 8.18E-33 | 2.56E-29 |
| microtubule cytoskeleton organization involved in mitosis (GO:1902850) | 3.26 | 3.91E-10 | 1.33E-07 |
| regulation of microtubule-based movement (GO:0060632) | 3.25 | 2.24E-05 | 2.34E-03 |
| microtubule-based transport (GO:0099111) | 3.19 | 1.59E-14 | 1.39E-11 |
| regulation of cilium assembly (GO:1902017) | 3.18 | 1.23E-05 | 1.38E-03 |
| regulation of mitotic nuclear division (GO:0007088) | 3.17 | 6.34E-09 | 1.74E-06 |
| regulation of chromosome separation (GO:1905818) | 3.13 | 3.09E-08 | 7.45E-06 |
| sister chromatid segregation (GO:0000819) | 3.02 | 8.35E-10 | 2.67E-07 |
| determination of bilateral symmetry (GO:0009855) | 2.93 | 1.89E-08 | 4.70E-06 |
| determination of left/right symmetry (GO:0007368) | 2.93 | 4.24E-08 | 9.78E-06 |
| microtubule cytoskeleton organization (GO:0000226) | 2.92 | 3.06E-30 | 6.86E-27 |
| specification of symmetry (GO:0009799) | 2.91 | 2.07E-08 | 5.08E-06 |
| mitotic nuclear division (GO:0140014) | 2.9 | 3.19E-10 | 1.14E-07 |
| microtubule-based process (GO:0007017) | 2.89 | 1.55E-43 | 1.22E-39 |
| regulation of mitotic sister chromatid separation (GO:0010965) | 2.89 | 3.66E-06 | 4.63E-04 |
| regulation of metaphase/anaphase transition of cell cycle (GO:1902099) | 2.88 | 5.50E-06 | 6.48E-04 |
| regulation of mitotic metaphase/anaphase transition (GO:0030071) | 2.88 | 4.91E-06 | 5.83E-04 |
| mitotic metaphase plate congression (GO:0007080) | 2.87 | 6.52E-04 | 4.39E-02 |
| regulation of chromosome segregation (GO:0051983) | 2.86 | 8.62E-08 | 1.71E-05 |
| protein localization to chromosome (GO:0034502) | 2.8 | 1.01E-04 | 8.92E-03 |
| spindle organization (GO:0007051) | 2.8 | 7.25E-09 | 1.96E-06 |
| regulation of nuclear division (GO:0051783) | 2.77 | 5.25E-08 | 1.11E-05 |
| spindle assembly (GO:0051225) | 2.76 | 1.19E-05 | 1.35E-03 |
| metaphase plate congression (GO:0051310) | 2.75 | 5.18E-04 | 3.76E-02 |
| centrosome cycle (GO:0007098) | 2.74 | 2.65E-05 | 2.75E-03 |
| regulation of sister chromatid segregation (GO:0033045) | 2.73 | 4.60E-06 | 5.59E-04 |
| nuclear chromosome segregation (GO:0098813) | 2.69 | 3.59E-11 | 1.52E-08 |
| negative regulation of chromosome organization (GO:2001251) | 2.67 | 9.90E-05 | 8.82E-03 |
| cytoplasmic translation (GO:0002181) | 2.65 | 2.93E-06 | 3.80E-04 |
| chromosome segregation (GO:0007059) | 2.61 | 1.21E-12 | 6.78E-10 |
| microtubule organizing center organization (GO:0031023) | 2.59 | 3.08E-05 | 3.18E-03 |
| smoothened signaling pathway (GO:0007224) | 2.54 | 1.26E-04 | 1.07E-02 |
| chromosome localization (GO:0050000) | 2.53 | 2.43E-04 | 1.90E-02 |
| regulation of cytokinesis (GO:0032465) | 2.53 | 1.47E-04 | 1.20E-02 |
| establishment of chromosome localization (GO:0051303) | 2.52 | 4.71E-04 | 3.45E-02 |
| organelle assembly (GO:0070925) | 2.5 | 8.13E-30 | 1.59E-26 |
| protein-containing complex localization (GO:0031503) | 2.48 | 2.07E-06 | 2.88E-04 |
| meiotic chromosome segregation (GO:0045132) | 2.47 | 1.25E-04 | 1.06E-02 |
| xenobiotic metabolic process (GO:0006805) | 2.42 | 4.54E-05 | 4.42E-03 |
| nuclear division (GO:0000280) | 2.4 | 2.38E-11 | 1.07E-08 |
| transport along microtubule (GO:0010970) | 2.38 | 4.15E-06 | 5.17E-04 |
| meiotic nuclear division (GO:0140013) | 2.38 | 1.69E-06 | 2.48E-04 |
| mitotic cell cycle checkpoint signaling (GO:0007093) | 2.37 | 4.33E-05 | 4.25E-03 |
| nucleosome assembly (GO:0006334) | 2.36 | 4.78E-04 | 3.49E-02 |

**Supplementary Table 7.** Top 100 gene ontology results for the downregulated DEGs in the for the IPF vs. Normal comparison.

| **GO biological process complete** | **Fold Enrichment** | **Raw P-value** | **FDR** |
| --- | --- | --- | --- |
| regulation of Rho-dependent protein serine/threonine kinase activity (GO:2000298) | 8.15 | 1.96E-03 | 3.94E-02 |
| norepinephrine uptake (GO:0051620) | 8.15 | 1.96E-03 | 3.94E-02 |
| norepinephrine transport (GO:0015874) | 8.15 | 1.96E-03 | 3.93E-02 |
| positive regulation of synaptic plasticity (GO:0031915) | 7.61 | 3.15E-04 | 9.35E-03 |
| bone trabecula formation (GO:0060346) | 7.34 | 9.85E-04 | 2.31E-02 |
| positive regulation of extracellular matrix disassembly (GO:0090091) | 6.52 | 1.51E-03 | 3.23E-02 |
| peptidyl-lysine hydroxylation (GO:0017185) | 6.52 | 1.51E-03 | 3.23E-02 |
| skeletal myofibril assembly (GO:0014866) | 6.52 | 1.51E-03 | 3.22E-02 |
| response to macrophage colony-stimulating factor (GO:0036005) | 6.52 | 2.45E-04 | 7.50E-03 |
| positive regulation of podosome assembly (GO:0071803) | 6.22 | 7.41E-04 | 1.84E-02 |
| retina vasculature morphogenesis in camera-type eye (GO:0061299) | 6.22 | 7.41E-04 | 1.84E-02 |
| collagen biosynthetic process (GO:0032964) | 5.87 | 2.23E-03 | 4.36E-02 |
| vascular associated smooth muscle cell development (GO:0097084) | 5.71 | 1.08E-03 | 2.48E-02 |
| postsynaptic cytoskeleton organization (GO:0099188) | 5.27 | 1.53E-03 | 3.24E-02 |
| monocyte differentiation (GO:0030224) | 5.15 | 1.77E-04 | 5.81E-03 |
| retina vasculature development in camera-type eye (GO:0061298) | 5.15 | 1.77E-04 | 5.80E-03 |
| regulation of podosome assembly (GO:0071801) | 4.89 | 2.11E-03 | 4.18E-02 |
| fructose metabolic process (GO:0006000) | 4.89 | 2.11E-03 | 4.17E-02 |
| vitamin D metabolic process (GO:0042359) | 4.89 | 2.11E-03 | 4.17E-02 |
| mesenchymal to epithelial transition (GO:0060231) | 4.89 | 1.02E-03 | 2.38E-02 |
| surfactant homeostasis (GO:0043129) | 4.89 | 1.02E-03 | 2.37E-02 |
| regulation of extracellular matrix disassembly (GO:0010715) | 4.89 | 1.02E-03 | 2.37E-02 |
| neurotransmitter reuptake (GO:0098810) | 4.68 | 1.62E-04 | 5.38E-03 |
| regulation of endothelial cell chemotaxis (GO:2001026) | 4.66 | 3.30E-04 | 9.64E-03 |
| positive regulation of cell migration involved in sprouting angiogenesis (GO:0090050) | 4.66 | 3.30E-04 | 9.62E-03 |
| collagen fibril organization (GO:0030199) | 4.56 | 2.96E-09 | 3.84E-07 |
| cellular response to ATP (GO:0071318) | 4.35 | 1.84E-03 | 3.77E-02 |
| chemical homeostasis within a tissue (GO:0048875) | 4.35 | 1.84E-03 | 3.77E-02 |
| vascular associated smooth muscle cell differentiation (GO:0035886) | 4.35 | 1.84E-03 | 3.76E-02 |
| endodermal cell differentiation (GO:0035987) | 4.22 | 2.49E-06 | 1.57E-04 |
| positive regulation of extracellular matrix organization (GO:1903055) | 4.19 | 1.86E-04 | 6.02E-03 |
| positive regulation of heart rate (GO:0010460) | 4.14 | 3.74E-04 | 1.07E-02 |
| cell adhesion mediated by integrin (GO:0033627) | 4.06 | 1.27E-05 | 6.46E-04 |
| positive regulation of humoral immune response (GO:0002922) | 4 | 1.52E-03 | 3.23E-02 |
| skeletal muscle contraction (GO:0003009) | 3.91 | 3.09E-04 | 9.26E-03 |
| neurotransmitter uptake (GO:0001504) | 3.91 | 3.09E-04 | 9.24E-03 |
| regulation of osteoblast proliferation (GO:0033688) | 3.84 | 6.16E-04 | 1.62E-02 |
| regulation of extracellular matrix organization (GO:1903053) | 3.84 | 1.46E-06 | 9.97E-05 |
| epiboly involved in wound healing (GO:0090505) | 3.84 | 6.16E-04 | 1.61E-02 |
| wound healing, spreading of cells (GO:0044319) | 3.84 | 6.16E-04 | 1.61E-02 |
| neuromuscular synaptic transmission (GO:0007274) | 3.83 | 1.95E-03 | 3.95E-02 |
| kidney vasculature development (GO:0061440) | 3.83 | 1.95E-03 | 3.94E-02 |
| renal system vasculature development (GO:0061437) | 3.83 | 1.95E-03 | 3.94E-02 |
| endocardial cushion formation (GO:0003272) | 3.83 | 1.95E-03 | 3.93E-02 |
| multicellular organism aging (GO:0010259) | 3.76 | 1.23E-03 | 2.74E-02 |
| steroid catabolic process (GO:0006706) | 3.76 | 1.23E-03 | 2.74E-02 |
| epiboly (GO:0090504) | 3.71 | 7.80E-04 | 1.92E-02 |
| positive regulation of vascular endothelial growth factor production (GO:0010575) | 3.71 | 7.80E-04 | 1.91E-02 |
| cell-substrate junction assembly (GO:0007044) | 3.67 | 1.02E-04 | 3.67E-03 |
| cell surface receptor signaling pathway involved in heart development (GO:0061311) | 3.67 | 2.46E-03 | 4.72E-02 |
| aortic valve morphogenesis (GO:0003180) | 3.67 | 4.95E-04 | 1.35E-02 |
| extracellular matrix assembly (GO:0085029) | 3.63 | 3.14E-04 | 9.35E-03 |
| protein hydroxylation (GO:0018126) | 3.62 | 1.55E-03 | 3.27E-02 |
| positive regulation of macrophage migration (GO:1905523) | 3.62 | 1.55E-03 | 3.27E-02 |
| developmental induction (GO:0031128) | 3.62 | 1.55E-03 | 3.26E-02 |
| semi-lunar valve development (GO:1905314) | 3.58 | 1.27E-04 | 4.43E-03 |
| regulation of calcium ion-dependent exocytosis (GO:0017158) | 3.58 | 1.27E-04 | 4.42E-03 |
| cell-substrate junction organization (GO:0150115) | 3.58 | 1.27E-04 | 4.41E-03 |
| apoptotic cell clearance (GO:0043277) | 3.56 | 8.13E-05 | 3.04E-03 |
| collagen metabolic process (GO:0032963) | 3.53 | 4.51E-06 | 2.60E-04 |
| cytokine production (GO:0001816) | 3.49 | 1.93E-03 | 3.92E-02 |
| mesenchymal cell proliferation (GO:0010463) | 3.49 | 1.93E-03 | 3.91E-02 |
| integrin-mediated signaling pathway (GO:0007229) | 3.49 | 6.15E-09 | 7.36E-07 |
| morphogenesis of an epithelial sheet (GO:0002011) | 3.45 | 4.10E-05 | 1.77E-03 |
| endoderm formation (GO:0001706) | 3.44 | 2.62E-05 | 1.22E-03 |
| regulation of dopamine secretion (GO:0014059) | 3.44 | 4.85E-04 | 1.33E-02 |
| aortic valve development (GO:0003176) | 3.44 | 4.85E-04 | 1.33E-02 |
| musculoskeletal movement (GO:0050881) | 3.41 | 1.95E-04 | 6.25E-03 |
| multicellular organismal movement (GO:0050879) | 3.41 | 1.95E-04 | 6.24E-03 |
| branching involved in ureteric bud morphogenesis (GO:0001658) | 3.4 | 1.24E-04 | 4.33E-03 |
| heart valve morphogenesis (GO:0003179) | 3.38 | 3.22E-05 | 1.44E-03 |
| regulation of astrocyte differentiation (GO:0048710) | 3.37 | 2.38E-03 | 4.61E-02 |
| substrate adhesion-dependent cell spreading (GO:0034446) | 3.37 | 2.06E-05 | 9.88E-04 |
| vasoconstriction (GO:0042310) | 3.35 | 9.44E-04 | 2.23E-02 |
| regulation of cell migration involved in sprouting angiogenesis (GO:0090049) | 3.35 | 5.96E-04 | 1.58E-02 |
| semaphorin-plexin signaling pathway (GO:0071526) | 3.26 | 4.61E-04 | 1.28E-02 |
| positive regulation of axon extension (GO:0045773) | 3.26 | 7.29E-04 | 1.83E-02 |
| heterotypic cell-cell adhesion (GO:0034113) | 3.26 | 1.15E-03 | 2.61E-02 |
| regulation of transforming growth factor beta production (GO:0071634) | 3.26 | 7.29E-04 | 1.83E-02 |
| ureteric bud morphogenesis (GO:0060675) | 3.26 | 4.81E-05 | 2.00E-03 |
| positive regulation of protein tyrosine kinase activity (GO:0061098) | 3.26 | 3.07E-05 | 1.38E-03 |
| metanephric nephron development (GO:0072210) | 3.26 | 1.83E-03 | 3.78E-02 |
| response to ATP (GO:0033198) | 3.26 | 1.83E-03 | 3.78E-02 |
| regulation of axon extension involved in axon guidance (GO:0048841) | 3.26 | 1.83E-03 | 3.77E-02 |
| regulation of vascular endothelial growth factor production (GO:0010574) | 3.26 | 1.83E-03 | 3.77E-02 |
| postsynapse organization (GO:0099173) | 3.22 | 4.36E-07 | 3.47E-05 |
| mesonephric tubule morphogenesis (GO:0072171) | 3.2 | 5.84E-05 | 2.33E-03 |
| platelet aggregation (GO:0070527) | 3.18 | 5.60E-04 | 1.50E-02 |
| positive regulation of blood circulation (GO:1903524) | 3.18 | 8.84E-04 | 2.12E-02 |
| heart valve development (GO:0003170) | 3.16 | 2.88E-05 | 1.32E-03 |
| leukocyte cell-cell adhesion (GO:0007159) | 3.14 | 1.11E-04 | 3.91E-03 |
| positive regulation of signaling receptor activity (GO:2000273) | 3.12 | 4.29E-04 | 1.21E-02 |
| negative regulation of hemostasis (GO:1900047) | 3.12 | 4.29E-04 | 1.20E-02 |
| modulation of excitatory postsynaptic potential (GO:0098815) | 3.11 | 6.76E-04 | 1.73E-02 |
| positive regulation of endothelial cell migration (GO:0010595) | 3.11 | 1.57E-07 | 1.39E-05 |
| smooth muscle contraction (GO:0006939) | 3.1 | 8.49E-05 | 3.15E-03 |
| smooth muscle cell differentiation (GO:0051145) | 3.09 | 1.69E-03 | 3.52E-02 |
| endothelial cell migration (GO:0043542) | 3.08 | 1.71E-05 | 8.36E-04 |
| positive regulation of blood vessel endothelial cell migration (GO:0043536) | 3.08 | 2.09E-04 | 6.59E-03 |
| artery morphogenesis (GO:0048844) | 3.07 | 4.16E-05 | 1.79E-03 |

**Supplementary Table 8.** Top 100 canonical pathway analysis results for the post-COVID fibrosis vs. Normal DEG comparison, as calculated by the Ingenuity Pathway Analysis tool (results sorted by -log(p-value)). A positive z-score indicates that the pathway is predicted to be activated, while a negative z-score signifies predicted inhibition.

| **Post-COVID Fibrosis Common Genes - IPA Canonical Pathway Analysis** | | | |
| --- | --- | --- | --- |
| **Ingenuity Canonical Pathways** | **-log(p-value)** | **Ratio** | **z-score** |
| Role of Hypercytokinemia/hyperchemokinemia in the Pathogenesis of Influenza | 8.56 | 0.291 | -3.4 |
| Interferon Signaling | 5.94 | 0.361 | -2.887 |
| CREB Signaling in Neurons | 4.98 | 0.125 | -0.12 |
| Pathogen Induced Cytokine Storm Signaling Pathway | 4.58 | 0.137 | -0.14 |
| LPS/IL-1 Mediated Inhibition of RXR Function | 4.36 | 0.15 | -1 |
| S100 Family Signaling Pathway | 4.32 | 0.115 | -2.014 |
| G-Protein Coupled Receptor Signaling | 3.98 | 0.115 | -0.333 |
| Phagosome Formation | 3.71 | 0.114 | -0.563 |
| Eicosanoid Signaling | 3.7 | 0.214 | 1.342 |
| Breast Cancer Regulation by Stathmin1 | 3.58 | 0.116 | -0.855 |
| Neurovascular Coupling Signaling Pathway | 3.46 | 0.142 | 1.061 |
| Melatonin Degradation I | 3.19 | 0.21 | 0.832 |
| Multiple Sclerosis Signaling Pathway | 3.15 | 0.14 | 0.898 |
| Cell Cycle Control of Chromosomal Replication | 3.07 | 0.214 | -3.464 |
| Coagulation System | 3.03 | 0.257 | -1.667 |
| Macrophage Classical Activation Signaling Pathway | 2.97 | 0.143 | 0.577 |
| Chondroitin Sulfate Biosynthesis (Late Stages) | 2.96 | 0.22 | -0.302 |
| Superpathway of Melatonin Degradation | 2.86 | 0.194 | 0.832 |
| Synaptic Long Term Depression | 2.67 | 0.136 | 0.626 |
| Estrogen-mediated S-phase Entry | 2.59 | 0.269 | -2.646 |
| Regulation of Cellular Mechanics by Calpain Protease | 2.57 | 0.169 | 2.449 |
| Salvage Pathways of Pyrimidine Deoxyribonucleotides | 2.52 | 0.444 | -2 |
| Role Of Osteoblasts In Rheumatoid Arthritis Signaling Pathway | 2.49 | 0.127 | 1.095 |
| Chondroitin Sulfate Biosynthesis | 2.42 | 0.19 | -0.302 |
| Oxytocin In Spinal Neurons Signaling Pathway | 2.4 | 0.229 | 1.414 |
| Dermatan Sulfate Biosynthesis | 2.3 | 0.183 | -0.302 |
| VDR/RXR Activation | 2.25 | 0.167 | -0.632 |
| Gustation Pathway | 2.22 | 0.128 | -2.041 |
| Pyrimidine Deoxyribonucleotides De Novo Biosynthesis I | 2.21 | 0.261 | -2.449 |
| IL-10 Signaling | 2.2 | 0.136 | -1.964 |
| Role of BRCA1 in DNA Damage Response | 2.16 | 0.163 | -1.414 |
| Role of Pattern Recognition Receptors in Recognition of Bacteria and Viruses | 2.14 | 0.135 | -1.897 |
| Dermatan Sulfate Biosynthesis (Late Stages) | 2.1 | 0.191 | -1 |
| Synaptogenesis Signaling Pathway | 2.05 | 0.114 | -0.354 |
| Thyroid Hormone Metabolism II (via Conjugation and/or Degradation) | 2.03 | 0.2 | 0.707 |
| Bupropion Degradation | 2.02 | 0.24 | 0 |
| Neuroinflammation Signaling Pathway | 2 | 0.114 | -0.365 |
| Xenobiotic Metabolism CAR Signaling Pathway | 2 | 0.126 | -0.816 |
| Corticotropin Releasing Hormone Signaling | 1.96 | 0.132 | 0.688 |
| LXR/RXR Activation | 1.94 | 0.138 | 1.5 |
| Factors Promoting Cardiogenesis in Vertebrates | 1.93 | 0.131 | -0.688 |
| Retinoate Biosynthesis I | 1.91 | 0.19 | -0.816 |
| Hepatic Fibrosis Signaling Pathway | 1.87 | 0.106 | -0.493 |
| Regulation Of The Epithelial Mesenchymal Transition In Development Pathway | 1.86 | 0.149 | 0.632 |
| PD-1, PD-L1 cancer immunotherapy pathway | 1.82 | 0.14 | -0.577 |
| White Adipose Tissue Browning Pathway | 1.78 | 0.13 | -0.243 |
| MSP-RON Signaling In Macrophages Pathway | 1.75 | 0.134 | -1.5 |
| Estrogen Biosynthesis | 1.73 | 0.178 | 0 |
| CDX Gastrointestinal Cancer Signaling Pathway | 1.72 | 0.119 | 0.408 |
| Complement System | 1.71 | 0.189 | -1 |
| Kinetochore Metaphase Signaling Pathway | 1.69 | 0.135 | -1.155 |
| PXR/RXR Activation | 1.62 | 0.154 | -0.632 |
| Activation of IRF by Cytosolic Pattern Recognition Receptors | 1.62 | 0.154 | -1.265 |
| Nicotine Degradation II | 1.58 | 0.152 | 1.265 |
| Glutamate Receptor Signaling | 1.58 | 0.152 | -0.447 |
| The Visual Cycle | 1.58 | 0.217 | -1.342 |
| Gαs Signaling | 1.57 | 0.128 | 0 |
| Mitotic Roles of Polo-Like Kinase | 1.54 | 0.149 | -2.646 |
| Nicotine Degradation III | 1.53 | 0.155 | 1 |
| Endocannabinoid Neuronal Synapse Pathway | 1.48 | 0.121 | -1.069 |
| Xenobiotic Metabolism PXR Signaling Pathway | 1.47 | 0.115 | -1.279 |
| Phospholipases | 1.46 | 0.145 | 1.134 |
| cAMP-mediated signaling | 1.45 | 0.11 | 0.408 |
| SPINK1 Pancreatic Cancer Pathway | 1.44 | 0.15 | -1 |
| Intrinsic Prothrombin Activation Pathway | 1.43 | 0.167 | 0.378 |
| HOTAIR Regulatory Pathway | 1.4 | 0.117 | -1.698 |
| Serotonin Degradation | 1.39 | 0.141 | 0.632 |
| Acute Phase Response Signaling | 1.38 | 0.114 | 0.832 |
| Neuropathic Pain Signaling In Dorsal Horn Neurons | 1.38 | 0.129 | -0.832 |
| Heparan Sulfate Biosynthesis (Late Stages) | 1.38 | 0.136 | -0.905 |
| Th1 Pathway | 1.37 | 0.123 | 0 |
| Fatty Acid β-oxidation I | 1.34 | 0.171 | 1 |
| Cardiac Hypertrophy Signaling (Enhanced) | 1.33 | 0.0959 | 0 |
| Adrenomedullin signaling pathway | 1.32 | 0.111 | 0.447 |
| Endocannabinoid Cancer Inhibition Pathway | 1.27 | 0.116 | 1.069 |
| Macrophage Alternative Activation Signaling Pathway | 1.26 | 0.109 | 0.426 |
| Cyclins and Cell Cycle Regulation | 1.25 | 0.129 | -1.508 |
| GP6 Signaling Pathway | 1.24 | 0.118 | 0.258 |
| Glutathione-mediated Detoxification | 1.24 | 0.162 | -2.236 |
| Semaphorin Neuronal Repulsive Signaling Pathway | 1.21 | 0.113 | 0 |
| Endothelin-1 Signaling | 1.2 | 0.108 | -0.5 |
| Salvage Pathways of Pyrimidine Ribonucleotides | 1.2 | 0.124 | -2.887 |
| GPCR-Mediated Nutrient Sensing in Enteroendocrine Cells | 1.2 | 0.119 | -2.111 |
| Gαi Signaling | 1.19 | 0.114 | 0.302 |
| Wound Healing Signaling Pathway | 1.17 | 0.103 | 0.784 |
| Role Of Chondrocytes In Rheumatoid Arthritis Signaling Pathway | 1.17 | 0.113 | -1 |
| Heparan Sulfate Biosynthesis | 1.16 | 0.125 | -0.905 |
| Role of WNT/GSK-3β Signaling in the Pathogenesis of Influenza | 1.16 | 0.128 | 0 |
| Role of MAPK Signaling in Inhibiting the Pathogenesis of Influenza | 1.13 | 0.127 | 0 |
| Oxytocin In Brain Signaling Pathway | 1.11 | 0.106 | -0.218 |
| Pulmonary Healing Signaling Pathway | 1.11 | 0.106 | 0.655 |
| Sperm Motility | 1.09 | 0.101 | 1.069 |
| Crosstalk between Dendritic Cells and Natural Killer Cells | 1.08 | 0.121 | 0.707 |
| STAT3 Pathway | 1.06 | 0.111 | 0.333 |
| Dopamine Degradation | 1.04 | 0.156 | 0.447 |
| Differential Regulation of Cytokine Production in Intestinal Epithelial Cells by IL-17A and IL-17F | 1.04 | 0.174 | 1 |
| Glioma Invasiveness Signaling | 1 | 0.123 | 0.707 |
| Prolactin Signaling | 0.983 | 0.116 | -0.447 |
| Acetone Degradation I (to Methylglyoxal) | 0.983 | 0.14 | 0 |
| TREM1 Signaling | 0.893 | 0.117 | 1 |

**Supplementary Table 9.** Top 100 upstream regulator results for the post-COVID fibrosis vs. Normal DEG comparison, as calculated by the Ingenuity Pathway Analysis tool (results sorted by p-value of the overlap) A positive z-score indicates that the regulator is predicted to be activated, while a negative z-score signifies predicted inhibition.

| **Post-COVID Fibrosis Common Genes - IPA Upstream Regulator Analysis** | | | |
| --- | --- | --- | --- |
| **Upstream Regulator** | **Expr Log Ratio** | **Activation z-score** | **p-value of overlap** |
| IFNL1 |  | -6.85 | 3.97E-38 |
| STAG2 |  | 5.73 | 5.45E-30 |
| NONO |  | -6.955 | 5.65E-30 |
| Interferon alpha |  | -6.262 | 1.08E-25 |
| TREX1 |  | 5.975 | 2.85E-24 |
| IFNA2 |  | -6.927 | 9.48E-23 |
| IRGM |  | 5.513 | 1.92E-22 |
| RNASEH2B |  | 6.301 | 8.28E-22 |
| PRL |  | -4.997 | 1.49E-21 |
| Irgm1 |  | 6.293 | 1.83E-21 |
| STAT1 | -2.006 | -4.847 | 2.26E-21 |
| MAPK1 |  | 4.101 | 9.53E-21 |
| IFNG |  | -4.378 | 2.05E-20 |
| NKX2-3 |  | 4.752 | 3.58E-20 |
| PGR |  | 2.65 | 2.21E-18 |
| IRF7 | -2.654 | -6.28 | 3.38E-18 |
| CNOT7 |  | 2.621 | 7.68E-18 |
| RNY3 |  | -4.359 | 2.91E-16 |
| TRIM24 |  | 5.032 | 3.38E-16 |
| IFN Beta |  | -4.999 | 3.71E-16 |
| IFNL4 |  | -3.568 | 1.6E-15 |
| FOXC1 |  | -4.148 | 3.63E-15 |
| RC3H1 |  | 4.6 | 1.08E-14 |
| IFNB1 |  | -4.878 | 1.98E-14 |
| TASL |  | -3.479 | 2.71E-14 |
| EIF2AK2 | -1.585 | -4.061 | 1.05E-13 |
| IRF1 |  | -5.241 | 1.8E-13 |
| SLC15A4 |  | -4.357 | 2.15E-13 |
| STAT2 | -1.099 | -3.513 | 2.18E-13 |
| STING1 |  | -4.517 | 2.48E-13 |
| TNF |  | -2.263 | 3.89E-13 |
| IFNA1/IFNA13 | 1.486 | -4.177 | 5.1E-13 |
| IL6 |  | 0.476 | 6.13E-13 |
| IL1RN | -1.267 | 4.845 | 6.59E-13 |
| Ttc39aos1 |  | 4.951 | 8.01E-13 |
| STAT3 |  | 1.987 | 8.59E-13 |
| PNPT1 | -1.432 | 4.583 | 1.05E-12 |
| CEBPB |  | -3.359 | 1.51E-12 |
| Ifnar |  | -4.744 | 1.55E-12 |
| CGAS |  | -3.716 | 1.79E-12 |
| RARA |  | -1.264 | 2.29E-12 |
| IRF3 |  | -5.297 | 2.78E-12 |
| KDM1A |  | -3.621 | 2.81E-12 |
| Eldr |  | -4.762 | 2.94E-12 |
| miR-182-5p (and other miRNAs w/seed UUGGCAA) |  | 3.396 | 9.58E-12 |
| IRF5 |  | -4.536 | 1.2E-11 |
| SENP3 |  | -4.472 | 1.47E-11 |
| IFNAR2 |  | -2.646 | 1.47E-11 |
| IFNAR1 |  | -3.028 | 1.78E-11 |
| ACKR2 |  | 4.472 | 6E-11 |
| CG |  | 0.962 | 6.61E-11 |
| SMARCA4 |  | 0.211 | 8.15E-11 |
| USP8 |  | 4.835 | 8.32E-11 |
| MAVS |  | -4.848 | 1.12E-10 |
| ESR2 |  | 0.519 | 1.55E-10 |
| ZBTB10 |  | -4.275 | 2.01E-10 |
| NGEF | -1.886 | 0.784 | 2.3E-10 |
| FOXM1 | -1.556 | -2.065 | 2.78E-10 |
| TGFB1 |  | -1.4 | 6.12E-10 |
| DDX58 | -2.26 | -4.226 | 7.16E-10 |
| Vegf |  | -2.555 | 9.04E-10 |
| AGT |  | -1.395 | 1.83E-09 |
| TLR3 |  | -0.68 | 1.84E-09 |
| JAK |  | -2 | 2.97E-09 |
| Ifn |  | -3.237 | 3.12E-09 |
| VDR |  | -2.319 | 5.4E-09 |
| TGM2 |  | -2.241 | 5.68E-09 |
| NR1H3 |  | 0.696 | 6.5E-09 |
| mir-96 |  | -4.101 | 6.75E-09 |
| CSF2 | 3.403 | -3.226 | 8.51E-09 |
| Histone h4 |  |  | 1.01E-08 |
| TERT |  | -3.489 | 1.45E-08 |
| CCND1 |  | -3.271 | 1.69E-08 |
| SPI1 |  | -2.8 | 1.79E-08 |
| Immunoglobulin |  | -0.408 | 1.83E-08 |
| PTGER2 |  | -3.283 | 2.01E-08 |
| KAT6A |  | -2.584 | 2.28E-08 |
| E2f |  | -3.711 | 2.31E-08 |
| PML |  | -3.816 | 0.000000029 |
| HDAC1 |  | 1.658 | 2.99E-08 |
| mir-183 |  | -4.231 | 3.36E-08 |
| SP1 |  | -1.375 | 3.72E-08 |
| DUSP1 |  | 1.333 | 4.32E-08 |
| IRF9 | -1.34 | -3.256 | 4.62E-08 |
| RNASEH2A |  |  | 5.93E-08 |
| TBX2 |  | -3.714 | 7.01E-08 |
| IRF8 |  | -0.196 | 7.08E-08 |
| OSM |  | 0.041 | 8.85E-08 |
| CTNNB1 |  | -1.143 | 0.000000103 |
| TCF3 |  | 2.366 | 0.000000115 |
| SOCS1 |  | 2.784 | 0.000000122 |
| LIF |  | 0.145 | 0.000000135 |
| WWTR1 |  | 1.05 | 0.000000144 |
| CLEC12A |  | -2.425 | 0.00000015 |
| CITED2 |  | 3.243 | 0.000000153 |
| CASR | -1.905 | -2.79 | 0.000000161 |
| TLR7 |  | -2.819 | 0.000000162 |
| SOX9 |  | 0.384 | 0.000000179 |
| IFNA14 |  | -2.581 | 0.00000021 |
| G protein alpha i |  | -3.897 | 0.000000271 |

**Supplementary Table 10.** Top 100 gene ontology results for the upregulated DEGs in the for the post-COVID fibrosis vs. Normal comparison.

| **GO biological process complete** | **Fold Enrichment** | **Raw P-value** | **FDR** |
| --- | --- | --- | --- |
| positive regulation of high-density lipoprotein particle clearance (GO:0010983) | 33.86 | 4.40E-04 | 2.85E-02 |
| regulation of antigen processing and presentation of peptide antigen via MHC class II (GO:0002586) | 25.4 | 7.54E-04 | 4.30E-02 |
| regulation of high-density lipoprotein particle clearance (GO:0010982) | 25.4 | 7.54E-04 | 4.28E-02 |
| calcium ion-regulated exocytosis of neurotransmitter (GO:0048791) | 16.93 | 1.26E-06 | 4.72E-04 |
| negative regulation of glial cell apoptotic process (GO:0034351) | 15.05 | 3.90E-04 | 2.63E-02 |
| peptide antigen assembly with MHC class II protein complex (GO:0002503) | 14.82 | 2.54E-06 | 7.80E-04 |
| MHC class II protein complex assembly (GO:0002399) | 14.82 | 2.54E-06 | 7.65E-04 |
| surfactant homeostasis (GO:0043129) | 14.82 | 2.54E-06 | 7.50E-04 |
| prostate gland growth (GO:0060736) | 14.11 | 8.87E-05 | 8.43E-03 |
| hyperosmotic salinity response (GO:0042538) | 13.55 | 5.33E-04 | 3.36E-02 |
| chemical homeostasis within a tissue (GO:0048875) | 13.17 | 4.73E-06 | 1.03E-03 |
| negative regulation of synapse organization (GO:1905809) | 12.31 | 7.11E-04 | 4.16E-02 |
| regulation of glial cell apoptotic process (GO:0034350) | 12.31 | 7.11E-04 | 4.14E-02 |
| peptide antigen assembly with MHC protein complex (GO:0002501) | 11.85 | 8.31E-06 | 1.53E-03 |
| MHC protein complex assembly (GO:0002396) | 11.85 | 8.31E-06 | 1.52E-03 |
| neuron cell-cell adhesion (GO:0007158) | 10.58 | 2.65E-04 | 2.00E-02 |
| nitric oxide mediated signal transduction (GO:0007263) | 9.68 | 9.61E-05 | 9.03E-03 |
| antigen processing and presentation of exogenous peptide antigen via MHC class II (GO:0019886) | 9.03 | 1.00E-05 | 1.74E-03 |
| antigen processing and presentation of peptide or polysaccharide antigen via MHC class II (GO:0002504) | 8.96 | 2.92E-06 | 8.04E-04 |
| antigen processing and presentation of peptide antigen via MHC class II (GO:0002495) | 8.47 | 1.50E-05 | 2.39E-03 |
| regulation of antigen processing and presentation (GO:0002577) | 8.06 | 7.62E-04 | 4.30E-02 |
| regulation of urine volume (GO:0035809) | 7.7 | 9.13E-04 | 4.90E-02 |
| mesenchymal cell proliferation (GO:0010463) | 7.26 | 3.68E-04 | 2.53E-02 |
| antigen processing and presentation of exogenous peptide antigen (GO:0002478) | 6.77 | 6.00E-05 | 6.53E-03 |
| negative regulation of blood coagulation (GO:0030195) | 6.63 | 2.43E-05 | 3.34E-03 |
| long-term memory (GO:0007616) | 6.58 | 2.03E-04 | 1.62E-02 |
| negative regulation of hemostasis (GO:1900047) | 6.48 | 2.82E-05 | 3.74E-03 |
| regulation of cholesterol efflux (GO:0010874) | 6.41 | 2.35E-04 | 1.81E-02 |
| negative regulation of coagulation (GO:0050819) | 6.1 | 4.33E-05 | 4.99E-03 |
| animal organ formation (GO:0048645) | 5.93 | 3.58E-04 | 2.50E-02 |
| hydrogen peroxide metabolic process (GO:0042743) | 5.93 | 3.58E-04 | 2.49E-02 |
| vasodilation (GO:0042311) | 5.64 | 1.85E-04 | 1.50E-02 |
| antigen processing and presentation of exogenous antigen (GO:0019884) | 5.53 | 2.09E-04 | 1.67E-02 |
| secondary metabolic process (GO:0019748) | 5.42 | 2.37E-04 | 1.81E-02 |
| modulation of excitatory postsynaptic potential (GO:0098815) | 5.39 | 5.97E-04 | 3.62E-02 |
| regulation of blood coagulation (GO:0030193) | 5.32 | 1.94E-05 | 2.93E-03 |
| calcium-ion regulated exocytosis (GO:0017156) | 5.27 | 6.73E-04 | 3.99E-02 |
| synaptic vesicle exocytosis (GO:0016079) | 5.25 | 1.20E-04 | 1.07E-02 |
| regulation of morphogenesis of a branching structure (GO:0060688) | 5.21 | 3.01E-04 | 2.20E-02 |
| regulation of hemostasis (GO:1900046) | 5.17 | 2.45E-05 | 3.35E-03 |
| immunoglobulin production involved in immunoglobulin-mediated immune response (GO:0002381) | 5.15 | 7.57E-04 | 4.29E-02 |
| negative regulation of smooth muscle cell proliferation (GO:0048662) | 5.11 | 3.38E-04 | 2.41E-02 |
| regulation of systemic arterial blood pressure mediated by a chemical signal (GO:0003044) | 5.04 | 8.49E-04 | 4.70E-02 |
| regulation of neurotransmitter secretion (GO:0046928) | 5.04 | 2.57E-06 | 7.47E-04 |
| extracellular matrix disassembly (GO:0022617) | 5.04 | 8.49E-04 | 4.69E-02 |
| regulation of coagulation (GO:0050818) | 4.97 | 3.44E-05 | 4.25E-03 |
| renal system process (GO:0003014) | 4.96 | 2.71E-07 | 1.47E-04 |
| antigen processing and presentation of peptide antigen (GO:0048002) | 4.92 | 1.89E-04 | 1.53E-02 |
| negative regulation of wound healing (GO:0061045) | 4.91 | 8.51E-05 | 8.24E-03 |
| regulation of synaptic vesicle exocytosis (GO:2000300) | 4.84 | 2.11E-04 | 1.67E-02 |
| vascular transport (GO:0010232) | 4.73 | 2.39E-05 | 3.34E-03 |
| transport across blood-brain barrier (GO:0150104) | 4.73 | 2.39E-05 | 3.31E-03 |
| positive regulation of lipid transport (GO:0032370) | 4.57 | 3.24E-05 | 4.17E-03 |
| acute inflammatory response (GO:0002526) | 4.54 | 7.16E-05 | 7.15E-03 |
| regulation of smooth muscle contraction (GO:0006940) | 4.44 | 7.85E-04 | 4.38E-02 |
| regulation of neurotransmitter transport (GO:0051588) | 4.39 | 1.09E-05 | 1.84E-03 |
| antigen processing and presentation (GO:0019882) | 4.23 | 6.32E-05 | 6.74E-03 |
| kidney epithelium development (GO:0072073) | 4.2 | 2.19E-06 | 7.00E-04 |
| xenobiotic metabolic process (GO:0006805) | 4.05 | 2.46E-05 | 3.33E-03 |
| synaptic vesicle cycle (GO:0099504) | 4.03 | 1.35E-05 | 2.20E-03 |
| blood vessel diameter maintenance (GO:0097746) | 4 | 4.03E-06 | 9.72E-04 |
| regulation of tube diameter (GO:0035296) | 4 | 4.03E-06 | 9.58E-04 |
| negative regulation of response to wounding (GO:1903035) | 3.98 | 3.96E-04 | 2.66E-02 |
| neurotransmitter secretion (GO:0007269) | 3.98 | 3.96E-04 | 2.64E-02 |
| signal release from synapse (GO:0099643) | 3.98 | 3.96E-04 | 2.63E-02 |
| regulation of tube size (GO:0035150) | 3.97 | 4.39E-06 | 9.69E-04 |
| positive regulation of lipid localization (GO:1905954) | 3.97 | 5.83E-05 | 6.40E-03 |
| vascular process in circulatory system (GO:0003018) | 3.93 | 6.80E-10 | 1.18E-06 |
| artery development (GO:0060840) | 3.92 | 2.34E-04 | 1.80E-02 |
| nephron epithelium development (GO:0072009) | 3.83 | 1.49E-04 | 1.27E-02 |
| vesicle-mediated transport in synapse (GO:0099003) | 3.76 | 2.82E-05 | 3.72E-03 |
| regulation of systemic arterial blood pressure (GO:0003073) | 3.72 | 3.50E-04 | 2.48E-02 |
| neutrophil migration (GO:1990266) | 3.68 | 6.99E-04 | 4.12E-02 |
| regulation of lipid transport (GO:0032368) | 3.65 | 3.84E-05 | 4.57E-03 |
| regulation of wound healing (GO:0061041) | 3.65 | 7.05E-05 | 7.23E-03 |
| regulation of regulated secretory pathway (GO:1903305) | 3.64 | 2.26E-05 | 3.25E-03 |
| regulation of synapse assembly (GO:0051963) | 3.62 | 4.40E-04 | 2.86E-02 |
| cellular hormone metabolic process (GO:0034754) | 3.59 | 8.19E-05 | 8.03E-03 |
| cellular response to xenobiotic stimulus (GO:0071466) | 3.52 | 6.18E-06 | 1.24E-03 |
| nephron development (GO:0072006) | 3.51 | 1.02E-04 | 9.53E-03 |
| branching morphogenesis of an epithelial tube (GO:0048754) | 3.46 | 1.18E-04 | 1.06E-02 |
| digestion (GO:0007586) | 3.45 | 6.34E-04 | 3.78E-02 |
| regulation of lipid localization (GO:1905952) | 3.45 | 2.38E-05 | 3.36E-03 |
| neurotransmitter transport (GO:0006836) | 3.44 | 1.26E-04 | 1.11E-02 |
| hormone metabolic process (GO:0042445) | 3.4 | 3.26E-06 | 8.39E-04 |
| leukocyte chemotaxis (GO:0030595) | 3.39 | 8.51E-05 | 8.28E-03 |
| regulation of ossification (GO:0030278) | 3.33 | 4.87E-04 | 3.09E-02 |
| memory (GO:0007613) | 3.33 | 4.87E-04 | 3.08E-02 |
| morphogenesis of a branching structure (GO:0001763) | 3.29 | 4.10E-05 | 4.80E-03 |
| regulation of neurotransmitter levels (GO:0001505) | 3.19 | 8.21E-06 | 1.53E-03 |
| regulation of synapse structure or activity (GO:0050803) | 3.14 | 1.68E-05 | 2.64E-03 |
| regulation of smooth muscle cell proliferation (GO:0048660) | 3.1 | 5.42E-04 | 3.40E-02 |
| morphogenesis of a branching epithelium (GO:0061138) | 3.08 | 2.24E-04 | 1.74E-02 |
| hemostasis (GO:0007599) | 3.04 | 1.58E-04 | 1.32E-02 |
| extracellular matrix organization (GO:0030198) | 3.02 | 2.79E-06 | 7.82E-04 |
| extracellular structure organization (GO:0043062) | 3.01 | 2.96E-06 | 8.01E-04 |
| regulation of blood pressure (GO:0008217) | 3 | 1.18E-04 | 1.06E-02 |
| external encapsulating structure organization (GO:0045229) | 2.99 | 3.33E-06 | 8.42E-04 |
| homophilic cell adhesion via plasma membrane adhesion molecules (GO:0007156) | 2.99 | 3.01E-04 | 2.20E-02 |
| regulation of body fluid levels (GO:0050878) | 2.98 | 9.96E-08 | 6.25E-05 |

**Supplementary Table 11.** Top 100 gene ontology results for the downregulated DEGs in the for the post-COVID fibrosis vs. Normal comparison.

| **GO biological process complete** | **Fold Enrichment** | **Raw P-value** | **FDR** |
| --- | --- | --- | --- |
| rhombomere development (GO:0021546) | 11.95 | 3.22E-04 | 4.03E-02 |
| interleukin-27-mediated signaling pathway (GO:0070106) | 11.95 | 3.22E-04 | 4.00E-02 |
| double-strand break repair via break-induced replication (GO:0000727) | 9.76 | 4.94E-05 | 8.71E-03 |
| mitotic DNA replication (GO:1902969) | 8.36 | 3.28E-04 | 4.06E-02 |
| DNA unwinding involved in DNA replication (GO:0006268) | 6.84 | 3.66E-05 | 6.92E-03 |
| outer dynein arm assembly (GO:0036158) | 6.54 | 4.84E-05 | 8.62E-03 |
| DNA replication initiation (GO:0006270) | 6.34 | 8.94E-06 | 2.42E-03 |
| nuclear DNA replication (GO:0033260) | 6.27 | 6.32E-05 | 1.07E-02 |
| cell cycle DNA replication (GO:0044786) | 6.02 | 8.17E-05 | 1.28E-02 |
| negative regulation of viral genome replication (GO:0045071) | 5.87 | 6.17E-09 | 6.92E-06 |
| axonemal dynein complex assembly (GO:0070286) | 5.85 | 1.19E-06 | 4.66E-04 |
| response to interferon-alpha (GO:0035455) | 5.82 | 2.47E-04 | 3.20E-02 |
| motile cilium assembly (GO:0044458) | 5.4 | 1.99E-08 | 1.73E-05 |
| axoneme assembly (GO:0035082) | 5.28 | 9.42E-12 | 1.85E-08 |
| epithelial cilium movement involved in extracellular fluid movement (GO:0003351) | 4.78 | 3.72E-05 | 6.94E-03 |
| sperm flagellum assembly (GO:0120316) | 4.65 | 1.99E-04 | 2.65E-02 |
| extracellular transport (GO:0006858) | 4.46 | 6.55E-05 | 1.09E-02 |
| negative regulation of viral process (GO:0048525) | 4.4 | 1.14E-08 | 1.19E-05 |
| microtubule bundle formation (GO:0001578) | 4.11 | 8.32E-10 | 1.00E-06 |
| regulation of viral genome replication (GO:0045069) | 4.04 | 5.63E-07 | 2.38E-04 |
| response to type I interferon (GO:0034340) | 3.86 | 2.11E-04 | 2.78E-02 |
| regulation of DNA-templated DNA replication (GO:0090329) | 3.81 | 1.29E-04 | 1.84E-02 |
| cilium movement (GO:0003341) | 3.74 | 1.18E-10 | 1.54E-07 |
| response to vitamin (GO:0033273) | 3.35 | 8.82E-05 | 1.36E-02 |
| negative regulation of innate immune response (GO:0045824) | 3.3 | 1.65E-04 | 2.27E-02 |
| defense response to virus (GO:0051607) | 3.17 | 6.62E-11 | 1.04E-07 |
| defense response to symbiont (GO:0140546) | 3.16 | 7.46E-11 | 1.06E-07 |
| recombinational repair (GO:0000725) | 3.14 | 1.07E-05 | 2.72E-03 |
| double-strand break repair via homologous recombination (GO:0000724) | 3.11 | 1.97E-05 | 4.12E-03 |
| DNA duplex unwinding (GO:0032508) | 3.09 | 1.26E-04 | 1.83E-02 |
| DNA geometric change (GO:0032392) | 3.07 | 8.62E-05 | 1.34E-02 |
| mitotic sister chromatid segregation (GO:0000070) | 3.07 | 1.52E-05 | 3.45E-03 |
| cilium assembly (GO:0060271) | 3.01 | 4.37E-12 | 1.37E-08 |
| DNA-templated DNA replication (GO:0006261) | 3.01 | 5.51E-06 | 1.60E-03 |
| microtubule-based movement (GO:0007018) | 2.95 | 1.37E-13 | 7.14E-10 |
| cilium-dependent cell motility (GO:0060285) | 2.95 | 1.11E-05 | 2.77E-03 |
| cilium or flagellum-dependent cell motility (GO:0001539) | 2.95 | 1.11E-05 | 2.73E-03 |
| regulation of viral life cycle (GO:1903900) | 2.94 | 7.64E-06 | 2.10E-03 |
| cilium organization (GO:0044782) | 2.87 | 6.98E-12 | 1.82E-08 |
| DNA conformation change (GO:0071103) | 2.81 | 2.28E-04 | 2.98E-02 |
| regulation of viral process (GO:0050792) | 2.75 | 9.98E-06 | 2.61E-03 |
| cilium movement involved in cell motility (GO:0060294) | 2.72 | 1.57E-04 | 2.18E-02 |
| regulation of DNA replication (GO:0006275) | 2.71 | 8.08E-05 | 1.28E-02 |
| DNA replication (GO:0006260) | 2.61 | 5.21E-06 | 1.54E-03 |
| sister chromatid segregation (GO:0000819) | 2.6 | 1.43E-04 | 2.02E-02 |
| double-strand break repair (GO:0006302) | 2.55 | 9.97E-06 | 2.65E-03 |
| female gamete generation (GO:0007292) | 2.53 | 1.74E-04 | 2.34E-02 |
| microtubule cytoskeleton organization (GO:0000226) | 2.45 | 7.13E-12 | 1.60E-08 |
| response to virus (GO:0009615) | 2.44 | 4.94E-08 | 3.69E-05 |
| nuclear chromosome segregation (GO:0098813) | 2.43 | 1.13E-05 | 2.72E-03 |
| spermatid differentiation (GO:0048515) | 2.37 | 1.06E-04 | 1.57E-02 |
| spermatid development (GO:0007286) | 2.36 | 1.48E-04 | 2.07E-02 |
| plasma membrane bounded cell projection assembly (GO:0120031) | 2.36 | 1.54E-08 | 1.42E-05 |
| cell projection assembly (GO:0030031) | 2.33 | 1.50E-08 | 1.47E-05 |
| nuclear division (GO:0000280) | 2.33 | 1.02E-06 | 4.21E-04 |
| DNA recombination (GO:0006310) | 2.32 | 3.35E-05 | 6.40E-03 |
| microtubule-based process (GO:0007017) | 2.32 | 9.28E-15 | 7.28E-11 |
| chromosome segregation (GO:0007059) | 2.24 | 1.70E-05 | 3.75E-03 |
| organelle fission (GO:0048285) | 2.2 | 4.09E-06 | 1.28E-03 |
| meiotic cell cycle (GO:0051321) | 2.2 | 7.91E-05 | 1.27E-02 |
| chromosome organization (GO:0051276) | 2.07 | 2.74E-06 | 9.56E-04 |
| positive regulation of cell cycle process (GO:0090068) | 2.07 | 3.74E-04 | 4.54E-02 |
| regulation of microtubule-based process (GO:0032886) | 2.06 | 3.02E-04 | 3.85E-02 |
| germ cell development (GO:0007281) | 2.04 | 9.66E-05 | 1.46E-02 |
| cellular process involved in reproduction in multicellular organism (GO:0022412) | 1.95 | 4.15E-05 | 7.58E-03 |
| organelle assembly (GO:0070925) | 1.92 | 2.07E-08 | 1.71E-05 |
| mitotic cell cycle process (GO:1903047) | 1.88 | 1.19E-05 | 2.79E-03 |
| mitotic cell cycle (GO:0000278) | 1.81 | 1.14E-05 | 2.70E-03 |
| cell cycle process (GO:0022402) | 1.73 | 2.83E-06 | 9.44E-04 |
| cell projection organization (GO:0030030) | 1.73 | 3.18E-08 | 2.49E-05 |
| cell division (GO:0051301) | 1.72 | 2.97E-04 | 3.82E-02 |
| plasma membrane bounded cell projection organization (GO:0120036) | 1.71 | 1.44E-07 | 9.41E-05 |
| regulation of cell cycle process (GO:0010564) | 1.66 | 1.02E-04 | 1.52E-02 |
| DNA metabolic process (GO:0006259) | 1.62 | 8.93E-05 | 1.36E-02 |
| cytoskeleton organization (GO:0007010) | 1.58 | 3.55E-06 | 1.14E-03 |
| gamete generation (GO:0007276) | 1.57 | 3.92E-04 | 4.69E-02 |
| cell cycle (GO:0007049) | 1.57 | 5.06E-06 | 1.53E-03 |
| multicellular organismal reproductive process (GO:0048609) | 1.54 | 3.11E-04 | 3.93E-02 |
| defense response to other organism (GO:0098542) | 1.52 | 1.11E-04 | 1.62E-02 |
| defense response (GO:0006952) | 1.43 | 1.28E-04 | 1.84E-02 |
| response to external stimulus (GO:0009605) | 1.35 | 2.98E-05 | 5.77E-03 |
| primary metabolic process (GO:0044238) | 0.85 | 1.71E-04 | 2.31E-02 |
| organic substance metabolic process (GO:0071704) | 0.84 | 2.38E-05 | 4.84E-03 |
| regulation of metabolic process (GO:0019222) | 0.83 | 6.85E-05 | 1.12E-02 |
| nitrogen compound metabolic process (GO:0006807) | 0.83 | 5.84E-05 | 1.01E-02 |
| metabolic process (GO:0008152) | 0.83 | 2.67E-06 | 9.51E-04 |
| cellular metabolic process (GO:0044237) | 0.82 | 1.54E-05 | 3.46E-03 |
| regulation of primary metabolic process (GO:0080090) | 0.81 | 3.93E-05 | 7.24E-03 |
| regulation of nitrogen compound metabolic process (GO:0051171) | 0.81 | 5.17E-05 | 9.01E-03 |
| regulation of cellular metabolic process (GO:0031323) | 0.78 | 4.33E-06 | 1.33E-03 |
| protein metabolic process (GO:0019538) | 0.78 | 1.70E-04 | 2.32E-02 |
| regulation of gene expression (GO:0010468) | 0.77 | 1.01E-05 | 2.59E-03 |
| regulation of nucleobase-containing compound metabolic process (GO:0019219) | 0.77 | 6.22E-05 | 1.06E-02 |
| macromolecule metabolic process (GO:0043170) | 0.77 | 2.69E-07 | 1.32E-04 |
| regulation of cellular biosynthetic process (GO:0031326) | 0.71 | 3.18E-07 | 1.47E-04 |
| regulation of biosynthetic process (GO:0009889) | 0.71 | 2.04E-07 | 1.19E-04 |
| regulation of macromolecule biosynthetic process (GO:0010556) | 0.71 | 3.79E-07 | 1.70E-04 |
| regulation of RNA metabolic process (GO:0051252) | 0.69 | 2.24E-07 | 1.25E-04 |
| regulation of RNA biosynthetic process (GO:2001141) | 0.69 | 4.95E-07 | 2.16E-04 |
| regulation of DNA-templated transcription (GO:0006355) | 0.68 | 2.55E-07 | 1.29E-04 |

**Supplementary Table 12.** Top 100 canonical pathway analysis results for the DEGs of the IPF vs. post-COVID fibrosis comparison. A positive z-score indicates that the pathway is predicted to be activated, while a negative z-score signifies predicted inhibition.

| **Ingenuity Canonical Pathways** | **-log(p-value)** | **Ratio** | **z-score** |
| --- | --- | --- | --- |
| Kinetochore Metaphase Signaling Pathway | 11.3 | 0.27 | 2.746 |
| Role of Hypercytokinemia/hyperchemokinemia in the Pathogenesis of Influenza | 5.93 | 0.221 | 3.441 |
| Mitotic Roles of Polo-Like Kinase | 5.56 | 0.239 | 1.941 |
| Pyrimidine Deoxyribonucleotides De Novo Biosynthesis I | 5.28 | 0.391 | 2.333 |
| Role of BRCA1 in DNA Damage Response | 5.13 | 0.212 | 2.496 |
| Regulation of Cellular Mechanics by Calpain Protease | 3.93 | 0.18 | -2.828 |
| Estrogen-mediated S-phase Entry | 3.88 | 0.308 | 2.121 |
| Cell Cycle: G2/M DNA Damage Checkpoint Regulation | 3.67 | 0.22 | -2.714 |
| Coagulation System | 3.64 | 0.257 | 0.333 |
| Interferon Signaling | 3.54 | 0.25 | 2.828 |
| Salvage Pathways of Pyrimidine Ribonucleotides | 3.48 | 0.165 | 2.5 |
| Glioma Invasiveness Signaling | 3.26 | 0.178 | -1.667 |
| Cyclins and Cell Cycle Regulation | 3.11 | 0.165 | 2.309 |
| Pathogen Induced Cytokine Storm Signaling Pathway | 2.96 | 0.105 | 0.16 |
| Pyrimidine Ribonucleotides Interconversion | 2.68 | 0.211 | 2.121 |
| Wound Healing Signaling Pathway | 2.63 | 0.111 | -1.512 |
| Pulmonary Fibrosis Idiopathic Signaling Pathway | 2.6 | 0.104 | -2.611 |
| Actin Cytoskeleton Signaling | 2.53 | 0.111 | -2.324 |
| Pyrimidine Ribonucleotides De Novo Biosynthesis | 2.46 | 0.195 | 2.121 |
| Chondroitin Sulfate Biosynthesis (Late Stages) | 2.45 | 0.18 | -2.333 |
| NAD Signaling Pathway | 2.19 | 0.119 | 1.414 |
| Integrin Signaling | 2.14 | 0.108 | -2.065 |
| Cell Cycle Control of Chromosomal Replication | 2.12 | 0.161 | 3 |
| Dermatan Sulfate Biosynthesis (Late Stages) | 2.09 | 0.17 | -2.121 |
| HMGB1 Signaling | 2.07 | 0.114 | -1.265 |
| Role of CHK Proteins in Cell Cycle Checkpoint Control | 2.02 | 0.155 | -0.816 |
| Chondroitin Sulfate Biosynthesis | 2.02 | 0.155 | -2.333 |
| Glioblastoma Multiforme Signaling | 1.96 | 0.111 | -1.155 |
| Dermatan Sulfate Biosynthesis | 1.92 | 0.15 | -2.333 |
| Multiple Sclerosis Signaling Pathway | 1.91 | 0.104 | -0.209 |
| Breast Cancer Regulation by Stathmin1 | 1.85 | 0.0859 | -1.98 |
| Sphingosine-1-phosphate Signaling | 1.74 | 0.117 | -1.667 |
| Role of Pattern Recognition Receptors in Recognition of Bacteria and Viruses | 1.74 | 0.109 | 2.828 |
| Activation of IRF by Cytosolic Pattern Recognition Receptors | 1.71 | 0.138 | 1.667 |
| Pyridoxal 5'-phosphate Salvage Pathway | 1.71 | 0.138 | 1 |
| Ovarian Cancer Signaling | 1.69 | 0.108 | -0.447 |
| STAT3 Pathway | 1.66 | 0.111 | -1.155 |
| ATM Signaling | 1.65 | 0.12 | 0.302 |
| DNA damage-induced 14-3-3σ Signaling | 1.63 | 0.152 | -1.134 |
| Cell Cycle: G1/S Checkpoint Regulation | 1.6 | 0.132 | -0.707 |
| Pancreatic Adenocarcinoma Signaling | 1.58 | 0.111 | -0.707 |
| GP6 Signaling Pathway | 1.55 | 0.11 | -1.732 |
| Actin Nucleation by ARP-WASP Complex | 1.51 | 0.118 | -2 |
| Coronavirus Pathogenesis Pathway | 1.5 | 0.098 | -3.13 |
| Inhibition of Matrix Metalloproteases | 1.48 | 0.154 | -1 |
| Ribonucleotide Reductase Signaling Pathway | 1.42 | 0.1 | 1.698 |
| G-Protein Coupled Receptor Signaling | 1.42 | 0.0797 | -1.336 |
| Hepatic Fibrosis Signaling Pathway | 1.41 | 0.0851 | -3.651 |
| Synaptogenesis Signaling Pathway | 1.4 | 0.0889 | -1.961 |
| Aryl Hydrocarbon Receptor Signaling | 1.38 | 0.101 | 0 |
| Pulmonary Healing Signaling Pathway | 1.35 | 0.0955 | -0.688 |
| Signaling by Rho Family GTPases | 1.32 | 0.0899 | -1.886 |
| ID1 Signaling Pathway | 1.32 | 0.0945 | -1.606 |
| LPS/IL-1 Mediated Inhibition of RXR Function | 1.31 | 0.0906 | 0 |
| Xenobiotic Metabolism CAR Signaling Pathway | 1.26 | 0.0942 | -0.943 |
| Colorectal Cancer Metastasis Signaling | 1.26 | 0.0886 | 0 |
| Sumoylation Pathway | 1.24 | 0.107 | 1.633 |
| Role of JAK family kinases in IL-6-type Cytokine Signaling | 1.24 | 0.114 | 0.333 |
| Role of MAPK Signaling in Inhibiting the Pathogenesis of Influenza | 1.24 | 0.114 | 1.414 |
| Bladder Cancer Signaling | 1.23 | 0.103 | 0.447 |
| Phagosome Formation | 1.22 | 0.0777 | -1.511 |
| RHOGDI Signaling | 1.22 | 0.0909 | 2.496 |
| Neuregulin Signaling | 1.2 | 0.103 | 0 |
| Neuroinflammation Signaling Pathway | 1.19 | 0.0852 | 0.816 |
| Heparan Sulfate Biosynthesis (Late Stages) | 1.18 | 0.111 | -1.667 |
| Cholecystokinin/Gastrin-mediated Signaling | 1.16 | 0.101 | -0.707 |
| Gα12/13 Signaling | 1.13 | 0.0977 | 0.577 |
| CREB Signaling in Neurons | 1.11 | 0.0774 | -1.769 |
| GADD45 Signaling | 1.1 | 0.117 | 0.378 |
| Basal Cell Carcinoma Signaling | 1.1 | 0.111 | -0.447 |
| Macrophage Classical Activation Signaling Pathway | 1.07 | 0.0899 | -0.243 |
| UVA-Induced MAPK Signaling | 1.06 | 0.102 | 0 |
| Semaphorin Neuronal Repulsive Signaling Pathway | 1.06 | 0.0933 | -1.069 |
| Regulation Of The Epithelial Mesenchymal Transition In Development Pathway | 1.03 | 0.103 | -0.707 |
| Heparan Sulfate Biosynthesis | 1 | 0.102 | -1.667 |
| HIF1α Signaling | 0.987 | 0.0865 | -3.153 |
| Glutathione Redox Reactions I | 0.979 | 0.138 | 0 |
| Regulation of Actin-based Motility by Rho | 0.975 | 0.0957 | -2.449 |
| PXR/RXR Activation | 0.955 | 0.108 | 1.134 |
| NER (Nucleotide Excision Repair, Enhanced Pathway) | 0.936 | 0.0962 | 0 |
| Nicotine Degradation II | 0.928 | 0.106 | -0.378 |
| Apelin Adipocyte Signaling Pathway | 0.914 | 0.0978 | -0.378 |
| MSP-RON Signaling In Macrophages Pathway | 0.9 | 0.0924 | -0.302 |
| Pyroptosis Signaling Pathway | 0.893 | 0.0968 | 1.667 |
| Ephrin Receptor Signaling | 0.879 | 0.0842 | -1.89 |
| Agrin Interactions at Neuromuscular Junction | 0.854 | 0.101 | 0 |
| Prolactin Signaling | 0.854 | 0.0947 | 0 |
| Th1 Pathway | 0.845 | 0.0902 | 1.134 |
| Role of PKR in Interferon Induction and Antiviral Response | 0.839 | 0.0882 | 1.732 |
| Xenobiotic Metabolism PXR Signaling Pathway | 0.827 | 0.0833 | -1 |
| Regulation Of The Epithelial Mesenchymal Transition By Growth Factors Pathway | 0.827 | 0.0833 | -1.732 |
| Tumor Microenvironment Pathway | 0.812 | 0.0838 | -2.324 |
| Reelin Signaling in Neurons | 0.807 | 0.087 | -1.265 |
| p53 Signaling | 0.796 | 0.0918 | -1.89 |
| Glioma Signaling | 0.796 | 0.088 | -2 |
| Role Of Osteoclasts In Rheumatoid Arthritis Signaling Pathway | 0.788 | 0.0777 | -2.294 |
| DNA Methylation and Transcriptional Repression Signaling | 0.785 | 0.0893 | 1.89 |
| IL-8 Signaling | 0.775 | 0.081 | -2.673 |
| PCP (Planar Cell Polarity) Pathway | 0.764 | 0.1 | 0.816 |
| Endocannabinoid Developing Neuron Pathway | 0.764 | 0.0866 | 0 |

**Supplementary Table 13.** All upstream regulator results for the DEGs of the IPF vs. post-COVID fibrosis comparison, as calculated by the Ingenuity Pathway Analysis tool (results sorted by p-value of the overlap) A positive z-score indicates that the regulator is predicted to be activated, while a negative z-score signifies predicted inhibition.

| **Upstream Regulator** | **Expr Log Ratio** | **Activation z-score** | **p-value of overlap** |
| --- | --- | --- | --- |
| Eldr |  | 7.111 | 7.67E-56 |
| CEBPB |  | 4.752 | 7.15E-32 |
| Irgm1 |  | -6.745 | 2.95E-31 |
| IFNL1 |  | 6.438 | 1.31E-30 |
| E2F4 |  | -1.974 | 1.54E-30 |
| NONO |  | 6.742 | 4.44E-30 |
| CKAP2L | 5.755 | 5.831 | 9.72E-29 |
| STAG2 |  | -5.471 | 1.11E-27 |
| IFNA2 |  | 6.087 | 1.03E-26 |
| IRGM |  | -5.263 | 2.77E-26 |
| ZBTB17 |  |  | 5.45E-26 |
| Interferon alpha |  | 5.482 | 5.85E-26 |
| PRL |  | 4.666 | 2.72E-25 |
| FOXM1 | 3.662 | 3.65 | 3.08E-25 |
| TREX1 | 1.135 | -5.82 | 1.38E-24 |
| CCND1 |  | 2.714 | 1.45E-24 |
| MAPK1 |  | -4.945 | 1.24E-23 |
| NKX2-3 |  | -3.332 | 1.41E-23 |
| CDKN1A |  | -2.793 | 1.85E-23 |
| NUPR1 |  | -7.839 | 1.46E-22 |
| TP53 |  | -6.125 | 1.54E-22 |
| RNASEH2B |  | -5.475 | 3.37E-22 |
| PTGER2 |  | 5.063 | 4.37E-22 |
| RABL6 |  | 5.657 | 6.74E-22 |
| ERBB2 |  | 1.223 | 9.81E-22 |
| PCLAF | 4.127 | 4.686 | 1.29E-21 |
| IFNG |  | 2.461 | 3.79E-21 |
| MYOD1 |  | 2.147 | 2.42E-20 |
| IRF7 | 3.241 | 6.596 | 3.99E-20 |
| E2f |  | 4.207 | 1.03E-19 |
| PGR |  | -3.408 | 1.72E-19 |
| RNY3 |  | 4.472 | 2.18E-19 |
| Vegf |  | 2.821 | 1.03E-18 |
| STAT1 | 2.371 | 4.44 | 1.13E-18 |
| CDK4 |  | 0.6 | 6.35E-18 |
| TGFB1 |  | -5.799 | 6.44E-18 |
| NR1H3 |  | 1.002 | 4.14E-17 |
| IFNB1 |  | 5.25 | 1.77E-16 |
| IL1RN |  | -3.858 | 3.05E-16 |
| E2F3 |  | 4.086 | 6.74E-16 |
| Ttc39aos1 |  | -4.689 | 1.06E-15 |
| HGF |  | 3.759 | 1.46E-15 |
| STAT3 |  | -2.931 | 2.02E-15 |
| CNOT7 |  | -2.219 | 2.24E-15 |
| SLC15A4 |  | 3.068 | 3.27E-15 |
| CDKN2A | -2.748 | -4.948 | 1.28E-14 |
| ESR2 |  | -0.64 | 1.63E-14 |
| LIN9 | 1.354 | 3.795 | 1.86E-14 |
| AREG |  | 2.717 | 2.25E-14 |
| IFNL4 |  | 3.425 | 2.78E-14 |

**Supplementary Table 14.** Top 100 gene ontology results for the upregulated DEGs of the IPF vs. post-COVID fibrosis comparison.

| **GO biological process complete** | **Fold Enrichment** | **Raw P-value** | **FDR** |
| --- | --- | --- | --- |
| spindle assembly involved in female meiosis I (GO:0007057) | 20.63 | 2.74E-04 | 1.89E-02 |
| positive regulation of chromosome condensation (GO:1905821) | 16.5 | 4.75E-04 | 3.03E-02 |
| strand invasion (GO:0042148) | 16.5 | 4.75E-04 | 3.01E-02 |
| interleukin-27-mediated signaling pathway (GO:0070106) | 14.74 | 1.26E-04 | 9.79E-03 |
| intraciliary anterograde transport (GO:0035720) | 14.56 | 2.23E-09 | 4.22E-07 |
| spindle assembly involved in female meiosis (GO:0007056) | 13.75 | 7.62E-04 | 4.61E-02 |
| mitotic spindle midzone assembly (GO:0051256) | 13.13 | 8.98E-06 | 9.45E-04 |
| mitotic spindle elongation (GO:0000022) | 13.13 | 8.98E-06 | 9.39E-04 |
| spindle elongation (GO:0051231) | 12.7 | 2.41E-06 | 2.76E-04 |
| regulation of mitotic cytokinesis (GO:1902412) | 12.38 | 5.16E-05 | 4.31E-03 |
| meiotic spindle assembly (GO:0090306) | 11.46 | 2.95E-04 | 2.00E-02 |
| regulation of ribonuclease activity (GO:0060700) | 11.46 | 2.95E-04 | 1.99E-02 |
| intraciliary transport (GO:0042073) | 11.41 | 7.57E-17 | 4.75E-14 |
| mitotic spindle assembly checkpoint signaling (GO:0007094) | 11.31 | 1.93E-11 | 5.49E-09 |
| mitotic spindle checkpoint signaling (GO:0071174) | 11.31 | 1.93E-11 | 5.40E-09 |
| spindle assembly checkpoint signaling (GO:0071173) | 11.31 | 1.93E-11 | 5.30E-09 |
| negative regulation of mitotic metaphase/anaphase transition (GO:0045841) | 11.25 | 5.26E-12 | 1.72E-09 |
| spindle checkpoint signaling (GO:0031577) | 10.96 | 2.82E-11 | 7.26E-09 |
| negative regulation of mitotic sister chromatid separation (GO:2000816) | 10.89 | 2.10E-12 | 8.44E-10 |
| negative regulation of mitotic sister chromatid segregation (GO:0033048) | 10.89 | 2.10E-12 | 8.23E-10 |
| negative regulation of sister chromatid segregation (GO:0033046) | 10.89 | 2.10E-12 | 8.03E-10 |
| negative regulation of metaphase/anaphase transition of cell cycle (GO:1902100) | 10.61 | 1.12E-11 | 3.36E-09 |
| protein localization to condensed chromosome (GO:1903083) | 10.32 | 2.90E-05 | 2.66E-03 |
| negative regulation of chromosome segregation (GO:0051985) | 10.32 | 4.37E-12 | 1.49E-09 |
| negative regulation of chromosome separation (GO:1905819) | 10.32 | 4.37E-12 | 1.46E-09 |
| kinetochore organization (GO:0051383) | 10.32 | 5.50E-07 | 7.07E-05 |
| spindle midzone assembly (GO:0051255) | 10.32 | 2.90E-05 | 2.65E-03 |
| protein localization to kinetochore (GO:0034501) | 10.32 | 2.90E-05 | 2.63E-03 |
| protein localization to chromosome, centromeric region (GO:0071459) | 9.9 | 5.66E-08 | 8.45E-06 |
| negative regulation of mitotic nuclear division (GO:0045839) | 9.56 | 1.23E-11 | 3.63E-09 |
| cytoplasmic pattern recognition receptor signaling pathway in response to virus (GO:0039528) | 9.38 | 5.96E-04 | 3.72E-02 |
| intraciliary retrograde transport (GO:0035721) | 8.84 | 2.13E-04 | 1.52E-02 |
| negative regulation of nuclear division (GO:0051784) | 8.6 | 1.64E-11 | 4.76E-09 |
| positive regulation of mitotic cell cycle spindle assembly checkpoint (GO:0090267) | 8.6 | 8.12E-04 | 4.82E-02 |
| positive regulation of spindle checkpoint (GO:0090232) | 8.6 | 8.12E-04 | 4.81E-02 |
| mitotic DNA replication (GO:1902969) | 8.6 | 8.12E-04 | 4.79E-02 |
| mitotic chromosome condensation (GO:0007076) | 8.49 | 7.65E-05 | 6.22E-03 |
| kinetochore assembly (GO:0051382) | 8.25 | 2.87E-04 | 1.96E-02 |
| positive regulation of chromosome separation (GO:1905820) | 8.1 | 9.93E-07 | 1.23E-04 |
| regulation of attachment of spindle microtubules to kinetochore (GO:0051988) | 8.02 | 1.02E-04 | 8.04E-03 |
| regulation of mitotic sister chromatid segregation (GO:0033047) | 7.74 | 2.54E-11 | 6.63E-09 |
| meiotic spindle organization (GO:0000212) | 7.74 | 3.79E-04 | 2.50E-02 |
| centromere complex assembly (GO:0034508) | 7.64 | 4.75E-06 | 5.28E-04 |
| positive regulation of chromosome segregation (GO:0051984) | 7.14 | 2.21E-05 | 2.12E-03 |
| attachment of spindle microtubules to kinetochore (GO:0008608) | 7.14 | 2.21E-05 | 2.10E-03 |
| mitotic sister chromatid segregation (GO:0000070) | 6.88 | 7.80E-19 | 5.32E-16 |
| mitotic sister chromatid cohesion (GO:0007064) | 6.88 | 6.32E-04 | 3.92E-02 |
| mitotic spindle assembly (GO:0090307) | 6.88 | 8.51E-09 | 1.47E-06 |
| sperm axoneme assembly (GO:0007288) | 6.63 | 3.59E-05 | 3.16E-03 |
| negative regulation of type I interferon-mediated signaling pathway (GO:0060339) | 6.51 | 7.99E-04 | 4.80E-02 |
| positive regulation of cell cycle checkpoint (GO:1901978) | 6.51 | 7.99E-04 | 4.78E-02 |
| negative regulation of viral genome replication (GO:0045071) | 6.51 | 6.30E-09 | 1.11E-06 |
| negative regulation of double-strand break repair via homologous recombination (GO:2000042) | 6.51 | 7.99E-04 | 4.76E-02 |
| mitotic spindle organization (GO:0007052) | 6.38 | 9.27E-14 | 4.15E-11 |
| response to interferon-alpha (GO:0035455) | 6.28 | 3.53E-04 | 2.35E-02 |
| female meiotic nuclear division (GO:0007143) | 6.07 | 2.52E-05 | 2.38E-03 |
| nuclear DNA replication (GO:0033260) | 6.02 | 4.38E-04 | 2.83E-02 |
| left/right pattern formation (GO:0060972) | 6.02 | 4.38E-04 | 2.82E-02 |
| sister chromatid segregation (GO:0000819) | 5.99 | 2.75E-18 | 1.80E-15 |
| positive regulation of cell cycle G2/M phase transition (GO:1902751) | 5.99 | 6.96E-05 | 5.72E-03 |
| regulation of chromosome separation (GO:1905818) | 5.95 | 7.04E-14 | 3.35E-11 |
| microtubule cytoskeleton organization involved in mitosis (GO:1902850) | 5.92 | 8.97E-16 | 5.41E-13 |
| positive regulation of G2/M transition of mitotic cell cycle (GO:0010971) | 5.89 | 1.93E-04 | 1.40E-02 |
| epithelial cilium movement involved in extracellular fluid movement (GO:0003351) | 5.89 | 5.07E-06 | 5.56E-04 |
| negative regulation of chromosome organization (GO:2001251) | 5.79 | 3.61E-10 | 7.86E-08 |
| cell cycle DNA replication (GO:0044786) | 5.78 | 5.39E-04 | 3.39E-02 |
| chromosome condensation (GO:0030261) | 5.76 | 6.21E-06 | 6.67E-04 |
| mitotic nuclear division (GO:0140014) | 5.72 | 1.44E-19 | 1.07E-16 |
| regulation of chromosome segregation (GO:0051983) | 5.67 | 6.67E-15 | 3.87E-12 |
| response to interferon-beta (GO:0035456) | 5.63 | 1.05E-04 | 8.19E-03 |
| regulation of mitotic nuclear division (GO:0007088) | 5.59 | 2.88E-13 | 1.22E-10 |
| positive regulation of interferon-beta production (GO:0032728) | 5.53 | 2.06E-05 | 2.01E-03 |
| protein localization to chromatin (GO:0071168) | 5.5 | 2.88E-04 | 1.96E-02 |
| extracellular transport (GO:0006858) | 5.5 | 9.20E-06 | 9.55E-04 |
| regulation of mitotic sister chromatid separation (GO:0010965) | 5.47 | 7.30E-11 | 1.79E-08 |
| mitotic G2 DNA damage checkpoint signaling (GO:0007095) | 5.46 | 1.27E-04 | 9.75E-03 |
| axoneme assembly (GO:0035082) | 5.43 | 1.95E-10 | 4.51E-08 |
| regulation of mitotic metaphase/anaphase transition (GO:0030071) | 5.38 | 5.22E-10 | 1.08E-07 |
| ventricular system development (GO:0021591) | 5.32 | 3.47E-04 | 2.32E-02 |
| motile cilium assembly (GO:0044458) | 5.32 | 4.47E-07 | 5.89E-05 |
| protein localization to chromosome (GO:0034502) | 5.23 | 4.87E-08 | 7.35E-06 |
| regulation of metaphase/anaphase transition of cell cycle (GO:1902099) | 5.21 | 9.05E-10 | 1.80E-07 |
| regulation of sister chromatid segregation (GO:0033045) | 5.21 | 8.30E-11 | 1.97E-08 |
| sperm flagellum assembly (GO:0120316) | 5.16 | 1.83E-04 | 1.34E-02 |
| axonemal dynein complex assembly (GO:0070286) | 5.16 | 8.11E-05 | 6.55E-03 |
| nuclear chromosome segregation (GO:0098813) | 5.11 | 2.37E-21 | 1.95E-18 |
| mitotic cytokinesis (GO:0000281) | 5.09 | 3.13E-08 | 5.01E-06 |
| protein localization to cilium (GO:0061512) | 5.07 | 1.71E-06 | 2.03E-04 |
| regulation of cell cycle checkpoint (GO:1901976) | 5.05 | 1.91E-05 | 1.91E-03 |
| centrosome cycle (GO:0007098) | 5.04 | 7.57E-09 | 1.32E-06 |
| mitotic cell cycle checkpoint signaling (GO:0007093) | 5.04 | 6.95E-12 | 2.18E-09 |
| cilium organization (GO:0044782) | 5 | 1.41E-30 | 3.68E-27 |
| negative regulation of viral process (GO:0048525) | 4.99 | 4.06E-09 | 7.48E-07 |
| cell cycle G2/M phase transition (GO:0044839) | 4.99 | 2.03E-06 | 2.36E-04 |
| spindle assembly (GO:0051225) | 4.94 | 4.81E-09 | 8.67E-07 |
| mitotic metaphase plate congression (GO:0007080) | 4.85 | 2.69E-05 | 2.51E-03 |
| cilium assembly (GO:0060271) | 4.81 | 7.40E-26 | 1.16E-22 |
| spindle organization (GO:0007051) | 4.77 | 2.65E-13 | 1.16E-10 |
| establishment of mitotic spindle localization (GO:0040001) | 4.76 | 3.07E-04 | 2.06E-02 |
| regulation of viral genome replication (GO:0045069) | 4.74 | 8.61E-08 | 1.25E-05 |

**Supplementary Table 15.** Top 100 gene ontology results for the downregulated DEGs of the IPF vs. post-COVID fibrosis comparison.

| **GO biological process complete** | **Fold Enrichment** | **Raw P-value** | **FDR** |
| --- | --- | --- | --- |
| negative regulation of antigen processing and presentation of peptide or polysaccharide antigen via MHC class II (GO:0002581) | 40.93 | 2.56E-04 | 1.66E-02 |
| regulation of antigen processing and presentation of peptide or polysaccharide antigen via MHC class II (GO:0002580) | 32.75 | 3.66E-05 | 3.73E-03 |
| regulation of antigen processing and presentation of peptide antigen via MHC class II (GO:0002586) | 30.7 | 4.39E-04 | 2.41E-02 |
| T cell antigen processing and presentation (GO:0002457) | 30.7 | 4.39E-04 | 2.40E-02 |
| regulation of smooth muscle cell-matrix adhesion (GO:2000097) | 30.7 | 4.39E-04 | 2.39E-02 |
| regulation of prostatic bud formation (GO:0060685) | 27.29 | 5.99E-05 | 5.46E-03 |
| negative regulation of plasminogen activation (GO:0010757) | 25.58 | 8.31E-06 | 1.11E-03 |
| response to ozone (GO:0010193) | 20.47 | 1.02E-03 | 4.57E-02 |
| regulation of antigen processing and presentation of peptide antigen (GO:0002583) | 20.47 | 1.02E-03 | 4.56E-02 |
| negative regulation of cholesterol efflux (GO:0090370) | 20.47 | 1.02E-03 | 4.55E-02 |
| positive regulation of astrocyte differentiation (GO:0048711) | 15.74 | 5.01E-05 | 4.70E-03 |
| regulation of glial cell apoptotic process (GO:0034350) | 14.88 | 3.54E-04 | 2.06E-02 |
| regulation of fibrinolysis (GO:0051917) | 13.64 | 1.67E-05 | 1.97E-03 |
| regulation of plasminogen activation (GO:0010755) | 13.64 | 1.67E-05 | 1.95E-03 |
| negative regulation of fibrinolysis (GO:0051918) | 12.59 | 5.94E-04 | 3.01E-02 |
| trans-synaptic signaling, modulating synaptic transmission (GO:0099550) | 12.59 | 5.94E-04 | 3.00E-02 |
| positive regulation of endothelial cell chemotaxis (GO:2001028) | 10.92 | 9.32E-04 | 4.26E-02 |
| negative regulation of fatty acid oxidation (GO:0046322) | 10.92 | 9.32E-04 | 4.25E-02 |
| peptide antigen assembly with MHC class II protein complex (GO:0002503) | 10.23 | 1.14E-03 | 4.98E-02 |
| MHC class II protein complex assembly (GO:0002399) | 10.23 | 1.14E-03 | 4.96E-02 |
| engulfment of apoptotic cell (GO:0043652) | 10.23 | 1.14E-03 | 4.95E-02 |
| surfactant homeostasis (GO:0043129) | 10.23 | 1.14E-03 | 4.94E-02 |
| hyaluronan catabolic process (GO:0030214) | 10.23 | 1.14E-03 | 4.92E-02 |
| regulation of vascular endothelial cell proliferation (GO:1905562) | 9.82 | 7.92E-05 | 6.75E-03 |
| regulation of antigen processing and presentation (GO:0002577) | 9.75 | 3.29E-04 | 2.02E-02 |
| nitric oxide mediated signal transduction (GO:0007263) | 9.75 | 3.29E-04 | 2.01E-02 |
| regulation of endothelial cell chemotaxis (GO:2001026) | 9.75 | 3.29E-04 | 2.00E-02 |
| negative regulation of endothelial cell apoptotic process (GO:2000352) | 9.55 | 2.33E-05 | 2.59E-03 |
| regulation of blood coagulation (GO:0030193) | 9.36 | 1.80E-10 | 1.17E-07 |
| apoptotic cell clearance (GO:0043277) | 9.3 | 5.08E-07 | 1.11E-04 |
| regulation of hemostasis (GO:1900046) | 9.1 | 2.58E-10 | 1.55E-07 |
| regulation of animal organ formation (GO:0003156) | 9.1 | 1.14E-04 | 9.10E-03 |
| positive regulation of hemostasis (GO:1900048) | 8.77 | 1.36E-04 | 1.04E-02 |
| positive regulation of blood coagulation (GO:0030194) | 8.77 | 1.36E-04 | 1.03E-02 |
| regulation of coagulation (GO:0050818) | 8.73 | 4.34E-10 | 2.35E-07 |
| regulation of astrocyte differentiation (GO:0048710) | 8.47 | 1.61E-04 | 1.18E-02 |
| regulation of endothelial cell apoptotic process (GO:2000351) | 8.19 | 1.41E-06 | 2.57E-04 |
| positive regulation of coagulation (GO:0050820) | 8.19 | 1.89E-04 | 1.34E-02 |
| antigen processing and presentation of exogenous peptide antigen via MHC class II (GO:0019886) | 8.19 | 1.89E-04 | 1.34E-02 |
| negative regulation of blood coagulation (GO:0030195) | 8.01 | 5.57E-06 | 8.32E-04 |
| negative regulation of hemostasis (GO:1900047) | 7.84 | 6.50E-06 | 9.10E-04 |
| semaphorin-plexin signaling pathway (GO:0071526) | 7.8 | 2.20E-05 | 2.47E-03 |
| antigen processing and presentation of peptide antigen via MHC class II (GO:0002495) | 7.67 | 2.58E-04 | 1.66E-02 |
| negative regulation of axon extension involved in axon guidance (GO:0048843) | 7.58 | 8.94E-04 | 4.15E-02 |
| negative regulation of coagulation (GO:0050819) | 7.37 | 1.01E-05 | 1.29E-03 |
| epiboly involved in wound healing (GO:0090505) | 7.31 | 1.03E-03 | 4.58E-02 |
| wound healing, spreading of cells (GO:0044319) | 7.31 | 1.03E-03 | 4.56E-02 |
| antigen processing and presentation of peptide or polysaccharide antigen via MHC class II (GO:0002504) | 7.22 | 3.44E-04 | 2.06E-02 |
| establishment of epithelial cell polarity (GO:0090162) | 7.22 | 3.44E-04 | 2.05E-02 |
| cellular response to nutrient (GO:0031670) | 7.22 | 3.44E-04 | 2.04E-02 |
| negative regulation of endothelial cell proliferation (GO:0001937) | 6.66 | 1.73E-04 | 1.24E-02 |
| regulation of cholesterol efflux (GO:0010874) | 6.64 | 5.15E-04 | 2.68E-02 |
| regulation of transforming growth factor beta production (GO:0071634) | 6.3 | 6.61E-04 | 3.28E-02 |
| antigen processing and presentation of exogenous peptide antigen (GO:0002478) | 6.14 | 7.45E-04 | 3.58E-02 |
| negative regulation of lipid localization (GO:1905953) | 6.1 | 2.83E-04 | 1.80E-02 |
| positive regulation of blood vessel endothelial cell migration (GO:0043536) | 6.06 | 1.08E-04 | 8.73E-03 |
| regulation of wound healing (GO:0061041) | 5.98 | 2.93E-09 | 1.21E-06 |
| negative regulation of wound healing (GO:0061045) | 5.93 | 1.80E-05 | 2.07E-03 |
| regulation of collagen metabolic process (GO:0010712) | 5.85 | 9.37E-04 | 4.26E-02 |
| negative regulation of epithelial cell apoptotic process (GO:1904036) | 5.65 | 1.69E-04 | 1.23E-02 |
| morphogenesis of an epithelial sheet (GO:0002011) | 5.62 | 4.42E-04 | 2.39E-02 |
| modulation of excitatory postsynaptic potential (GO:0098815) | 5.58 | 1.17E-03 | 5.00E-02 |
| heterophilic cell-cell adhesion via plasma membrane cell adhesion molecules (GO:0007157) | 5.51 | 4.91E-04 | 2.59E-02 |
| positive regulation of wound healing (GO:0090303) | 5.46 | 2.08E-04 | 1.43E-02 |
| collagen fibril organization (GO:0030199) | 5.46 | 2.08E-04 | 1.42E-02 |
| postsynapse organization (GO:0099173) | 5.4 | 6.35E-06 | 8.97E-04 |
| regulation of endothelial cell proliferation (GO:0001936) | 5.3 | 4.12E-08 | 1.32E-05 |
| positive regulation of response to wounding (GO:1903036) | 5.12 | 1.32E-04 | 1.01E-02 |
| regulation of response to wounding (GO:1903034) | 5.05 | 1.51E-08 | 5.40E-06 |
| regulation of morphogenesis of an epithelium (GO:1905330) | 5.04 | 3.42E-04 | 2.05E-02 |
| positive regulation of endothelial cell migration (GO:0010595) | 4.97 | 5.93E-06 | 8.70E-04 |
| negative regulation of response to wounding (GO:1903035) | 4.82 | 8.97E-05 | 7.56E-03 |
| cortical cytoskeleton organization (GO:0030865) | 4.78 | 1.06E-03 | 4.67E-02 |
| regulation of cell shape (GO:0008360) | 4.75 | 1.83E-07 | 4.49E-05 |
| negative regulation of chemotaxis (GO:0050922) | 4.7 | 1.16E-03 | 4.99E-02 |
| regulation of epithelial cell apoptotic process (GO:1904035) | 4.59 | 5.95E-05 | 5.46E-03 |
| stem cell development (GO:0048864) | 4.23 | 4.80E-04 | 2.56E-02 |
| positive regulation of endothelial cell proliferation (GO:0001938) | 4.22 | 2.42E-04 | 1.61E-02 |
| regulation of axon extension (GO:0030516) | 4.22 | 2.42E-04 | 1.61E-02 |
| negative regulation of lipid metabolic process (GO:0045833) | 4.17 | 1.32E-04 | 1.02E-02 |
| positive regulation of chemotaxis (GO:0050921) | 4.15 | 8.82E-06 | 1.17E-03 |
| regulation of endothelial cell migration (GO:0010594) | 4.12 | 2.47E-06 | 4.08E-04 |
| regulation of postsynapse organization (GO:0099175) | 4.05 | 6.47E-04 | 3.23E-02 |
| regulation of blood vessel endothelial cell migration (GO:0043535) | 4.05 | 6.47E-04 | 3.22E-02 |
| epithelial cell migration (GO:0010631) | 4.01 | 3.51E-04 | 2.07E-02 |
| unsaturated fatty acid metabolic process (GO:0033559) | 3.97 | 3.77E-04 | 2.17E-02 |
| hemostasis (GO:0007599) | 3.91 | 4.69E-06 | 7.15E-04 |
| epithelium migration (GO:0090132) | 3.9 | 4.34E-04 | 2.40E-02 |
| angiogenesis (GO:0001525) | 3.88 | 7.04E-10 | 3.45E-07 |
| antigen processing and presentation (GO:0019882) | 3.84 | 9.21E-04 | 4.25E-02 |
| blood vessel morphogenesis (GO:0048514) | 3.79 | 7.94E-12 | 1.13E-08 |
| synapse assembly (GO:0007416) | 3.79 | 5.32E-04 | 2.75E-02 |
| blood coagulation (GO:0007596) | 3.79 | 1.29E-05 | 1.57E-03 |
| regulation of epithelial cell differentiation (GO:0030856) | 3.77 | 4.63E-05 | 4.43E-03 |
| coagulation (GO:0050817) | 3.74 | 1.47E-05 | 1.75E-03 |
| regulation of cell-substrate adhesion (GO:0010810) | 3.74 | 1.31E-06 | 2.48E-04 |
| endothelium development (GO:0003158) | 3.72 | 1.13E-03 | 4.93E-02 |
| vasculature development (GO:0001944) | 3.69 | 6.65E-14 | 2.61E-10 |
| memory (GO:0007613) | 3.69 | 3.51E-04 | 2.06E-02 |
| axon guidance (GO:0007411) | 3.66 | 9.88E-07 | 1.94E-04 |
